# A foundation model for brain organoid network dynamics uncovers intrinsic computation and convergent disease phenotypes

**DOI:** 10.64898/2026.10.11.758231

**Authors:** Hunter E. Schweiger, Ash Robbins, Sebastian Hernandez, Avelina Moreno-Ochando, Isabel Cline, Kivilcim Doganyigit, Matilda Mouzaya, Kamran Hussain, Zhiping P. Pang, Jubao Duan, Sofie R. Salama, Alex A. Pollen, Aparna Bhaduri, David Haussler, Mircea Teodorescu, Mohammed A. Mostajo-Radji

**Affiliations:** Genomics Institute, University of California Santa Cruz, Santa Cruz, CA 95064, United States; Department of Molecular, Cell, and Developmental Biology, University of California Santa Cruz, Santa Cruz, CA 95064, United States; Department of Electrical and Computer Engineering, University of California Santa Cruz, Santa Cruz, CA 95064, United States; Department of Biomolecular Engineering, University of California Santa Cruz, Santa Cruz, CA 95064, United States; Department of Applied Mathematics, University of California Santa Cruz, Santa Cruz, CA 95064, United States; Department of Neuroscience and Cell Biology, Child Health Institute of New Jersey, Rutgers Robert Wood Johnson Medical School, New Brunswick, NJ 08901, United States; Center for Psychiatric Genetics, Endeavor Health Research Institute, Evanston, IL 60201, United States; Department of Psychiatry and Behavioral Neuroscience, University of Chicago, Chicago, IL 60637, United States; The Eli and Edythe Broad Center of Regeneration Medicine and Stem Cell Research, University of California San Francisco, San Francisco, CA 94143, United States; Department of Neurology, University of California San Francisco, San Francisco, CA 94143, United States; Weill Institute for Neurosciences, University of California San Francisco, San Francisco, CA 94143, United States; Department of Biological Chemistry, David Geffen School of Medicine, University of California, Los Angeles, Los Angeles, CA 90095, United States

**Keywords:** brain organoids, foundation model, high-density microelectrode arrays, neurodevelopmental disorders, closed-loop stimulation

## Abstract

Brain organoids develop self-organizing neural networks that can be used to model neurodevelopment and disease. Here we introduce PULSE, a foundation model developed using over 900 hours of high-density microelectrode array recordings of *in vitro* and *ex vivo* neural tissue. We show that this single model can address two open questions: how disease-associated variants alter network development, and how naive circuits encode information. PULSE revealed accelerated network maturation in held-out human 16p11.2-deletion organoids. It also revealed a change in network coordination shared with *SHANK3* deficiency. Because organoids develop without sensory input, they let us ask how circuits encode information before any representation is learned. Closed-loop control of a virtual inverted pendulum engaged existing spontaneous activity patterns rather than creating new ones. PULSE provides a common reference frame for neural circuits across genotypes, protocols and tasks.

## INTRODUCTION

Brain organoids have become an important model for studying human brain development, evolution and disease^1–3^. They generate the major cortical cell types, recapitulate developmental stages that are otherwise inaccessible in human tissue, and allow different genotypes to be compared under identical culture conditions. Without sensory input, their networks self-organize and display synchronized population bursts and nested oscillations^4^, develop a modular topology^5^, and produce stereotyped burst sequences^6^. High-density microelectrode arrays (HD-MEAs) can record this activity at single-unit resolution^5–7^. When combined with closed-loop stimulation, organoid networks have been embedded in goal-directed tasks in which performance improves over time^8–10^.

Network activity is conventionally characterized using a few hand-chosen summary statistics, typically variants of firing rate, bursting and synchrony, each defined differently across studies^4,7,11^. Similarly, functional experiments train decoders specific to one task and organism. Because the measurement is redesigned for each study, results are difficult to compare across genotypes, protocols or tasks. In addition, brain-machine interface decoders that capture and integrate spike timing across several timescales, from milliseconds to slower patterns, outperform those that use a single timescale^12^. Both summary statistics and single-timescale decoders therefore discard information, so a negative result cannot distinguish a network that lacks a phenotype from a measure that misses it^13,14^.

The lack of a common measure limits two questions that organoids are well placed to answer. First, neurodevelopmental and psychiatric disorders (NPDs), including autism spectrum disorders, arise from hundreds of risk loci yet converge on recognizable behavioral phenotypes^15–19^. Network phenotypes, some of which reflect altered maturation timing, have been reported for a subset of these genes^13,14,20^, but testing whether the convergence extends to network dynamics requires a measure that can detect a common change across genotypes prepared and recorded independently.

Second, external input may either recruit dynamics a network already generates or write new ones into it, a distinction framed as selection versus instruction^21–30^. *In vivo*, sensory experience shapes circuits from the time they form, so it is difficult to know whether task-evoked activity was already present or created by the task. Organoids develop without sensory input, so the two accounts can be distinguished. Selection predicts that task-evoked activity stays within the patterns spontaneous activity produces, and instruction predicts new ones. The test requires describing both kinds of activity in one coordinate system, which per-task and per-animal decoders cannot provide.

Foundation models offer a way to learn such a coordinate system from data rather than design it for each study. Pretrained on large unlabeled datasets, they have transformed protein structure prediction^31,32^, DNA sequence modeling^33^ and single-cell transcriptomics^34,35^. In neural data, models pretrained on spiking recorded with implanted electrode arrays, in the largest case 2,000 hours from more than 30 monkeys and humans, learn to predict the activity of individual neurons, behavior, or both; this allows adaptation to new animals and brain regions with little additional data^36–40^. Their learned representations also distinguish cell types and brain regions from activity alone, without being trained on these labels^39,41,42^. All of these models, however, were trained on recordings from behaving animals, and none on *in vitro* circuits that develop without sensory input.

Here we introduce PULSE (Population Unified Latent Space Embedding), a foundation model built on the Mamba-2 architecture^43^ and developed using over 900 hours of HD-MEA spontaneous and electrically stimulated activity from *in vitro* cultures and *ex vivo* slices. Because it is trained without biological labels and applied with frozen weights, it provides one set of coordinates for every preparation, genotype and task. Because organoid protocols differ widely between laboratories, we compared two NPD mutations of different genetic architecture in cohorts made with two different protocols, asking whether a shared phenotype survives a change in protocol. We also tested whether open-loop stimulation and closed-loop task control select from the dynamics an organoid network already generates or instruct new ones. PULSE revealed accelerated network maturation in 16p11.2-deletion organoids and a change in network coordination shared with *SHANK3* deficiency. Over days of recording, electrode-specific responses persisted while the population as a whole drifted. Both open-loop stimulation and closed-loop control of a virtual inverted pendulum stayed within the dimensions of spontaneous activity, consistent with selection.

## RESULTS

### A foundation model pretrained across brain organoid preparations

We assembled 3,200 HD-MEA recordings, each with at least 30 active units, totaling 914.3 hours of neural spiking data (training 2,027 recordings, 632.3 hr; holdout 1,173 recordings, 282.0 hr; Figure 1A, Table S1). The dataset covers mouse embryonic stem-cell (mESC)-derived dorsal and ventral forebrain organoids, grown both as 3D organoids and as 2D dissociated cultures; *ex vivo* mouse cortical slices; and human induced pluripotent stem cell (iPSC)-derived organoids from four lines held out from pretraining: an isogenic *SHANK3*^+/^*^−^* line with its parental control (KOLF2.2J background, *n* = 6 control and *n* = 5 *SHANK3*^+/^*^−^* organoids), an unaffected-donor control line (CW20107, *n* = 6 organoids) and a 16p11.2 deletion line (14765-X2, *n* = 6 organoids).

**Figure 1:**
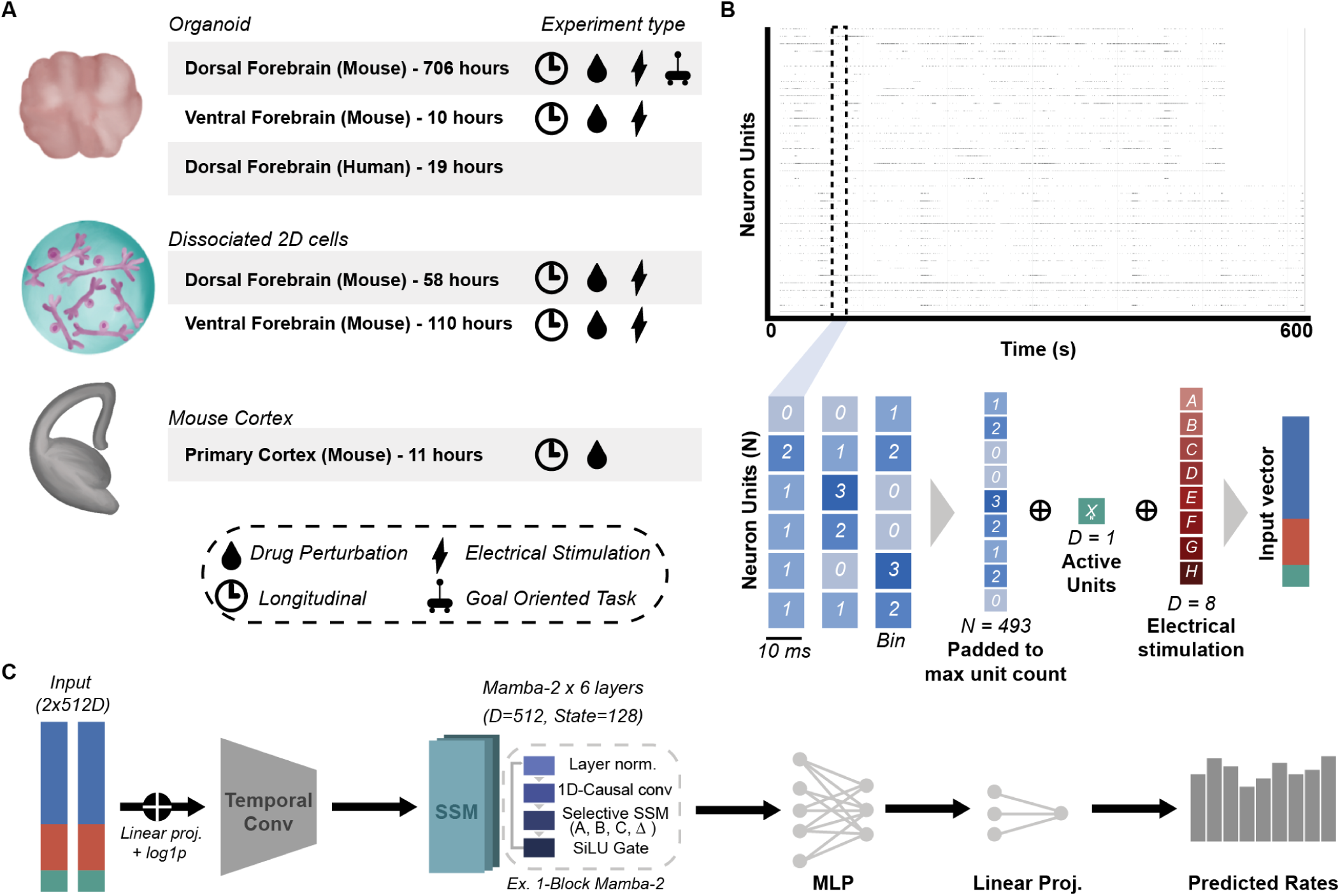
A foundation model learns to predict population spiking across preparations, species and regions. (A) Dataset spanning HD-MEA recordings across preparation types (3D organoid, 2D dissociated, *ex vivo* cortical slice), species (mouse, human) and brain region. Human recordings were held out from pretraining for cross-species evaluation. (B) Diagram of model input showing spike counts binned at 10 ms, concatenated with electrical stimulation features and an active unit count. (C) Model architecture showing the input is linearly projected, temporally downsampled via convolution, and processed by stacked Mamba-2 state-space layers followed by a Poisson prediction head. See also Table S1.

Mamba-2 state-space models^43^ have recently been applied successfully to neural population decoding for *in vivo* behavioral prediction^44,45^. We used Mamba-2 as the backbone of PULSE, which autoregressively predicts the next 20 ms of population firing from past activity. The input is each unit’s spike counts in 10 ms bins, together with the electrical stimulation delivered, the number of active units and a learned embedding of the spike sorter used (Figures 1B and 1C). Input vectors are downsampled by convolution to a 20 ms resolution and processed by six Mamba-2 layers (∼ 12 M parameters). The model compresses this history into a 512-dimensional latent state whose size does not depend on how many units a recording contains, so activity from different experiments can be compared at matched dimensionality. The model was trained on the Poisson likelihood of the observed counts; architecture and training are detailed in STAR Methods.

### PULSE outperforms a forecasting baseline and adapts to unseen organoids with two minutes of data

We split the corpus into training experiments, on which the loss was computed; validation experiments, seen during training but with whole recordings withheld; and holdout experiments from organoids never seen in training (STAR Methods). We excluded human and drug-treated organoids from the Figure 2 cohorts so that model comparisons reflect in-distribution biology, leaving 86 training, 30 of which had multiple recordings to be validation and 9 held-out mouse cultures (Table S2). Pearson *r* between predicted and observed spike counts measures whether PULSE captures the timing of activity regardless of its scale, and bits/spike, the information each spike carries beyond a constant-rate baseline, measures whether it predicts the magnitude^46,47^ (Figures 2B–2E). Skill is the fraction of a baseline’s error that the model removes (0, baseline; 1, perfect; STAR Methods). We scored each metric two ways: per-neuron, and for the summed activity of all neurons in a recording, because a single neuron’s spike train is noisier than the population^47,48^ (Video S1).

**Figure 2:**
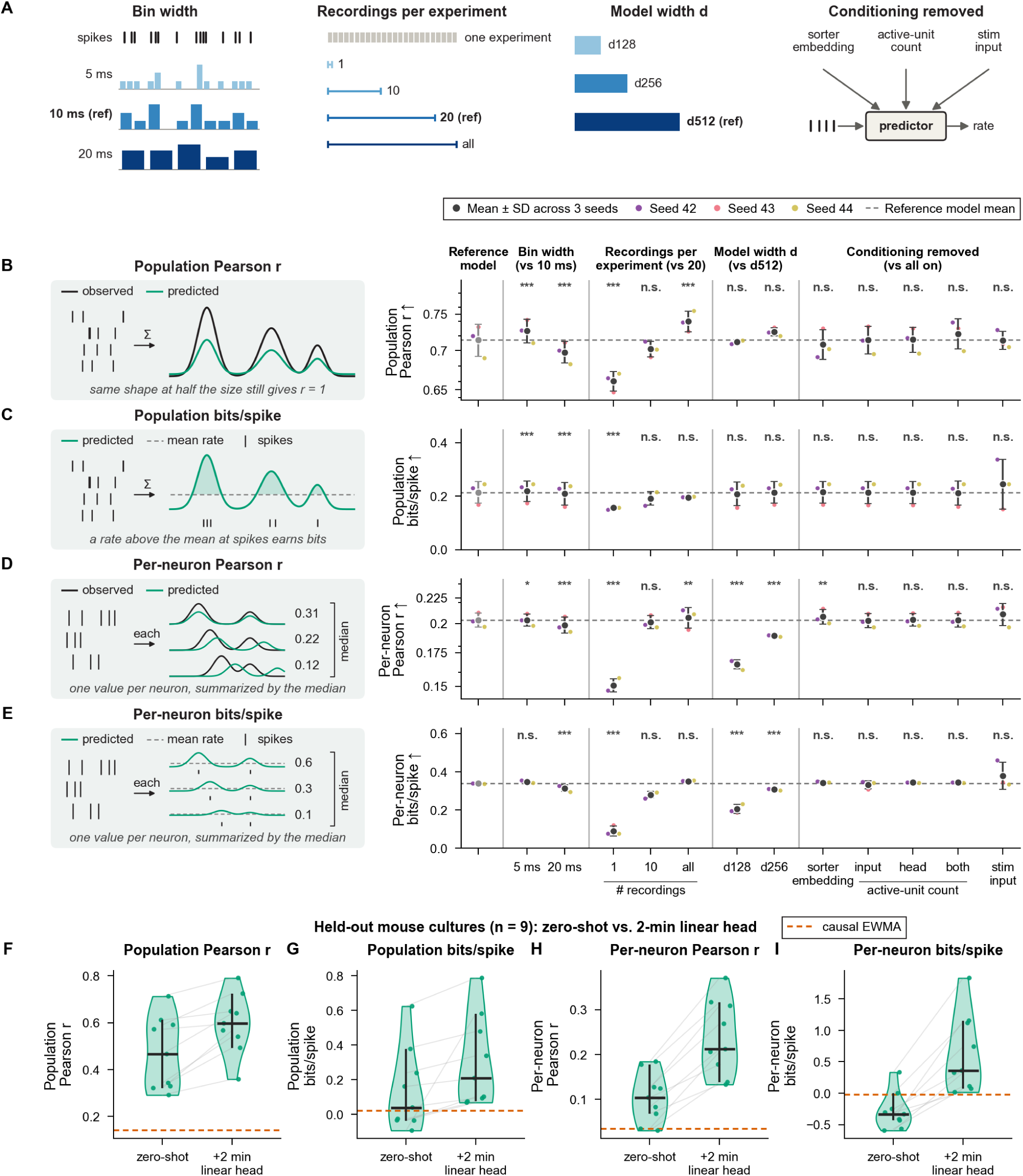
PULSE generalizes to held-out activity, and two minutes of fine-tuning data restore accuracy on held-out organoids. (A) Schematic of the four comparison axes evaluated in (B–E): input bin width, training data per experiment, model width and single-component feature ablations against the reference. (B–E) Population Pearson *r* (B), population bits/spike (C), mean per-neuron Pearson *r* (D) and per-neuron median bits/spike (E) on validation experiments (*n* = 30 organoids), each with a schematic of the metric. Each seed’s value is the median across organoids of per-organoid means; dots, mean ± SD across 3 seeds; dashed line, reference model. Paired two-sided Wilcoxon signed-rank test against the reference on organoid means of seed-averaged experiment values, Holm-corrected within each axis and metric (stimulation input tested alone, *n* = 28 organoids with stimulation); n.s., *P* 0.05; *^∗^P <* 0.05; *^∗∗^P <* 0.01; *^∗∗∗^P <* 0.001. (F–I) Population Pearson *r* (F), population bits/spike (G), per-neuron Pearson *r* (H) and per-neuron median bits/spike (I) on held-out mouse cultures (*n* = 9; six 2D and three 3D organoids), for the all-recordings model zero-shot and after a 2-minute linear head. Dots, individual cultures (seed-averaged, experiments averaged within each culture) joined across conditions; cross, median across cultures and its 95% bootstrap CI; orange dashes, median causal EWMA. Two-sided Wilcoxon signed-rank test across cultures; statistics in the text. See also Figures S1–S3 and Tables S2–S8.

To identify which design choices affect performance, we changed one variable at a time from a reference configuration (*d*_model_ = 512, 10 ms bins, at most 20 recordings per experiment and all conditioning inputs; Figure 2A). As a baseline we used a causal exponentially weighted moving average (EWMA) of each neuron’s own recent firing, a standard forecasting reference^49,50^, with the best-fitting of seven smoothing factors (STAR Methods; Table S3). The reference model exceeded the EWMA on all six metrics in training (*n* = 86 organoids) and validation experiments (validation per-neuron median bits/spike 0.339 against 0.042; all *P* ≤ 6.9 × 10*^−^*^6^; Table S2). On held-out cultures it still exceeded the EWMA on correlation (population and per-neuron Pearson *r*, *P* ≤ 0.008) but not on bits/spike or skill, so we compared design choices on validation organoids (*n* = 30; paired two-sided Wilcoxon signed-rank test across organoids, Holm-corrected within each axis; Table S4) and return to held-out experiments below.

Spike bin width changes both behavioral decoding and inferred functional connectivity *in vivo* ^5,12,51–54^. We trained models on 5, 10 and 20 ms bins, each downsampled to a common 20 ms output. On validation organoids, 10 ms bins outperformed 20 ms bins on every population and per-neuron correlation, bits/spike, skill and Poisson NLL metric (population Pearson *r* 0.723 against 0.697; all *P*_Holm_ ≤ 3.6 × 10*^−^*^5^), but not on per-neuron MAE (*P*_Holm_ = 0.27). Moving to 5 ms raised population Pearson *r* slightly further (0.738, *P*_Holm_ = 6.3 × 10*^−^*^5^) but gave small or non-significant per-neuron gains (Tables S5 and S4; Figures 2B–2E, S1 and S2), so we used 10 ms bins, which halve the input length relative to 5 ms bins.

Scaling recording depth per experiment (1, 10, 20 or all recordings; 266 to 1,756 recordings), rather than the number of experiments, raised validation population Pearson *r* from 0.638 to 0.733 and per-neuron median bits/spike from 0.053 to 0.349, with most of the gain reached by 20 recordings per experiment (Table S6). Widening the model from *d*_model_ = 128 to 512 improved all per-neuron metrics (median bits/spike 0.186 to 0.339; all *P*_Holm_ *<* 3 × 10*^−^*^4^) but not the population metrics (all *P*_Holm_ ≥ 0.18); *d*_model_ = 256 was indistinguishable from 512 on the population metrics (all *P*_Holm_ ≥ 0.065) (Table S7; Figures 2B–2E, S1 and S2).

We then ablated the conditioning inputs one at a time (Tables S8 and S9). Removing the learned spike-sorting source embedding left population bits/spike unchanged on training experiments (+0.00008, *P*_Holm_ = 0.90) and slightly raised per-neuron Pearson *r* on both training experiments (+0.00518, *P*_Holm_ = 3.7 × 10*^−^*^12^) and validation organoids (*P*_Holm_ = 0.001), so sorter identity does not improve accuracy. Removing the active-unit-count channel from the input, the prediction head or both left validation metrics unchanged, so the model does not depend on an explicit neuron-count signal. Zeroing the electrical-stimulation input channels reduced accuracy on the 36 training organoids with stimulation recordings (population bits/spike, *P* = 7.3 × 10*^−^*^10^) but not detectably on the 28 such validation organoids (smallest *P* = 0.11 across the eight metrics), so the stimulation channels carry information beyond the evoked activity itself, detectably only in training organoids.

Because the frozen model generalized on the scale-invariant metrics but not on calibration, we asked whether calibration could be restored with minimal data and parameters. A linear head fit on two minutes (120 s) of each held-out experiment’s activity raised per-neuron median bits/spike from 0.335 to +0.359, improving all 9 held-out mouse cultures (six 2D monolayers and three 3D organoids; paired two-sided Wilcoxon signed-rank test across cultures, *P* = 0.0039; Figure 2I, with the other three metrics in Figures 2F–2H and 0.5- to 4-minute recovery curves in Figure S3). The fine-tuned model exceeded the causal EWMA on all six metrics in at least 8 of the 9 cultures (all *P ≤* 0.008; Table S2).

### PULSE transfers across species, revealing reduced predictability of 16p11.2-deletion activity

Human organoids carrying defined mutations allow network dynamics to be compared across genotypes. We used human iPSCs from two donor backgrounds carrying distinct NPD-associated mutations: 14765-X2, derived from an individual with a 16p11.2 deletion^55^, compared against CW20107, an unaffected donor; and an isogenic *SHANK3*^+/^*^−^* line engineered from the parental KOLF2.2J *SHANK3*^+/+^ by stop-codon insertion^56^ (Figure 3A). The two pairs were made with different protocols, reflecting the range in use across laboratories: the KOLF2.2J organoids from pre-differentiated neural progenitors, and the CW20107 and 14765-X2 organoids by direct aggregation.

**Figure 3:**
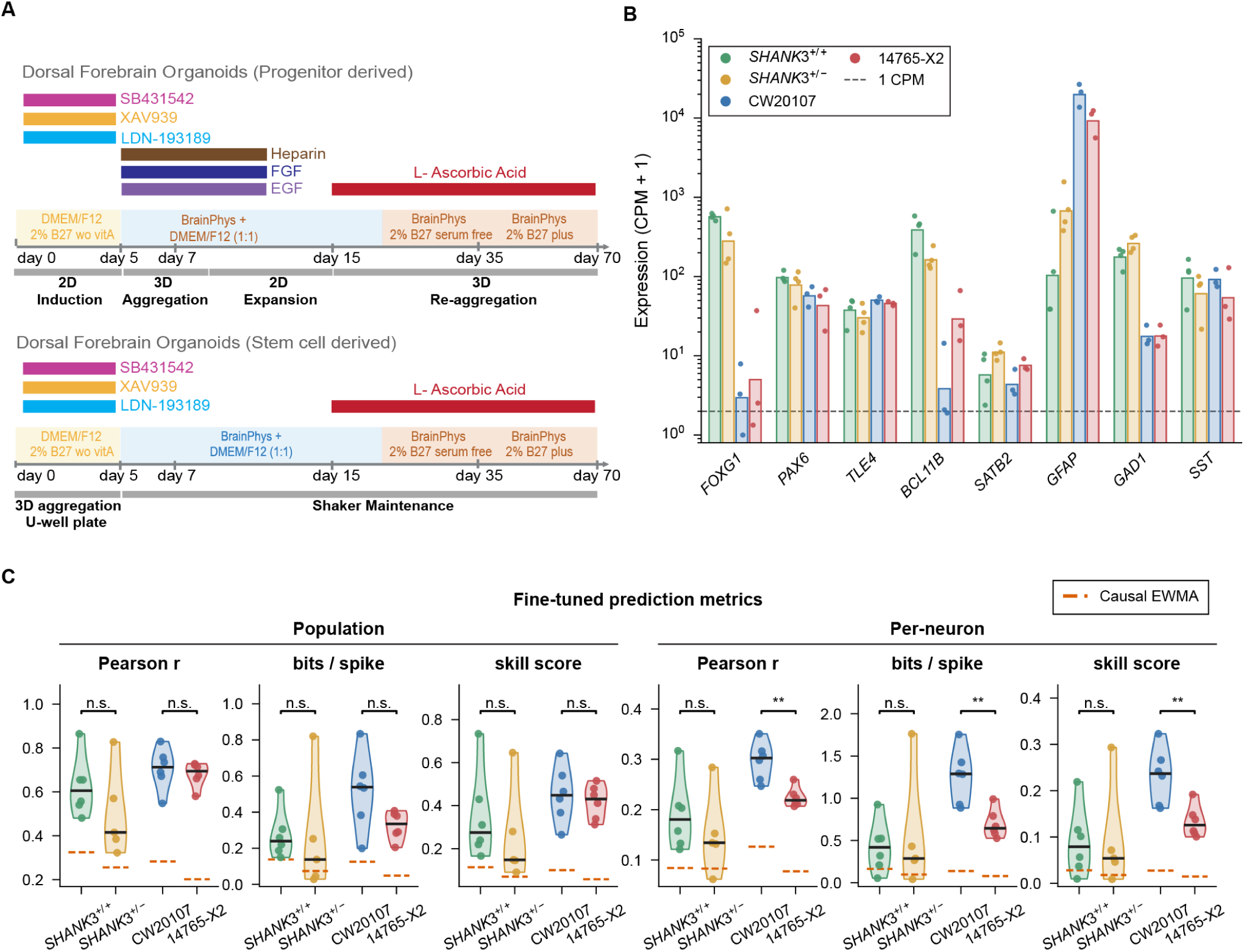
PULSE transfers to human organoids and predicts NPD organoids less accurately than their controls. (A) Differentiation and recording protocols for the KOLF2.2J isogenic pair (*SHANK3*^+/+^ and *SHANK3*^+/^*^−^*) and the CW20107 and 14765-X2 lines. (B) Bulk RNA-seq expression of forebrain, cortical layer, astroglial and GABAergic marker genes in *SHANK3*^+/+^ (*n* = 4), *SHANK3*^+/^*^−^* (*n* = 4), CW20107 (*n* = 3) and 14765-X2 (*n* = 3) organoid samples (each pooled from three organoids; day 180 for CW20107 and 14765-X2, day 100 for KOLF2.2J), plotted as CPM +1 on a log axis. Dots, samples; bars, geometric mean; dashed line, 1 CPM. (C) Prediction accuracy after linear-head fine-tuning on six metrics for the four lines (*n* = 6 *SHANK3*^+/+^, 5 *SHANK3*^+/^*^−^*, 6 CW20107 and 6 14765-X2 organoids). Dots, organoids (mean of their experiments); black bar, median of organoids; orange dashes, median across organoids of each line’s causal EWMA. Brackets, two-sided exact permutation Mann-Whitney *U* test across organoids within a preparation (each organoid summarized by the mean of its experiments); n.s., *P* 0.05; *^∗^P <* 0.05; *^∗∗^P <* 0.01; *^∗∗∗^P <* 0.001. See also Figures S4 and S5 and Tables S10, S11 and S12.

Bulk RNA sequencing of organoids at day 100 post-aggregation (KOLF2.2J) and day 180 (CW20107 and 14765-X2) showed that both preparations express marker genes of forebrain excitatory and GABAergic neurons (Figures 3B and S4; Table S10). Both preparations are cortical but differ in protocol, so each pair provides an independent test of whether the model can read out genotype-dependent differences in network dynamics.

We compared mouse holdout cultures (*n* = 9) with the two human control lines (6 CW20107 and 6 *SHANK3*^+/+^ organoids) on the six metrics of Figure 2 to test whether PULSE’s predictions transfer across species. Zero-shot, CW20107 was predicted better than mouse on the three per-neuron metrics (Cohen’s *d* = 2.0 to 2.7, *P*_Holm_ ≤ 0.0096) but not on the population metrics (*P*_Holm_ ≥ 0.051). After a 2-minute fine-tuned head, neither control line differed from fine-tuned mouse (all *P*_Holm_ ≥ 0.099; Table S11). Once fine-tuned, CW20107 organoids were above the EWMA baseline on every metric in all six organoids, and *SHANK3*^+/+^ organoids on four metrics in all six and on the other two in five of six (Figure 3C; Table S12).

However, the two control lines were not equally predictable. After fine-tuning, CW20107 organoids scored significantly above *SHANK3*^+/+^ on four of six metrics (two of them marginally, *P* = 0.041; e.g. per-neuron bits/spike, organoid medians: 1.289 versus 0.420, Cohen’s *d* = 2.66, *P* = 0.0087, six versus six organoids; Table S11), likely reflecting differences in differentiation protocol or donor background between the two preparations (Figure 3B).

Within each protocol, PULSE predicted 14765-X2 organoids worse than their matched controls, but not *SHANK3*^+/^*^−^* organoids (Figures 3C and S5; Table S12). 14765-X2 organoids were predicted worse on all three per-neuron metrics (per-neuron Pearson *r*, organoid medians: 0.219 versus 0.302, Cohen’s *d* = 2.39, *P* = 0.0043; bits/spike *d* = 2.18, skill *d* = 2.01, both *P* = 0.0087; six versus six organoids), whereas population Pearson *r*, bits/spike and skill were unchanged (*P* ≥ 0.13). No metric differed between *SHANK3*^+/^*^−^* and *SHANK3*^+/+^ organoids (all *P* ≥ 0.24; five versus six organoids). Because neither control line differed from mouse after fine-tuning, the lower accuracy in 14765-X2 organoids reflects a difference between lines within a preparation rather than a failure to transfer to human tissue.

### Two NPD mutations converge on a shared network phenotype

Comparing networks across conditions is difficult because protocols produce distinct developmental timelines^57–60^ and several NPD mutations themselves alter maturation timing^17–20,61–66^. To place organoids from different protocols on one developmental axis, we ran the frozen model over each whole recording and computed the covariance of its latent state across all time points, within the 64 leading principal components of the latent. This covariance is a summary statistic of the latent trajectory, not a parameter of the model, and captures which latent dimensions rise and fall together. We then fit a ridge regression to chronological age on a single control line, CW20107, the line closest to standard protocols in the field^57,67,68^ (*n* = 6 organoids; Figure 4A). Organoids of a line were recorded at the same ages, so we assessed ordering by age within each organoid.

**Figure 4:**
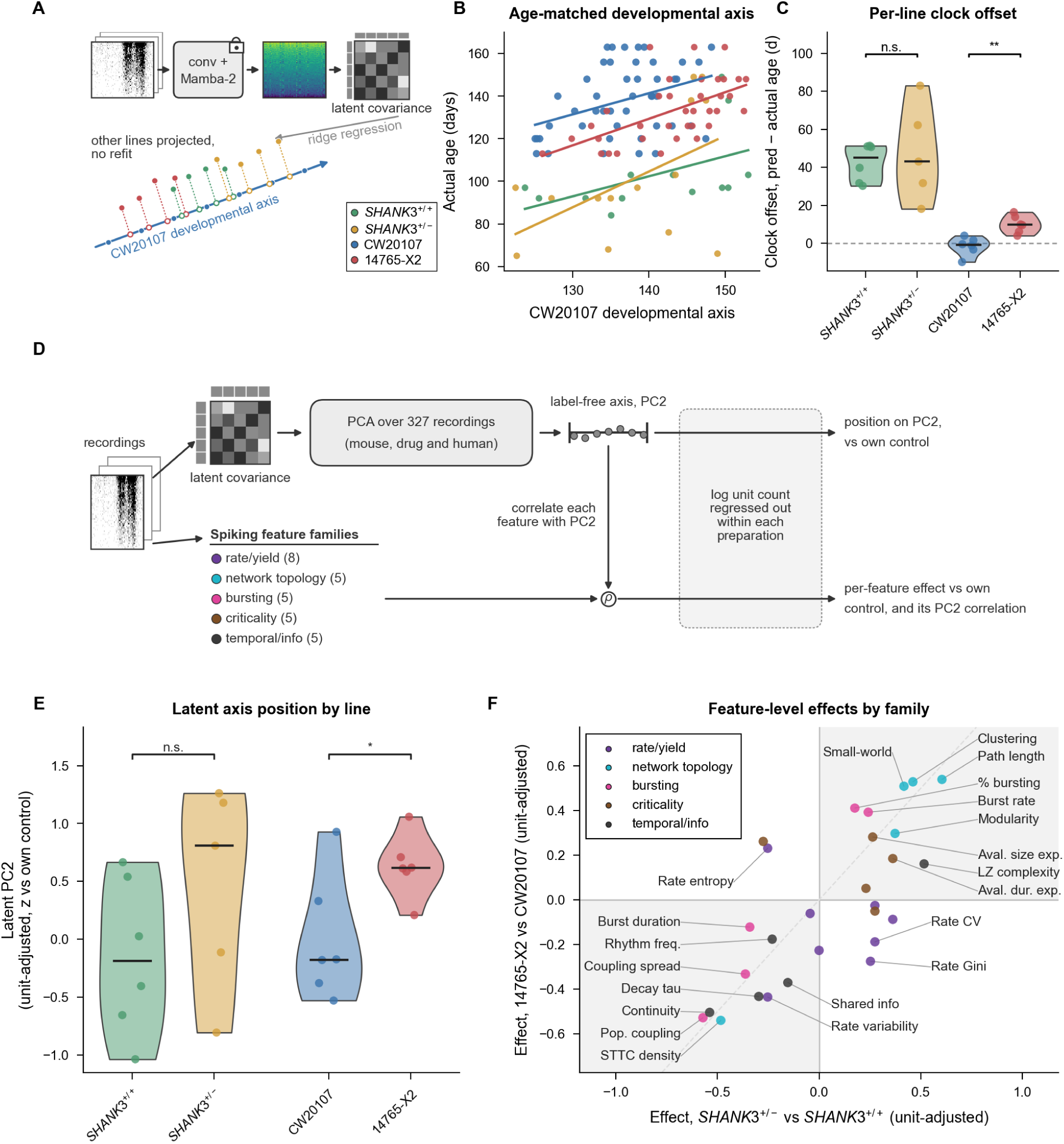
Two NPD mutations converge on network coordination but not firing rate. (A) Schematic of the developmental clock. Ridge regression of latent covariance matrices on chronological age, fit on CW20107 alone and cross-validated leave-one-organoid-out. *SHANK3*^+/+^ (green), *SHANK3*^+/^*^−^* (yellow), CW20107 (blue) and 14765-X2 (red); key applies to (A–C) and (E). (B) Chronological age against position on the CW20107 developmental axis for every recording, so that the ordering of each line’s recordings by age is visible. Dots, individual recordings (*n* = 109 recordings from 23 organoids); lines, least-squares fit per line. (C) Per-line offset from the CW20107-anchored clock. *P* from genotype labels permuted across organoids within each preparation. (D) Schematic. The latent covariance of each spontaneous recording is reduced by PCA to a label-free axis, PC2, and 28 classical spiking features in five families are correlated with it and compared between each NPD line and its own control. (E) Per-organoid position on PC2 (mean of an organoid’s experiments; leave-one-out refit), with log unit count regressed out within each preparation and z-scored against that preparation’s own control; the two controls are pooled for display. Brackets, Cliff’s *δ* over all experiments against the line’s own control, label permutation test across organoids within each preparation. (F) Per-feature effects (Cliff’s *δ*, log unit count regressed out within preparation) of each NPD line against its own control, *SHANK3*^+/^*^−^* versus *SHANK3*^+/+^ (x) and 14765-X2 versus CW20107 (y), colored by family. Each point is one of 28 features; shaded quadrants, same-sign effects; dashed line, identity; labels, selected features. In (C) and (E), *n* = 6 CW20107, 6 14765-X2, 6 *SHANK3*^+/+^ and 5 *SHANK3*^+/^*^−^* organoids; black bar, median of organoids; n.s., *P* 0.05; *^∗^P <* 0.05; *^∗∗^P <* 0.01; *^∗∗∗^P <* 0.001. See also Figure S6 and Table S13.

Performing leave-one-organoid-out, this axis ordered the experiments of each CW20107 organoid by age (Spearman *ρ*: median +0.64, mean +0.46, positive in five of six organoids; permutation *P* = 0.0015, age permuted within organoid), whereas unit count alone did not (mean *ρ* = +0.12, *P* = 0.23). Without refitting, it also ordered the experiments of each 14765-X2 organoid by age (mean *ρ* = +0.70, positive in all six organoids, *P* ≤ 1 × 10*^−^*^4^). The *SHANK3* organoids were mostly recorded once, at different ages, so there we compared organoid-mean predicted age with organoid-mean age: *ρ* = +0.54 in *SHANK3*^+/+^ (*P* = 0.15, six organoids) and *ρ* = −0.10 in *SHANK3*^+/^*^−^* (*P* = 0.61, five organoids), neither significant (Figure 4B). Progenitor-derived *SHANK3* organoids, whose culture age is counted from aggregation, sat at the same axis position as CW20107 despite a younger culture age (median organoid age 96 and 92 versus 137 days; Figure 4B). 14765-X2 experiments, recorded over the same period as CW20107, projected ahead of their chronological age (median organoid offset +9.7 versus 0.8 days; Cliff’s *δ* = +0.449 on experiment offsets, *P* = 0.0022, with all six 14765-X2 organoids ahead of all six CW20107 organoids, the smallest *P* attainable with six organoids per line), whereas *SHANK3*^+/^*^−^* did not differ from *SHANK3*^+/+^ (*P* = 0.95, five versus six organoids; Figure 4C).

We next tested whether an axis defined without labels separates NPD from control organoids. We applied principal component analysis to the latent covariance matrices of all 327 spontaneous recordings (mouse and human, including drug-treated cultures), so that the axis reflects many networks rather than the human cohort alone (Figure 4D). The first two components carried 19.0% and 6.5% of the variance, well above the third (2.6%), and were stable across leave-one-out refits (Figures S6A–S6C). Both components tracked log unit count (organoid means across 23 human organoids: PC1 *ρ* = −0.72, PC2 *ρ* = +0.76), so we regressed log unit count out within each preparation. On the second component, both NPD lines shifted in the same direction; the shift was significant for 14765-X2 and jointly, but not for *SHANK3* alone (14765-X2 Cliff’s *δ* = +0.411, permutation *P* = 0.037, six versus six organoids; *SHANK3*^+/^*^−^ δ* = +0.429, *P* = 0.26, five versus six organoids, joint stratified permutation *P* = 0.022, genotype labels permuted across organoids; Figure 4E; Table S13).

To interpret the second component, we correlated it with 28 classical spiking features in five families (rate and yield, network topology, bursting, criticality and temporal structure; STAR Methods), controlling for log unit count, preparation and genotype (Figure 4D). The component and the two mutations moved the same features in the same direction. Each feature’s partial correlation with the component tracked its mean unit-adjusted effect in the two NPD lines (Spearman *ρ* = +0.464, *P* = 0.013; Figure S6E).

The two mutations also shifted network activity in a common direction relative to their own controls, both in the full latent space (cosine similarity of displacement directions +0.464, permutation *P* = 0.0019; Figure S6D) and across the spiking features (cosine +0.709, *P* = 0.013; Figure 4F), in bursting, network topology and temporal structure (15 of 15 features) but not in rate and yield (2 of 8). Both alignments depended on the unit adjustment (Table S13). Two NPD mutations in independent preparations therefore converge on how network activity is coordinated rather than on how much it fires.

### Ordered population drift spares electrode-specific structure

For a network to compute, its responses to different inputs must be distinct enough for a down-stream readout to decode them^69,70^. In the brain, these responses remain decodable across days even as most of the neurons carrying them change^71–73^, although this stability erodes over weeks and depends on the stimulus^74,75^. *In vivo*, behavior and electrode displacement change alongside intrinsic network dynamics^76–78^, so the structure that survives cannot be assigned to the network itself. Brain organoids, which develop without sensory input, lack these confounds^4,5,7^, so any stable structure in their evoked responses is a property of the network before any representation has been learned; any drift in these responses is largely intrinsic. We asked whether stimulating different sites evokes distinct responses in organoids, and whether those responses persist as the network drifts.

Stimulating one electrode evokes a population response specific to that site. In a fixed-rate sweep (2 Hz, *n* = 17 organoids), a linear discriminant analysis (LDA) decoder identified the stimulated site from PULSE’s latent state at a median 4.14 × chance with the site’s identity hidden from the model (the ID-masked readout), against 1.74 × for the binned firing rate over the same 120 ms window and 2.92 × for its best single 10 ms bin (Figures S7 and S8). The decoder identified the site from the evoked response, not from elapsed time in the recording (Table S14).

We next tested whether the response persists as the stimulation frequency changes and as the network reorganizes over time. We stimulated every electrode at one frequency per recording, stepping through five frequencies from 0.5 to 8 Hz with spontaneous recordings interleaved; one pass through all five frequencies constitutes a cycle (Figure 5A; STAR Methods). We recorded organoids continuously for days as this cycle looped (a median of 49.7 hours of activity per experiment, up to 73.5 hours), tracking the same spike-sorted units throughout.

**Figure 5:**
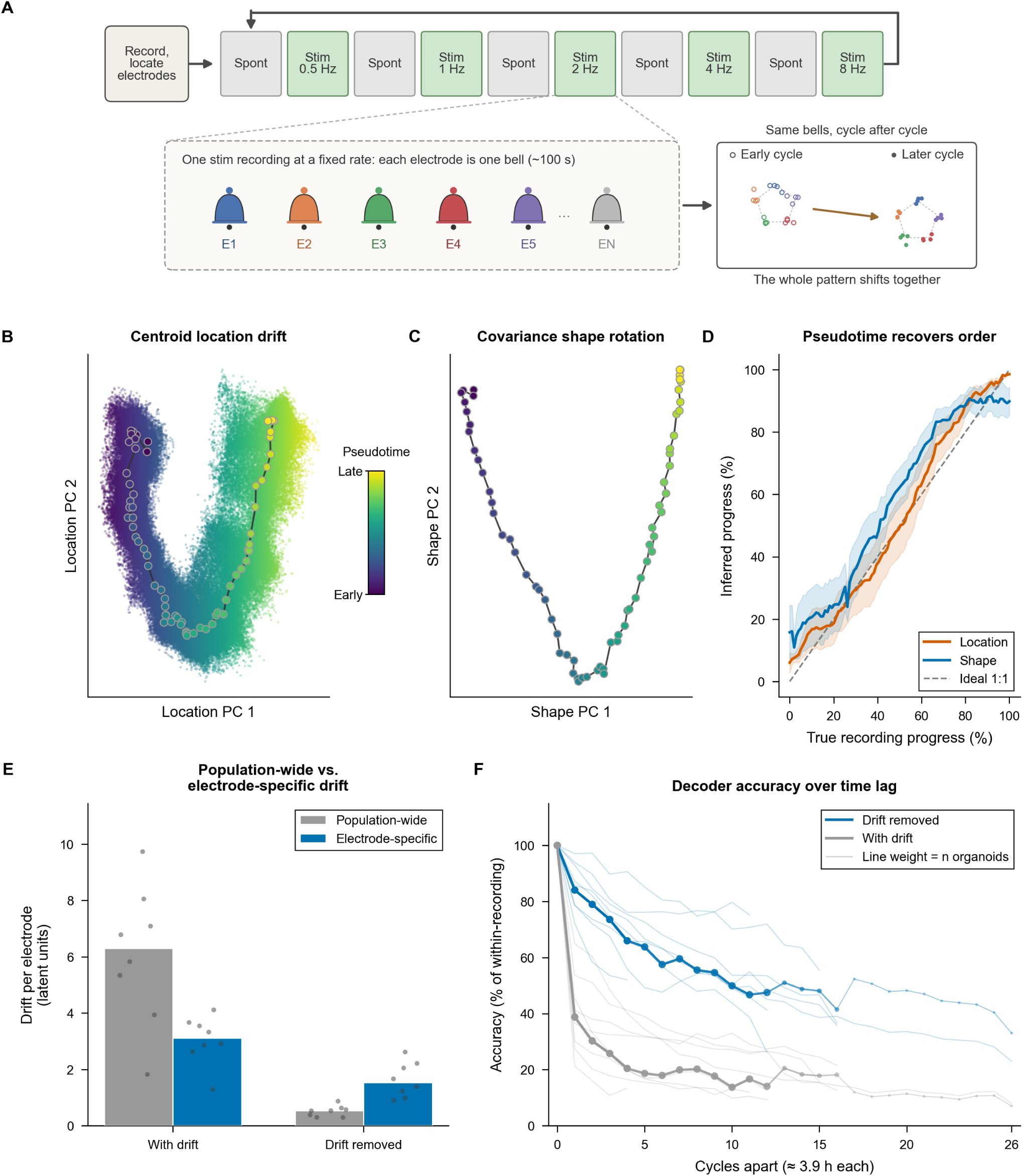
Ordered population drift is separate from electrical stimulus-response structure. (A) Stimulation protocol: 10 min spontaneous recordings alternate with stimulation recordings at a fixed rate, in which each electrode is stimulated alone for about 100 s; one pass through the five rates (0.5 to 8 Hz) is a cycle. Data shown are for illustration only. (B) Centroid location drift across the 65 spontaneous recordings of one experiment, colored from early to late. (C) Covariance shape rotation for the same recordings (multidimensional scaling on pairwise distances); color scale shared with (B). (D) Pseudotime from centroid location and from covariance shape against true recording progress; line, mean; band, SEM; dashed line, identity (*n* = 10 organoids). (E) Between-cycle drift of each electrode’s evoked response, split into the part shared by all electrodes (population-wide) and the remainder (electrode-specific), for the uncorrected (with drift) and baseline-subtracted (drift removed) readouts (*n* = 8 organoids). Bars, median; dots, organoids. (F) Site-decoder accuracy from the stim-masked latent readout, relative to accuracy within a recording, against cycles apart, with drift removed (per-pulse baseline subtraction) and with drift. Bold lines, median across organoids; thin lines, individual organoids; line weight, number of organoids at each lag (*n* = 8 organoids at 0–4 cycles, 7 at 5–11, 6 at 12, 4 at 13–15, 1 to 3 beyond). See also Figures S7–S10 and Tables S14 and S15.

Which electrode had been stimulated was decodable above chance from the latent readout with all stimulation inputs hidden (the stim-masked readout) at every frequency (cohort median 2.8 to 4.1× chance) and with no change as the stimulation frequency rose (paired 8 vs. 0.5 Hz, 2*/*8 organoids higher, *P* = 0.31; Figure S9A).

Across each experiment, the latent state drifted steadily. Its location (the network’s average state) shifted, and its shape (the covariance of the latent states around that average, that is, which dimensions vary together) rotated from one recording to the next (Figures 5B and 5C). Site decoding nonetheless held when the decoder was refit within each cycle (Figure S9A; Table S15). We then asked whether this drift was ordered, borrowing the concept of pseudotime from trajectory inference in single-cell transcriptomics^79^. Each spontaneous recording’s centroid and covariance were reduced to a single position along the dominant axis of drift, then recordings were ordered by that position alone. This ordering recovered the true chronological order (median Spearman *ρ* = 0.96 [IQR 0.96–0.98], range 0.88 to 0.99, *n* = 10 organoids; Figure 5D), a pattern also recovered in the binned firing rates (Figures S10B–S10D). The drift was also bounded. The mean-squared displacement (MSD) of recording centroids grew sublinearly with lag (floor-corrected exponent *α* ≈ 0.77; range 0.5–1.0, *n* = 10 organoids), consistent with confined wandering rather than free diffusion (*α* = 1) or directed motion (*α >* 1).

We measured how this drift affected the electrode code by fitting the LDA decoder on one cycle and testing it on every other cycle, using two readouts (Figure S9B). The uncorrected readout decodes the evoked latent state together with the global drift; the per-pulse readout removes that drift by subtracting each response’s own pre-stimulus baseline. Most of the drift was global (median 76% shared across all electrodes), and the per-pulse readout removed nearly all of it, leaving 9% of its residual drift shared (Figure 5E). At 7 to 12 cycles apart (median 32 h), the per-pulse readout declined gradually, from 3.43 to 1.86 chance, whereas the uncorrected readout fell from 12.55 to 2.29 and declined faster in all *n* = 7 organoids (*P* = 0.016; Figures 5F and S9C; Table S15; controls in Figures S9D–S9G). Electrode-specific structure therefore changed far more slowly than the global reorganization.

### Nonlinear responses to stimulation remain inside the spontaneous sub-space

Selection predicts that stimulation-evoked activity stays within the dimensions spontaneous activity already occupies, whereas instruction predicts new ones^5,6,21–26^. The two are ends of a continuum, and *in vivo* no naive baseline exists by the time activity can be recorded. Organoids provide that baseline, and projecting spontaneous and evoked activity into PULSE’s latent space places both in one coordinate system. We applied inputs of increasing complexity to the same networks, from open-loop stimulation to a closed-loop balancing task.

We first asked whether the network passively relayed stimulation or transformed it. A memoryless system, even a nonlinear one, can add harmonics (2*f*, 3*f*) to its input but cannot generate subharmonics. A subharmonic at half the stimulation frequency (*f/*2) requires memory that outlasts one stimulus cycle, so that the response alternates between successive pulses (Figure 6A). We projected each stimulation recording’s latent trajectory onto the axis along which stimulation displaces the population, estimated its power spectrum with a Lomb–Scargle periodogram suited to unevenly sampled timepoints^80,81^, and scored *f/*2 power against off-harmonic control frequencies (Figure 6B). At 4 Hz, *f/*2 power did not exceed the controls, consistent with passive following^82^; at 8 Hz it rose to a median 3.17 (7 of 8 organoids, *P* = 0.016; Figure 6C; Table S16). Poisson surrogates that reproduced each neuron’s response to the stimulation frequency and its harmonics, but carried no memory between pulses, abolished the rhythm, so the model did not create it (Figure S11A). The rhythm was also not an artifact of time binning (Figure S11C; Table S17).

**Figure 6:**
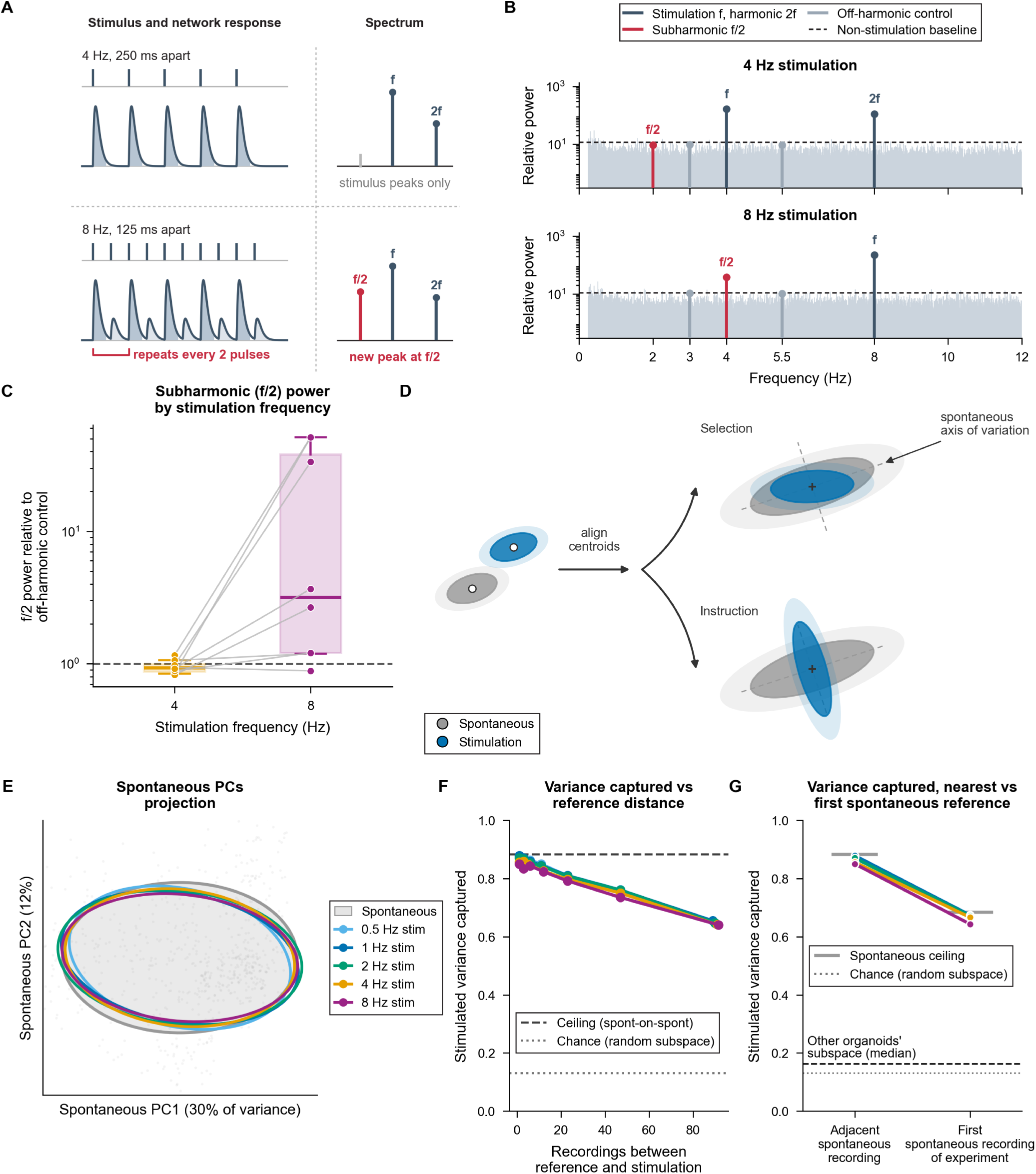
Nonlinear stimulus responses remain within the spontaneous subspace. (A) At 4 Hz every pulse evokes the same response, so the spectrum holds only the stimulation frequency and its harmonic (*f*, 2*f*), as the stimulus does. At 8 Hz a network whose memory outlasts one cycle responds weakly to every second pulse, so the pattern repeats every two pulses and places a peak at half the stimulation frequency (*f/*2). (B) Latent power spectrum for one organoid at 4 and 8 Hz stimulation. Markers give the stimulation frequency and its harmonic (*f*, 2*f*), the subharmonic (*f/*2), and two off-harmonic control frequencies (3.0 and 5.5 Hz); dashed line, 99.9th percentile of the recording’s power away from the marked frequencies. (C) *f/*2 power relative to the larger of two off-harmonic control frequencies (3.0 and 5.5 Hz) in the same recording, at 4 and 8 Hz; dashed line, equal power. Paired within organoid (*n* = 8 organoids); box, median and IQR; whiskers, 5th–95th percentiles. Two-sided Wilcoxon signed-rank test against 1. (D) Selection versus instruction schematic: spontaneous (gray) and stimulation (blue) latent clouds aligned on their centroids. (E) One experiment’s stimulation recordings, one per frequency, projected onto the top two principal components of the preceding spontaneous recording (gray cloud and ellipse). (F) Fraction of stimulation-recording variance inside the subspace of an earlier spontaneous recording against their separation, one curve per frequency (median across *n* = 8 organoids); the rightmost point pools references 65 or more recordings earlier. Dashed line, spontaneous-on-spontaneous ceiling; dotted line, chance. (G) The same fraction for the immediately preceding and the first spontaneous recording, one series per frequency (median across *n* = 8 organoids). Gray bars, spontaneous-on-spontaneous ceiling at each reference; dashed line, median fraction captured by other organoids’ spontaneous subspaces; dotted line, chance. One-sided Wilcoxon signed-rank test against chance. See also Figures S11 and S12.

If stimulation recruited existing spontaneous dimensions rather than creating new ones, its activity should fall inside the spontaneous subspace. We therefore projected each stimulation recording onto the subspace holding 90% of the preceding spontaneous recording’s variance. We normalized the captured variance between chance (a randomly oriented subspace of the same size) and a ceiling (one spontaneous recording projected onto the preceding one; Figures 6D and 6E). Stimulation-evoked activity fell almost entirely inside the spontaneous subspace, reaching a median 98.5% of the ceiling and exceeding chance in all eight organoids (one-sided *P* = 0.0039; Figures 6F and 6G; Table S16). This held at every stimulation frequency, including the 8 Hz stimulation that generated the subharmonic (Figure 6G). It also held at every subspace size from 1 to 256 dimensions and in binned firing rates (Figures S12A–S12D). Other organoids’ spontaneous subspaces captured barely more than chance (Figure 6G; Table S17).

Both spontaneous and stimulation activity reorganized over an experiment (Figures 5B, 5C and 5E), so we asked whether the overlap held as the spontaneous subspace drifted. Spontaneous recordings lost more than a third of their overlap with each other between the most distant pairs, declining in all 8 organoids (*P* = 0.0078; Figure S12E; Table S16). Against the experiment’s first spontaneous recording, stimulation overlap fell, but spontaneous overlap fell equally, so stimulation still reached 98.2% of the spontaneous ceiling (Figure 6G; Table S16). Stimulation therefore stayed inside the spontaneous subspace as that subspace drifted.

### Task control selects from spontaneous dynamics

Open-loop stimulation showed that stimulation-evoked activity reuses spontaneous dimensions, but a fixed pulse train makes no demand on the network beyond absorbing a pattern. In a closed-loop task, by contrast, the network’s output determines its next input.

To distinguish selection from instruction in task-oriented network activity, we analyzed 13 organoids under three regimes of increasing complexity (Figure 7A): *spontaneous activity*; *open-loop stimulation*, in which fixed pulse trains are delivered without feedback; and *closed-loop cartpole control*, in which stimulation encodes the angle of a virtual balancing pole and the network’s firing is decoded into the force applied to the cart^8^. We compared the regimes’ clouds of latent states (Figure 7B) by the location and shape measures used for drift, each expressed as a multiple of how far a single regime’s cloud moves within one recording (Figure 7C).

**Figure 7:**
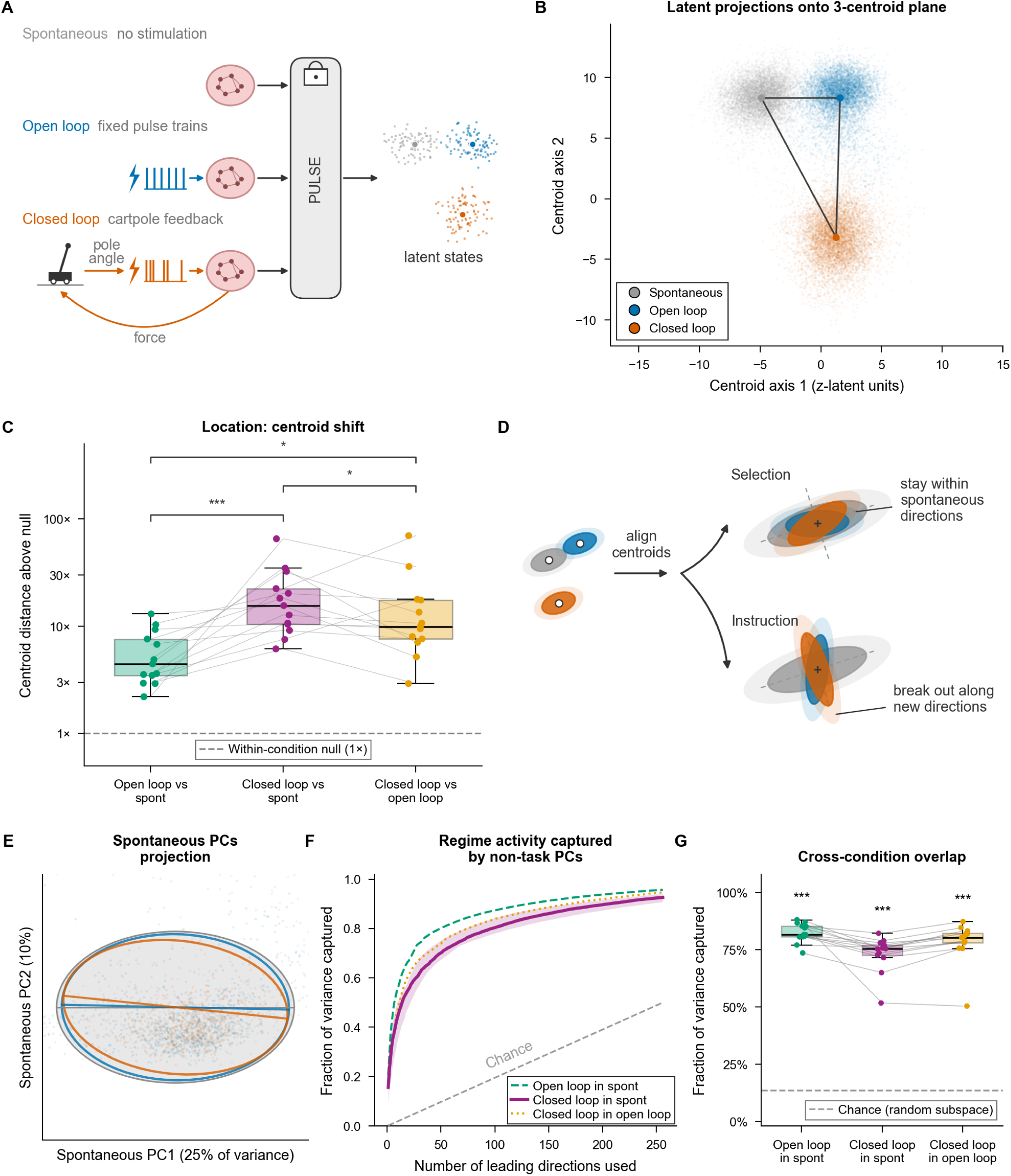
Task engagement recruits spontaneous dimensions rather than constructing new dynamics. (A) Schematic of three experimental regimes: spontaneous activity, open-loop stimulation and closed-loop cartpole control recorded from the same organoid. (B) Example of latent states from the three regimes projected onto the plane through their centroids; the triangle connecting them marks the between-centroid distances. (C) Centroid shift between regime pairs relative to the within-condition null (*n* = 13 organoids; dots, organoids, each the log-scale mean of its experiments; lines join organoids). Box, median and IQR; whiskers, 1.5 IQR. Brackets, paired Wilcoxon signed-rank tests on log_2_ ratios of the organoids, one-sided for the two contrasts of the spontaneous–closed-loop shift against the others and two-sided for the third, Holm-corrected; *^∗^P <* 0.05; *^∗∗∗^P <* 0.001. (D) Selection versus instruction schematic showing whether open-loop stimulation and closed-loop control recruit latent directions the spontaneous subspace already contains or constructs new ones. (E) Example of the spontaneous PC1–PC2 plane for one organoid, with the spontaneous covariance ellipse (gray) and the open-loop (blue) and closed-loop (orange) ellipses overlaid. (F) Cumulative fraction of each regime’s variance captured by the leading principal components of a non-task regime: closed loop in spontaneous (magenta; median and IQR band), open loop in spontaneous (green dashed, median) and closed loop in open loop (gold dotted, median); gray dashed line, chance (*k/*512; *n* = 13 organoids; each curve is the median across organoids of the organoids’ mean curves, band the IQR across organoids). (G) Fraction of variance captured for the same three pairs (*n* = 13 organoids; dots, organoids; lines join organoids; box as in C); dashed line, median random-subspace chance. Stars, one-sided Wilcoxon signed-rank test of the organoids’ values for each pair against its own chance, uncorrected (each *P* = 1.2 × 10*^−^*^4^, the smallest value possible for 13 organoids); *^∗∗∗^P <* 0.001. See also Figures S13 and S14 and Tables S19 and S21.

Closed-loop control moved the network’s average state furthest from spontaneous activity, a median 15.5 × within-recording movement against 4.4× for open-loop stimulation (*n* = 13 organoids, *P*_Holm_ ≤ 0.047; Figure 7C; Table S18). Global drift did not explain this shift. Closed-loop recordings up to 5 h apart were closer to each other in latent space than to spontaneous activity (organoid-level *P*_Holm_ ≤ 0.026; Figures S13A and S13F). The shape component followed the same ordering at a smaller scale (Figure S13B; Table S18).

A location shift this large would fit instruction, but it also fits selection, in which activity settles at a new operating point yet still fluctuates along directions spontaneous activity already used; instruction instead predicts new directions (Figure 7D). Using the same subspace analysis as in Figures 6D–6G, open-loop activity reached a median 95.0% and closed-loop activity 85.7% of the spontaneous ceiling (here defined by comparing two halves of the recording), far above chance in all 13 organoids (Figures 7E–7G; Table S19; Videos S2 and S3). The leading ten spontaneous components captured more open-loop than closed-loop variance (Figure 7F; Table S20), consistent with selection recruiting the same dimensions in different proportions.

The direction along which closed-loop control displaced the network from spontaneous activity also aligned with its open-loop response (mean cosine similarity 0.460 across organoids, against 0.227 for permutation nulls pairing each closed-loop direction with the open-loop response of a different organoid; *P* = 5 × 10*^−^*^5^, the resolution limit of 20,000 draws; 10 of the 13 organoids above the null mean; Figure S13C). This alignment persisted after removing each network’s firing-rate axis (across organoids, 0.414 against 0.212) and was stable between halves of an experiment (Figure S13D; Table S20). The subspace overlap was also present in binned firing rates, where all three pairwise comparisons exceeded their chance floor (median 86–88% captured versus ~50% chance; 10*/*10 organoids; Table S21; Figure S13E). The network’s representation of task-based information therefore falls closer to the selection end of the continuum.

Selection makes a further prediction: the direction that distinguishes success from failure should itself lie within the spontaneous subspace. Closed-loop play divides into episodes, each running from one pole failure to the next, so an episode’s length is how long the network kept the pole balanced. We summarized the network’s state as PULSE’s 512-dimensional latent averaged over non-overlapping 2 s windows of each episode. A logistic-regression classifier on these windows, tested on held-out episodes, separated sustained balance (episodes of ≥ 30 s) from quick failure (episodes of *<* 10 s) at a median area under the receiver operating characteristic curve (AUC) of 0.790 in the 12 organoids with enough labeled episodes (significant in 10 of 12 organoids; Figure S14A; Table S20). Windows were matched for pole angle, angular variability and firing rate^83^; additionally matching on time in the recording or since episode start left the separation significant in 6 of 7 organoids, so the decoder read the network’s control state rather than slow drift (Figures S14B and S14F; Table S17). A binned firing-rate decoder on the same windows was weaker but above chance (median AUC 0.703) and fell below the latent decoder in all 12 organoids (*P* = 4.9 × 10*^−^*^4^ across organoids). Failures collapsed onto a low-dimensional, stereotyped trajectory, while sustained balance explored higher-dimensional regions as episodes lengthened (1,387 episodes, three organoids; Figures S14C and S14D; Video S3).

The balance-versus-failure direction lay largely inside the spontaneous subspace. In the 10 of 13 organoids with enough episodes of each kind to define it, the leading spontaneous principal components spanned a median 62.7% of it, against 11.2% for a randomly oriented direction (10*/*10 organoids above chance, *P* = 9.8 × 10*^−^*^4^; Figure S14E). To test whether this direction reflected coordination among spontaneous dimensions rather than their individual variances, we shifted each dimension independently in time, which keeps each dimension’s variance but breaks their coordination. Spontaneous activity varied 9.7 ×more along the direction than it did after this shift (9 of 10 organoids above 2×). Binned firing rates, with no model in the path, reproduced the capture (93.1% at half the available dimensions, against 50% chance) but showed the coordination less clearly (2.6×, 7 of 10 organoids; Table S22). Directions separating randomly relabeled episodes were at least as aligned (Table S22), so the balance direction was not a dedicated axis but one of the directions spontaneous activity already used.

## DISCUSSION

Our results show that a single self-supervised model, pretrained on large-scale HD-MEA recordings from mouse cultures and slices, learns a representation that places networks across development, genotypes and tasks in one coordinate system.

A developmental axis fit on PULSE’s latent state from one control line transferred without refitting to other lines and protocols, separating disorder-specific effects from differences in procedure. Two different mutations, *SHANK3* deficiency and a 16p11.2 deletion, converged on a shared axis of network coordination in independently prepared comparisons (significant jointly, although the *SHANK3* comparison alone was not), suggesting that the behavioral convergence observed across NPD risk loci^15–17^ may extend to circuit-level dynamics. Large-scale efforts such as the PsychENCODE Consortium^84^, the BRAIN Initiative Cell Census Network^85^, and the Scalable and Systematic Neurobiology of Psychiatric and Neurodevelopmental Disorder Risk Genes (SSPsyGene) consortium are now characterizing high-confidence NPD risk genes using diverse protocols and modalities at scale. As these consortia release large, well-annotated datasets, a shared model of network activity could measure effects across disorders in common units and help explain how genetic risk reshapes circuit development.

How the brain transitions from spontaneously generated activity to represented information has long been debated^21,27–30^. Mature cortex reuses its spontaneous patterns when representing sensory input^22–24^; using PULSE’s shared coordinate system, we show that naive organoid networks, with no sensory history, already reuse spontaneously generated patterns during closed-loop control of an inverted pendulum. Over days of recording, electrode-specific responses persisted while the population as a whole drifted, suggesting that site-specific structure and global network state change on different timescales. The lower predictability of 16p11.2-deletion networks raises the possibility that this mutation destabilizes network activity^86^ in ways that would compromise task performance. Extending closed-loop tasks to NPD organoids could therefore measure directly how genetic perturbations change a network’s capacity to represent and act on input.

### Limitations of the study

Our model relies on the Mamba-2 state-space architecture^43^; transformers or hybrid designs^36–38,87^ may capture different aspects of neural dynamics. The current work focuses on cortical networks derived from a limited number of genetic backgrounds; extending pretraining to organoids modeling other brain regions and incorporating a broader panel of NPD-associated mutations will be necessary to test the generality of the convergent phenotypes we observe. The 16p11.2 comparison is also between donors rather than isogenic lines, so part of that difference may reflect genetic background. Planar electrodes subsample the network, which three-dimensional electrode arrays^88,89^ could address. Finally, the approach is not limited to electrical stimulation or *in vitro* HD-MEA data: optogenetic or chemogenetic perturbation data would enable causal dissection of the latent features the model learns, and public *in vivo* electrophysiology data^90^ could extend pretraining to intact circuits.

## RESOURCE AVAILABILITY

## Lead contact

Requests for further information and resources should be directed to and will be fulfilled by the lead contact, Mohammed A. Mostajo-Radji.

## Materials availability

This study did not generate new unique reagents.

## ACKNOWLEDGMENTS

We thank Kristof Tigyi and Demir Ozcakir for sharing mouse embryonic stem cell and human iPSC lines, Samira Vera-Choqqueccota for culture maintenance, the laboratory of Bin Chen (UC Santa Cruz) for providing postnatal mouse pups, and Jesus Gonzalez-Ferrer for bioinformatic support. We would also like to thank Kateryna Voitiuk for assistance with hardware setup and maintenance.

We acknowledge the National Research Platform (National Science Foundation [NSF] CNS1730158, ACI1540112, ACI1541349, OAC1826967), the University of California Office of the President (UCOP), and the UCSD Calit2/Qualcomm Institute. Imaging was provided by the UCSC Microscopy Core (RRID: SCR 021135).

This work was supported by Schmidt Futures (SF857 to S.R.S., D.H., and M.T.); the National Human Genome Research Institute (RM1HG011543 to S.R.S., D.H., and M.T.); the NSF (2134955 to S.R.S., D.H., and M.T.; 2034037 to M.T.; 2515389 to D.H., M.T., and M.A.M.-R.); the National Institute of Mental Health (SSPsyGene Consortium, U24MH132628 to D.H. and M.A.M.-R.); the National Institute of Neurological Disorders and Stroke (U24NS146314 to D.H. and M.A.M.-R.); the California Institute for Regenerative Medicine (CIRM; DISC4-16285 to S.R.S., A.A.P., M.T., and M.A.M.-R.; DISC4-16337 to A.B. and M.A.M.-R.; and DISC4-19334 to M.A.M.-R.); UCOP (M25PR9045 to S.R.S., M.T., and M.A.M.-R.); and the Brain and Behavior Research Foundation (33184 to M.A.M.-R.). H.E.S. was partially supported by the NSF Graduate Research Fellowship Program (GRFP). K.H. was partially supported by the Undergraduate Research in Science & Technology (URST) Award and the Koret Scholarship from UC Santa Cruz. The content is solely the responsibility of the authors and does not necessarily represent the official views of the National Institutes of Health, NSF, the University of California, CIRM, or any other agency of the State of California.

## AUTHOR CONTRIBUTIONS

H.E.S., A.R., M.T., and M.A.M.-R. conceptualized the project. H.E.S., A.R., S.H., A.M.O., and K.H. designed and ran experiments. H.E.S., S.H., A.M.O., I.C., K.D., and M.M. generated and maintained the cultures. Z.P.P., J.D., A.A.P., A.B., and S.R.S. contributed critical reagents. D.H., M.T., and M.A.M.-R. supervised the work and secured funding. H.E.S., S.H., A.M.O., and M.A.M.-R. wrote the manuscript with input from all authors.

## DECLARATION OF GENERATIVE AI AND AI-ASSISTED TECHNOLOGIES

During the preparation of this work, the authors used Claude (Anthropic) in order to refine phrasing and assist with manuscript organization. After using this tool or service, the authors reviewed and edited the content as needed and take full responsibility for the content of the publication.

## SUPPLEMENTAL INFORMATION INDEX

Figures S1–S14, Tables S1–S9 and S11–S22 and their legends in a PDF

Table S10. Genome-wide normalized expression of the bulk RNA-seq organoid samples, related to Figure 3 (Excel file)

Video S1. PULSE predicts population activity 20 ms ahead in a withheld validation recording, related to Figure 2

Video S2. Spontaneous, open-loop and closed-loop activity of one organoid on its spontaneous map, related to Figure 7

Video S3. Closed-loop cartpole control by the same organoid on its spontaneous map, related to Figure 7

## STAR METHODS

## Key resources table

## Experimental model and study participant details

All procedures using mice were performed under protocols approved by the Institutional Animal Care and Use Committee of the University of California Santa Cruz. Research with the human induced pluripotent stem cell lines was approved by the Institutional Review Board of the University of California Santa Cruz (HS-FY2026-78).

### Sex of cell lines and animals

All three mESC lines are male, so every mouse organoid and 2D culture in the pretraining corpus is male. The sex of the postnatal pups used for acute slices was not determined. The *SHANK3*^+/+^ and *SHANK3*^+/^*^−^* lines share the male KOLF2.2J background, so the isogenic comparison is matched for sex. In the 16p11.2 comparison, the control line CW20107 is female and 14765-X2 is male. The two lines also come from different donors, so donor sex, genetic background and 16p11.2 genotype cannot be separated in that comparison. With a single hiPSC line per genotype, this study could not assess the influence of sex on network dynamics, which is a limitation.

### Mouse embryonic stem cell lines

Mouse dorsal-forebrain (DF) organoids contributing to the pretraining corpus were generated from three male mESC lines: BRUCE4 (RRID:CVCL K037), ES-E14TG2a (RRID:CVCL 9108), and KH2 (RRID:CVCL C317), following the protocol in Hernandez et al. ^5^.

### mESC maintenance

mESC lines were maintained as described by Hernandez et al. ^5^. mESCs were maintained on plates precoated for 15 min at room temperature with 0.5 µg/mL recombinant human vitronectin (Thermo Fisher Scientific # A14700) in 1× PBS (pH 7.4; Thermo Fisher Scientific # 70011044). Cells were cultured in mESC maintenance medium consisting of Glasgow Minimum Essential Medium (GMEM; Thermo Fisher Scientific # 11710035) supplemented with 10% embryonic stem cell-qualified fetal bovine serum (Thermo Fisher Scientific # 10439001), 0.1 mM MEM Non-Essential Amino Acids (Thermo Fisher Scientific # 11140050), 1 mM sodium pyruvate (Millipore Sigma # S8636), 2 mM GlutaMAX supplement (Thermo Fisher Scientific # 35050061), 0.1 mM 2-mercaptoethanol (Millipore Sigma # M3148), 0.05 mg/mL Primocin (InvivoGen # ant-pm-05), and 1000 U/mL recombinant mouse leukemia inhibitory factor (Millipore Sigma # ESG1107), with daily medium changes. Cells were passaged using ReLeSR (Stem Cell Technologies # 05872) according to manufacturer instructions and cryopreserved in mFreSR medium (Stem Cell Technologies # 05855).

### Mouse dorsal-forebrain organoid generation

mESCs were dissociated into single cells using TrypLE Express Enzyme (Thermo Fisher Scientific # 12604021) for 5 minutes at 37°C. After dissociation, the cells were re-aggregated in Lipidure-coated 96-well V-bottom plates at a density of 3,000 cells per well in 150 µL of mESC maintenance medium, supplemented with 10 µM Rho Kinase Inhibitor Y-27632 (Tocris Bioscience # 1254) and 1,000 units/mL Recombinant Mouse Leukemia Inhibitory Factor (Millipore Sigma # ESG1107). Following 24 hours of re-aggregation, the medium was replaced with forebrain patterning medium composed of DMEM/F12 with GlutaMAX (Thermo Fisher Scientific # 10565018), 10% Knockout Serum Replacement (Thermo Fisher Scientific # 10828028), 0.1 mM MEM Non-Essential Amino Acids (Thermo Fisher Scientific # 11140050), 1 mM Sodium Pyruvate (Millipore Sigma # S8636), 1X N-2 Supplement (Thermo Fisher Scientific # 17502048), 2X B-27 minus Vitamin A (Thermo Fisher Scientific # 12587010), 0.1 mM 2-Mercaptoethanol (Millipore Sigma # M3148), and 0.05 mg/mL Primocin (InvivoGen # ant-pm-05).

For dorsal forebrain patterning, the medium was further supplemented with 10 µM Rho Kinase Inhibitor Y-27632 (Tocris Bioscience # 1254), 5 µM WNT inhibitor XAV939 (StemCell Technologies # 100-1052), and 5 µM TGF-*β* inhibitor SB431542 (Tocris Bioscience # 1614). Medium was changed daily, with N-2 and B-27 supplements added post-filtration to preserve hydrophobic components. On day 5, organoids were transferred to ultra-low adhesion plates (Millipore Sigma # CLS3471) containing fresh neuronal differentiation medium and maintained on an orbital shaker at 68 rpm.

From days 6 to 12, progenitor expansion medium consisted of DMEM/F12 with GlutaMAX (Thermo Fisher Scientific # 10565018), BrainPhys Neuronal Medium (Stem Cell Technologies # 05790), 1X B-27 minus Vitamin A, 1X N-2 Supplement, 0.1 mM MEM Non-Essential Amino Acids, 0.05 mg/mL Primocin (InvivoGen # ant-pm-05), and 200 µM Ascorbic Acid (Millipore Sigma # 49752). Organoids were cultured under 5% CO_2_ with medium changes every 2–3 days.

From day 15 onward, neural maturation medium contained BrainPhys Neuronal Medium supplemented with 1X B-27 Plus Supplement (Thermo Fisher Scientific, # A3582801), 1X N-2 Supplement, 1X Chemically Defined Lipid Concentrate (Thermo Fisher Scientific # 11905031), 5 µg/mL Heparin (Millipore Sigma # H3149), and 0.05 mg/mL Primocin (InvivoGen # ant-pm-05). The medium also included 200 µM Ascorbic Acid until day 25. Medium was changed every 2–3 days with organoids maintained at 60 rpm (16 organoids per well) to minimize fusion.

### Mouse ventral-forebrain organoid generation

Ventral-forebrain (VF) organoids were generated using the same protocol as dorsal-forebrain organoids above, with the following modifications: 250 nM LDN193189 (StemCell Technologies # 72147) was added from days 0–5, and 100 nM PD0325901 (StemCell Technologies # 72184) and 100 nM SAG (Millipore Sigma # SML1314) were included from days 0–14.

### Two-dimensional dissociated cultures

Day 25 organoids were dissociated using the Worthington Papain Dissociation System (Worthington Biochemical # LK003150) following the manufacturer’s instructions, and 100,000 dissociated cells were plated onto each MaxOne HD-MEA.

### Human iPSC maintenance

Four human iPSC lines formed two independent comparisons. For the isogenic *SHANK3* comparison, the control *SHANK3*^+/+^ line was KOLF2.2J, a male (age 55–59, European ancestry) human induced pluripotent stem cell (hiPSC) line^91^ (The Jackson Laboratory #JIPSC003902; RRID:CVCL F4CV). The heterozygous loss-of-function *SHANK3*^+/^*^−^* line was generated by cytosine base editing (CBE) in KOLF2.2J, introducing a G*>*A substitution at chr22:50,679,170 (hg38) that converts a TGG (Trp) codon to a premature stop codon. The generation and characterization of the mutant line is described in detail by Zhang et al. ^56^.

For the 16p11.2 comparison, the unaffected control line CW20107 (RRID:CVCL JH62) is a female (age 21, European ancestry) hiPSC line obtained from the California Institute for Regenerative Medicine (CIRM) iPSC repository^92^. The disease line 14765-X2 (Rutgers RUID SV0001455) was derived from a male proband (age 12) carrying a heterozygous 16p11.2 microdeletion; fibroblasts were obtained from the Simons Variation in Individuals Project (Simons VIP) and reprogrammed to iPSCs using episomal vectors as described by Deshpande et al. ^55^.

hiPSCs were maintained on plates coated with Geltrex LDEV-Free Reduced Growth Factor Basement Membrane Matrix (Thermo Fisher Scientific # A1413201) diluted to 2% (v/v) in DMEM/F12 with GlutaMAX (Thermo Fisher Scientific # 10565018) according to manufacturer instructions. Cells were cultured in StemFlex medium (Thermo Fisher Scientific # A3349401), consisting of StemFlex basal medium supplemented with the StemFlex supplement and 0.05 mg/mL Primocin (InvivoGen # ant-pm-05), with medium changes every other day. When passaging, cells were dissociated using Accutase (Fisher Scientific # NC9839010) and replated in medium supplemented with 10 µM Rho kinase inhibitor Y-27632 (Tocris Bioscience # 1254). Cells were cryopreserved in Bambanker standard freezing medium (Bulldog Bio # BB01, Cat # 104085-050). All cell lines tested negative for *Mycoplasma* by sequencing (Plasmidsaurus).

### Human dorsal forebrain organoids by direct aggregation

Organoids were generated as previously described^5,93^. hiPSCs were dissociated into single cells using Accutase (Fisher Scientific # NC9839010) and re-aggregated in 96-well Slit-Well plates (Sbio # MS-9096SZ) at a density of 9,000 cells per well in StemFlex medium supplemented with 10 µM Rho Kinase Inhibitor Y-27632 (Tocris Bioscience # 1254).

From days 0 to 5, organoids were cultured in patterning medium composed of DMEM/F12 with GlutaMAX (Thermo Fisher Scientific # 10565018), 1X B-27 minus Vitamin A (Thermo Fisher Scientific # 12587010), 0.1 mM MEM Non-Essential Amino Acids (Thermo Fisher Scientific # 11140050), 1 mM Sodium Pyruvate (Millipore Sigma # S8636), 1X Chemically Defined Lipid Concentrate (Thermo Fisher Scientific # 11905031), and 0.05 mg/mL Primocin (InvivoGen # ant-pm-05).

For dorsal forebrain patterning, the medium was further supplemented with 10 µM Rho Kinase Inhibitor Y-27632 (Tocris Bioscience # 1254), 2.5 µM WNT inhibitor XAV939 (StemCell Technologies # 100-1052), and 10 µM TGF-*β* inhibitor SB431542 (Tocris Bioscience # 1614). Medium was changed daily, with B-27 added post-filtration.

From days 6 to 35, progenitor expansion medium consisted of a 1:1 mixture of DMEM/F12 with GlutaMAX (Thermo Fisher Scientific # 10565018) and BrainPhys Neuronal Medium (Stem Cell Technologies # 05790), supplemented with 1X B-27 minus Vitamin A, 0.1 mM MEM Non-Essential Amino Acids, 1X Chemically Defined Lipid Concentrate (Thermo Fisher Scientific # 11905031), 0.5 mM additional GlutaMAX supplement (Thermo Fisher Scientific # 35050061), 0.05 mg/mL Primocin (InvivoGen # ant-pm-05), and 200 µM Ascorbic Acid (Millipore Sigma # 49752). Organoids were cultured under 5% CO_2_ with medium changes every 2–3 days onward.

From day 35 onward, neural differentiation medium contained BrainPhys Neuronal Medium (Stem Cell Technologies # 05790) supplemented with 1X B-27 (B-27 serum free from days 35–70 [Thermo Fisher Scientific # 17504044], switching to B-27 Plus Supplement [Thermo Fisher Scientific # A3582801] from day 70 onward), 1X Chemically Defined Lipid Concentrate (Thermo Fisher Scientific # 11905031), and 0.05 mg/mL Primocin (InvivoGen # ant-pm-05). The medium also included 200 µM Ascorbic Acid until day 50. B-27 was added to filtered medium in 1 mL aliquots per 50 mL.

### Human dorsal forebrain organoids from neural progenitors

hiPSCs were grown until approximately 80% confluency in a 6-well plate (Corning # 3516), where the media was changed to patterning media supplemented with 10 µM Rho Kinase Inhibitor Y-27632 (Tocris Bioscience # 1254), 2.5 µM WNT inhibitor XAV939 (StemCell Technologies # 100-1052), and 10 µM TGF-*β* inhibitor SB431542 (Tocris Bioscience # 1614). Medium was changed daily.

Once neural rosettes were observed (~day 10), cells were dissociated using Accutase (Fisher Scientific # NC9839010) and transferred to a 6-well low-attachment plate (Corning # 3471) to form neurospheres. Patterning medium was supplemented with FGF2 (PeproTech # 100-18B) and EGF (Thermo Fisher Scientific # PHG0311) and, on the day of replating only, Rho kinase inhibitor Y-27632 (Tocris Bioscience # 1254). Medium was changed every other day.

Once neurospheres reached approximately 250 µm in diameter, they were plated onto Geltrex-coated 6-well plates and grown to 100% confluency in patterning medium supplemented with FGF2 (PeproTech # 100-18B), EGF (Thermo Fisher Scientific # PHG0311) and, on the day of replating only, Rho kinase inhibitor Y-27632. Medium was changed every day. After reaching 100% confluency, wells were split 1:3.

To form organoids, neural progenitors were dissociated using Accutase and re-aggregated in 96-well Slit-Well plates (Sbio # MS-9096SZ) at a density of 20,000 cells per well in progenitor expansion medium supplemented the day of aggregation with 10 µM Rho Kinase Inhibitor Y-27632. Medium was changed every 3 days.

On day 14 organoids were changed to neural differentiation medium containing 1X B-27 minus Vitamin A. From day 35 onward, organoids were changed to neural differentiation medium containing 1X B-27 Plus Supplement. Medium was changed every 3 days.

### Mouse primary tissue dissection

Electrophysiological slices were prepared from postnatal day 0–3 C57BL/6 mice according to Tsyporin et al. ^94^. Briefly, pups were anesthetized on ice and decapitated. Slices were cut in ice-cold ACSF (containing 126 mM NaCl, 2.5 mM KCl, 26 mM NaHCO_3_, 2 mM CaCl_2_, 1 mM MgCl_2_, 1.25 mM NaH_2_PO_4_ and 10 mM glucose, saturated with 95% O_2_ and 5% CO_2_). Coronal sections (300 µm) were obtained and washed with 1× D-PBS before transfer to neural differentiation medium and kept in standard incubator conditions for 2-3 days in a 24-well low attachment plate (Corning # 3473) before plating on MaxOne HD-MEAs.

## Method details

### Bulk RNA sequencing

CW20107 and 14765-X2 organoids were collected at day 180 and KOLF2.2J organoids at day 100. Each sample was a pool of three organoids, from which total RNA was extracted with the PureLink RNA Mini Kit (Thermo Fisher Scientific # 12183025) according to the manufacturer’s instructions. Libraries were prepared and sequenced by Plasmidsaurus with their 3′ end counting RNA-seq workflow, in which reverse transcription from a poly(dT) primer tags each transcript with a unique molecular identifier (UMI), followed by stranded single-end sequencing on an Illumina NovaSeq X Plus (one 94-base cDNA read per fragment with the UMI in the read name, enriched within the last approximately 400 nucleotides of each transcript). KOLF2.2J samples (*n* = 4 *SHANK3*^+/+^, *n* = 4 *SHANK3*^+/^*^−^*) came from two sequencing runs and CW20107 and 14765-X2 samples (*n* = 3 each) from a third. Plasmidsaurus processed the reads: fastp (v0.24.0)^95^ trimmed poly-X tails and low-quality 3′ ends, STAR (v2.7.11)^96^ aligned reads to the human reference genome GRCh38 (hg38), UMIcollapse (v1.1.0)^97^ removed PCR duplicates by UMI, and featureCounts (subread v2.1.1)^98^ counted reads per gene over exons and 3′ untranslated regions, strand-specifically, with fractional assignment of multi-mapping reads. Expression was normalized within each sample to counts per million (CPM; a gene’s UMI-deduplicated count divided by the sample’s total gene-assigned count, times 10^6^), with no between-sample normalization; genes with several Ensembl identifiers were summed by gene symbol for the figures. Table S10 lists CPM for every Ensembl gene with a nonzero count in at least one of the 14 samples.

### Electrophysiological preparation

MaxOne HD-MEAs were coated with 0.01% polyethylenimine (PEI; Millipore Sigma #408727) reconstituted in 1× borate buffer (Fisher Scientific #28341) for 1 h at 37 *^◦^*C, followed by three washes with deionized water and air-drying for 10 min, then with 20 µg*/*mL mouse laminin (Fisher Scientific #CB40232) and 5 µg*/*mL human fibronectin (Fisher Scientific #CB40008) in 1× PBS for 1 h at 37 *^◦^*C. Organoids were placed on coated HD-MEAs, excess medium was removed, and samples were incubated at 37 *^◦^*C for 5 min to promote adhesion before adding pre-warmed neuronal differentiation medium.

MaxOne HD-MEAs (Maxwell Biosystems #PSM) provide 26,400 recording sites with simultaneous readout from up to 1,024 channels. Activity was first sampled across the array in a sweeping checkerboard of 1,024-electrode configurations (30 s each); the 1,020 most active electrodes with minimum 50 µm spacing were then selected for recording to ensure single-unit resolution. Neuronal activity was recorded at 20 kHz from these electrodes under standard incubation conditions (5% CO_2_, 37 *^◦^*C), with a 1 Hz hardware high-pass filter, and saved in HDF5 format. Spontaneous recordings were interleaved with stimulation-evoked recordings.

### Pharmacological treatments

Activity in 60- to 65-day-old dorsal-forebrain organoids was recorded before and after treatment with Gabazine (SR95531; Abcam #ab120042; 1 µM), NBQX (Abcam #ab120045; 20 µM), or APV (DL-AP5; Selleck Chemicals #E2979; 100 µM), with dimethyl sulfoxide (DMSO; Cell Signaling Technology #12611S), the vehicle for drug treatments, as a control. The recording protocol consisted of a 10-minute baseline recording, followed by a 30-minute drug incubation period, and a 10-minute post-incubation recording.

### Electrophysiological spike sorting and data preprocessing

Electrophysiological activity was monitored every 2–3 days using Maxwell Biosystems acquisition software. Raw extracellular recordings were band-pass filtered from 300 to 6000 Hz and spike-sorted using Kilosort2^99,100^ through a custom Python pipeline. Quality control excluded units with interspike interval violation rates exceeding 0.5, mean firing rates below 0.1 Hz, or signal-to-noise ratios (SNR) below 3.

Recordings acquired during electrical stimulation were sorted with RT-Sort^101^, paired with a custom Python artifact-removal algorithm^8^. RT-Sort proceeded in two stages: propagation-sequence detection, run once per electrode configuration, followed by template-based sorting of every recording sharing that configuration.

### Real-time acquisition and processing workstation

Real-time acquisition, online spike sorting, and stimulation control ran on a dedicated workstation. Voltage signals from the MaxOne HD-MEA were streamed to the workstation and acquired with MaxLab Live (Maxwell Biosystems), with acquisition and stimulation controlled programmatically through the MaxLab Python API. Incoming voltage traces were preprocessed and spike-sorted online with RT-Sort^101^. The workstation comprised a single NVIDIA GeForce RTX 4090 GPU (24 GB GDDR6X), an Intel Core i9-14900K CPU (24 cores, 32 threads), 128 GB of DDR5 memory and an ASUS ProArt Z790-Creator WiFi motherboard (Intel Z790 chipset) with onboard 10 Gb/s Ethernet. The MaxOne recording unit was connected to the workstation over a direct category 6A ethernet cable. Online RT-Sort output was used only for real-time control; all recordings were re-sorted offline for analysis as described above.

### Electrical stimulation protocol

Stimulation-evoked recordings were either acquired as part of the closed-loop experiments reported previously^8^ or recorded on the same platform under similar parameters, using the real-time workstation described above. All stimulation used charge-balanced biphasic square-wave pulses of 400 mV amplitude, delivered through the on-chip stimulation units of the MaxOne array; causal sweeps used a 200 µs phase duration. In the causal sweep paradigm each candidate site is pulsed in turn at 2 Hz for a contiguous 24.5 s stretch of 50 pulses while the population response is recorded; the electrode-discrimination cohort consists of the causal sweeps that screen candidate encoder electrodes ahead of each closed-loop experiment. Tetanic and closed-loop paradigms share this amplitude and waveform but differ in phase duration, electrode selection and pulse timing; in closed-loop experiments, pulse rate encoded the task variable.

All recordings, including those from the closed-loop study, were spike-sorted uniformly for this work, enabling analysis at the level of individual units.

### Stimulation-rate ladder protocol

Ladder experiments alternated spontaneous and stimulation recordings continuously, each stimulation recording preceded by a spontaneous recording of 600 s. A stimulation recording (median 1841 s) stepped through a fixed list of 7 to 32 electrodes, pulsing each in turn for 100 s, in one spatially scattered order per experiment that was identical at every rate and repeat. Rate was constant within a recording and ascended between recordings through 0.5 Hz, 1 Hz, 2 Hz, 4 Hz and 8 Hz before repeating (one pass through the five rates is a cycle), delivering 50 to 800 pulses per recording at the fixed recording duration. Acquisition ran for up to about three days (at most 73.5 h of recorded activity per experiment, excluding gaps between recordings), giving a median of 13 repeats per rate.

Analyses draw on nested subsets of the 14 ladder experiments, 12 of which are untreated controls (from 10 organoids). Drift trajectories (Figures 5B–5D) use all 12 (10 organoids). Site decoding and the subharmonic and subspace analyses (Figures 5F, S9 and 6) use the nine experiments with at least 20 spontaneous recordings, at least 20 stimulation recordings and at least five distinct rates; of those nine, one contributes only 22 stimulation recordings (4 to 5 repeats per rate, against 57 to 131 recordings for the other eight), so the decoder-decay analyses, which need three training and three held-out repeats at every rate, run on the remaining eight (8 organoids for the nine experiments, 7 for the decoder-decay analyses).

### Pretraining dataset and splits

The corpus comprises the recordings passing a floor of ≥ 30 active units (Table S1), split into training and holdout at the experiment level. A separate validation split, used throughout Figure 2 and Tables S5–S8, withheld 15% of the recordings of each multi-recording training experiment (30 organoids; 110 recordings under the 20-recording reference cap), so its experiments also contribute other recordings to training; all 30 are also training organoids.

### Model architecture and input representation

Per-neuron spike counts were binned at 10 ms, log-transformed (log(1 + *x*)), zero-padded to 493 units, and concatenated with an 8-dimensional stimulation feature vector and a 1-dimensional active unit count, giving a 502-dimensional input per bin. Learned spike-sorter source embeddings (Kilosort2 / RT-Sort) were summed with the input projections.

The model is a Mamba-2 state-space network^43^ with a causal strided convolution downsampler. The input is projected to *d*_model_ = 512, downsampled 2 × to 20 ms output resolution, and processed by 6 Mamba-2 layers (*d*_state_ = 128, ≈ 11.9 M parameters) followed by a residual MLP and a linear Poisson head predicting next-20 ms population rates.

### Pretraining objective and optimization

Context lengths are fixed in wall-clock time across binning configurations. Cached windows span 10 s. Truncated backpropagation through time operated on 5 s sub-chunks, the hidden state detached at each boundary and carried forward for up to 20 s before reset. The first ~250 ms of each carry segment was burn-in; all later time steps were weighted equally in the loss.

For output bin *t* the input spans input bins *t·s* through (*t*+1) *s* 1 at convolutional stride *s* = 2, and the target is the summed count over the next *s* input bins, so input and target never overlap and the forecast horizon is one output bin. The objective was the Poisson negative log-likelihood,

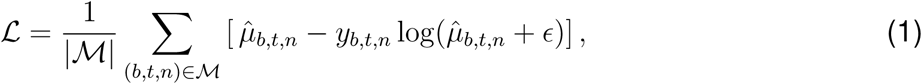

averaged over valid (batch, time, neuron) triples *M*, with *µ*^ the softplus-output rate, *y* the target count over the 20 ms window, and *ɛ* = 10*^−^*^4^.

Optimization used AdamW (learning rate 2 × 10*^−^*^4^, weight decay 0.01 on matrix parameters only) for 250 epochs at batch size 64, with 5-epoch linear warm-up, cosine annealing to zero and gradient-norm clipping at 1.0. A run seed (42, 43 or 44) fixed the validation split, shuffle order and chunk sampling but not weight initialization.

### Hardware and software environment

Training ran in full fp32 as single-GPU jobs on the National Research Platform (Ampere or newer, as the Mamba-2 fused kernels require compute capability ≥ 8.0), using Python 3.10, CUDA 12.4.1, PyTorch 2.5.1, mamba-ssm 2.3.0 and causal-conv1d 1.6.0, and retaining the best-validation-loss checkpoint. Downstream analyses ran in Python 3.11 with NumPy 2.4.6, SciPy 1.17.0^102^, scikit-learn 1.8.0, statsmodels 0.14.6, pandas 2.3.3, Matplotlib 3.10.8 and umap-learn 0.5.11^103^. All latent analyses (Figures 4–7) use one finalized checkpoint, the 6-layer, 10 ms-binning, seed-43 model trained on the full corpus with no per-experiment recording cap.

### Evaluation metrics

Next-window predictions were scored on the training, validation and holdout splits and reported in Tables S5 through S8. All models, whatever their input binning, are scored on the common 20 ms output grid; correlation metrics use mean-pooled rates and likelihood metrics summed counts.

The headline metric, **bits/spike above naive**, is the Poisson negative log-likelihood difference between the model and a constant-rate baseline, evaluated on the population trace and normalized to information per population spike. With *y_t_* the observed population count, 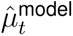 the predicted rate and *µ*^^naive^ the windowed mean,

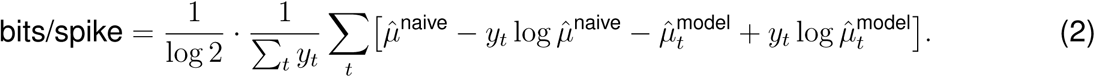

Positive values indicate that the model is more informative than that baseline.

The six prediction-accuracy metrics reported in the figures are bits/spike, Poisson skill and Pearson *r*, each computed for the population and per neuron. **Poisson skill** expresses the identical improvement as the fraction of the baseline’s Poisson deviance removed, so 0 matches the baseline and 1 is perfect. **Population** and **per-neuron Pearson** *r* correlate predicted against observed rates across time bins, pooled over units or computed per unit and averaged. **Per-neuron bits/spike** applies the same likelihood difference to each unit’s own spike train against that unit’s mean rate and reports the median over units that spiked; **per-neuron skill** is the median of the per-unit Poisson skill over all units. Three further metrics appear only in the supplemental tables. **Per-neuron MAE** and **MAE skill** report absolute error per unit per bin, the latter relative to a per-neuron mean-rate baseline. **Per-neuron Poisson NLL** is in nats per bin. The two calibration metrics are also recomputed over high-activity **event bins** alone, defined by 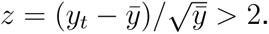

### Architecture and data ablations

#### Input binning

The reference configuration (10 ms bins, kernel and stride 2) was compared against 5 ms bins (kernel and stride 4) and 20 ms bins (no downsampling), each trained from scratch on three seeds with output rate, context, sequence length, burn-in, loss target, optimization and the Mamba-2 stack and MLP trunk held fixed; the undownsampled condition removes only the added downsampler. This grid was trained with a cap of 20 recordings per experiment rather than on the full corpus.

#### Model width

The width sweep (Figures 2B–2E, Table S7) retrained the architecture end to end at *d*_model_ ∊ {128, 256} with all else fixed.

#### Training-data capacity

The capacity sweep (a maximum of 1, 10, 20 or all recordings per experiment, against the 20-recording reference shared by the other axes) varied the maximum number of recordings supplied per experiment while holding the set of training experiments fixed, drawing recordings without replacement at a fixed seed where an experiment exceeded the cap. Every arm was scored on the same 30 validation organoids; the number of validation recordings per experiment varied with the cap.

#### Conditioning features

Each single-component ablation (Figures 2B–2E, Table S8) is a separately trained checkpoint with one conditioning input removed, not a masking of the deployed model at inference. The ablations removed the spike-sorter source embedding; the active-unit count from the input, the head’s conditioning, or both (three checkpoints); or all eight stimulation channels, the last scored only on experiments with evoked recordings (36 training organoids, 673 recordings; 28 validation organoids). All share the reference architecture, schedule and three-seed protocol.

### Causal exponentially weighted moving-average baseline

The causal EWMA baseline is an exponentially weighted moving average of each neuron’s own spike counts *y_t_* on the model’s 20 ms output grid, *s_t_* = *αy_t_* + (1 − *α*)*s_t−_*_1_, with *s_t−_*_1_ taken as the prediction for bin *t* so that, like the model, it uses only activity up to the preceding bin. It was scored with the same neurons, windows and metric code as the model (Table S2). For training and validation experiments it was scored on exactly the recordings, pretraining seeds and neurons behind Table S5 (86 training and 30 validation organoids), with a single smoothing factor chosen from *α* ∊ {0.005, 0.01, 0.02, 0.05, 0.1, 0.2, 0.4} (time constants of roughly 40 ms to 4 s) by pooled Poisson negative log-likelihood on training recordings and applied unchanged to validation. The slowest value, *α* = 0.005, was selected (Table S3), so the EWMA approaches each neuron’s recent mean rate. For held-out experiments, which are predicted by the model trained on all recordings per experiment, *α* was instead chosen per recording by Poisson negative log-likelihood on the 2-minute adaptation window, the filter was run causally from the start of that window, and predictions were scored on the evaluation window (see Two-minute linear-head fine-tuning).

### Two-minute linear-head fine-tuning

For the adaptation panels of Figure 2 the pretrained trunk of the all-recordings model was frozen and a per-neuron linear read-out fit on its 512-dimensional output as a Poisson generalized linear model, its ridge penalty chosen on held-out negative log-likelihood within the adaptation window. Within each recording the evaluation window was the final *E* = min(20, max(3, 0.30 *T*)) minutes for duration *T*, and the adaptation window the *K* minutes immediately preceding it, with *K* swept from 0.5 to 48 minutes and the neuron set held fixed; the main-text two-minute condition is one point on this ladder. The ladder was run once per pretraining seed and averaged across three seeds; Figure 2 reports the 9 held-out mouse cultures; the control condition refits the identical read-out on 11 training experiments.

Because the adaptation ladder scores a short window disjoint from its fit window whereas the design-axis sweeps (Tables S5–S7) score the whole recording, zero-shot values from the two protocols are not comparable and are reported on separate axes.

### Latent extraction for downstream analyses

Downstream analyses take the latent from the final hidden state of the sixth Mamba-2 layer plus the residual MLP trunk, immediately before the per-recording output head, giving a 512-dimensional vector every 20 ms. Model weights were frozen throughout unless stated.

Latents were extracted with the recurrent state threaded continuously across the whole recording rather than reset every 20 s as in training. The validation metrics (30 organoids) of Figure 2 and Tables S5–S7 are scored the same way, with the state carried across each recording.

### Human developmental clock and the spiking feature battery

#### Recording-level latent vector

Windows tile each recording contiguously with the state threaded across them. The 512-dimensional latent was standardized and projected onto its 64 leading principal components, fit once without labels on the latent pooled across all spontaneous recordings. The covariance of this projection across all timesteps was matrix-log transformed and vectorized (upper triangle including the diagonal, 2,080 values) into one summary vector per recording, used for both the developmental clock and the latent principal-component analysis (Figures 4A–4E).

#### Developmental clock

A ridge regression (*α* = 10) was fit from these vectors to chronological age on the CW20107 control line alone (*n* = 6 organoids, ages 113–163 d), with predictions leave-one-organoid-out (every experiment of the held-out organoid excluded) and the feature scaler refit inside each fold. The other three lines (14765-X2, *SHANK3*^+/+^, *SHANK3*^+/^*^−^*) were projected onto the fitted axis with no refitting, so their correlations with age are projections, not cross-validated by organoid. The organoids of a line were recorded repeatedly at the same ages, so ordering by age was assessed within organoid: for every organoid with at least three experiments, the Spearman *ρ* between predicted and chronological age, summarized by the mean and median over organoids and the number of organoids above zero, with a one-sided *P* for the mean from permuting age within each organoid 20,000 times (predictions held fixed). For the *SHANK3* lines, whose organoids were mostly recorded once, *ρ* was computed between organoid-mean predicted and chronological age, with exact permutation *P*. Plotted points in Figure 4C and E are organoid means; Figure 4B plots every recording. The offset of an experiment is its predicted minus chronological age. Each NPD line was compared with its own control by Cliff’s *δ* on experiment offsets, with *P* from exact enumeration of the ways to assign the line label to organoids; as a negative control, the identical pipeline, including its own permutation null, was rerun with sorted unit count as the only input. The axis serves as an ordinal maturation index, not a calibrated age estimate.

#### Spiking feature battery

Independently of the model latents, 28 classical spiking and network features in five families (rate and yield, network topology, bursting, criticality and temporal structure) were computed from the sorted spike trains of every corpus recording. The features were: rate and yield (8: population mean firing rate, standard deviation and skewness; per-unit rate skewness, kurtosis, coefficient of variation, Gini index and entropy); network topology (5: STTC graph density, modularity, normalized clustering coefficient, normalized path length and small-world index); bursting (5: burst event rate, mean burst duration, fraction of spikes in bursts, mean participation and participation variability); criticality (5: avalanche size and duration power-law exponents, branching ratio, and low- and high-frequency spectral slopes); and temporal structure (5: lag-1 autocorrelation, autocorrelation decay time, peak frequency, normalized total correlation and normalized Lempel-Ziv complexity).

#### Latent axis and feature correlations

Principal component analysis was applied to the latent covariance representation across 327 spontaneous recordings from mouse and human preparations, without genotype labels. These recordings included dorsal and ventral organoids, drug-treated cultures and monolayers. Each recording was projected onto the components through a leave-one-out refit that excludes it from the fit. Each of the 28 spiking features was related to PC2 by partial Spearman correlation across human recordings, controlling for log unit count, preparation and genotype (an NPD indicator), because PC2 itself separates genotypes. No PCA is applied to the feature battery itself.

#### Unit adjustment, effect sizes and convergence

Every genotype comparison regressed log unit count out within each preparation, with the slope fit on all recordings of that preparation regardless of line. Effect sizes are Cliff’s *δ* between each NPD line and its own control, computed on recordings; recordings were never pooled across preparations. A preparation’s latent displacement is the difference between its NPD and control means of the adjusted latent covariance vector, and convergence is the cosine similarity of the two displacements; the same statistic was computed on the two 28-feature effect profiles (Figure 4F). Both were tested against a null that permutes genotype labels across organoids within each preparation, by exact enumeration: 924 relabelings for the CW20107 versus 14765-X2 preparation, 462 for the *SHANK3* preparation, and all 426,888 combinations for the statistics that pool the two preparations (latent and feature convergence, and the joint PC2 test, which averages the two per-preparation *δ* under simultaneous relabelings). Recordings from one organoid always carry the same label. Per-preparation *P* values are two-sided and the convergence *P* values one-sided. As a sensitivity check, mixed-effects models with a random intercept per organoid gave the intraclass correlation and a Wald test of genotype.

The NPD predictability comparisons of Figure 3C were also repeated after regressing log unit count out of per-neuron Pearson *r* and bits/spike within each preparation, the same procedure used for the PC2 unit adjustment above, and the exact organoid-level permutation tests rerun on the residuals averaged within each organoid (fine-tuned 14765-X2 per-neuron Pearson *r P* = 0.0087 and bits/spike *P* = 0.026; no *SHANK3* comparison significant).

### Electrode-discrimination decoder analysis

Three analyses decode which electrode was stimulated from the single-trial population response (Figure S7; Figures 5F and S9), sharing the features, classifier and chance normalization below and differing in cohort, window and split.

Each trial was featurized as the mean response over a post-stimulus window minus the mean over the immediately preceding baseline window, canceling slow drift trial by trial: a 512-dimensional frozen-latent vector for the model arms, and the mean binned rate per unit for the raw firing-rate arm. The 0 ms edge is the first sample after the system’s stimulation blanking window. Identity was decoded with multiclass linear discriminant analysis (least-squares solver, Ledoit–Wolf shrinkage), and balanced accuracy normalized by the chance level 1*/C* for the *C* sites entering the fit (8 to 32; sites with fewer than six training trials dropped).

The model’s eight-channel stimulation input carries pulse presence, the scaled identities of up to five simultaneously stimulated electrodes, and the electrode count. ID-unmasked is the full model, ID-masked zeroes the five identity channels, and stim-masked zeroes all eight, so the ID-masked to stim-masked difference bounds the joint contribution of presence and count. All variants are evaluated on the same trials.

#### Causal sweep (Figure S7)

Forty-one causal sweep recordings from 17 organoids (31 recordings from 15 organoids in the pretraining corpus and 10 holdout recordings from 2 organoids) were decoded with all four arms using 0 to 120 ms post-stimulus and 60 to 0 ms baseline windows, scored by five-fold stratified cross-validation within each recording. A 42nd recording was excluded because only 51% of its pulses could be aligned to a detected stimulation artifact, with offsets up to 97 ms; every other recording aligned at least 96% of its pulses. As a drift control, the ID-masked decoder was also trained on each electrode’s early trials and tested on its late trials and the reverse, and compared against the raw firing-rate arm by two-sided Wilcoxon signed-rank test on per-organoid means (*n* = 17 organoids).

#### Frequency ladder (Figure S9A)

On the nine-experiment ladder cohort, latents were averaged over a 60 ms post-stimulus window and baseline-subtracted over the preceding 60 ms. Only the stim-masked variant was run, since the presence and count channels left intact by ID-masking would expose stimulation frequency. Frequency dependence was tested by paired two-sided Wilcoxon signed-rank test of each organoid’s 8 Hz against its 0.5 Hz accuracy-over-chance (the mean of its experiments’ values where an organoid has two experiments).

#### Cross-recording transfer (Figures 5E, 5F and S9)

On the same cohort, windows and stim-masked variant, a decoder fit on one stimulation recording was scored on another from the same experiment and frequency, over all ordered pairs sharing at least three sites; separation is the number of cycles between them. The zero-separation anchor is the median over twenty random half-splits of trials within one recording. Accuracy-over-chance was summarized as the median over frequencies and split directions within each experiment, averaged within each organoid, and then summarized across organoids. Because LDA whitening is estimated on the training recording, the analysis was repeated with a gain-invariant cosine nearest-template classifier, with an uncorrected readout lacking baseline subtraction, and with the unmasked arm (Figure S9).

#### Stimulation history (Figure S8)

Decoding was broken down by pulse position *k* within each causal-sweep block of 50 consecutive pulses on one electrode at 2 Hz. For each electrode, late pulses (*k* ≥ 11) were split into four folds, and a decoder trained on three folds scored the held-out fold and every earlier pulse, which it never saw. The cross-validated predictions of the main analyses were broken down the same way. On identical pulses, the latent was compared at each position with the firing rate in the single +0 ms bin and with the window-mean firing rate, by paired two-sided Wilcoxon signed-rank test on per-organoid means. Pull toward the previous block was the fraction of predictions naming the previous block’s electrode, relative to chance. Adaptation was assessed from the population spike count 0 to 10 ms after each pulse, relative to the recording’s late-pulse mean.

### Latent drift and pseudotime

Each recording was summarized by its latent centroid and covariance. Each latent dimension was first z-scored with the mean and standard deviation pooled over all of that organoid’s recordings. The centroid is the mean latent state over every time point of the recording. The covariance is the zero-lag covariance of the latent states themselves, not of their bin-to-bin differences, computed in the organoid’s 50 leading principal components (fit on 150 evenly spaced time points from each of its recordings). It was estimated from time points drawn at random within the recording, with the number of points matched across an organoid’s recordings (half the smallest recording’s sample, at most 4,000), so that recording length does not bias the comparison. Both pseudotimes used each experiment’s spontaneous recordings only. Location pseudotime is the projection of each recording’s centroid onto the first principal component of that experiment’s centroids, computed in the full 512-dimensional z-scored latent. Shape pseudotime is the first classical multidimensional-scaling coordinate of the matrix of squared Bures distances between trace-normalized covariances, after subtracting the organoid’s mean sampling floor (the distance between two random halves of one recording) and clipping at zero. The sign of each coordinate was set so that it increases with recording order. Each coordinate was compared with the true acquisition order by Spearman correlation, and each organoid’s value was the mean over its experiments. Drift exponents are the log–log slopes of mean-squared pairwise centroid displacement against the number of spontaneous recordings separating the pair, fit over the lags whose displacement exceeds a split-half sampling floor and after subtracting that floor. Here each centroid is the mean of the same matched random sample of time points used for the covariance, and the floor is the squared displacement between the means of two disjoint random halves of one recording, each of that size, averaged over the organoid’s recordings. The trajectories shown in Figure 5B are drawn, for display only, from smoothed and subsampled time points standardized within the spontaneous recordings; all reported statistics use the centroids defined above. Per-pulse template drift was decomposed into the component common to every stimulation electrode and the electrode-specific remainder; the population-wide and electrode-specific percentages quoted in the text are each readout’s fraction of between-cycle drift variance explained by the common component, computed separately for the uncorrected and per-pulse baseline-subtracted readouts.

### Subharmonic and subspace analyses

Each stimulation recording’s latent trajectory was projected onto the axis along which stimulation displaces the population and subsampled to 8,000 timepoints on the 20 ms latent grid. Power was estimated with a Lomb–Scargle periodogram; subharmonic strength is the peak power within 0.15 Hz of *f/*2 divided by the median power from 0.25 Hz to 12 Hz excluding the harmonic bands, and the reported ratio divides this value at *f/*2 by the larger of its values at two off-harmonic control frequencies (3.0 Hz and 5.5 Hz) in the same recording, tested against 1 across organoids (each organoid’s experiments averaged on the log scale) by two-sided Wilcoxon signed-rank test. For subspace overlap, every recording was centered on its own mean and projected onto the leading principal components holding 90% of the preceding spontaneous recording’s variance in the 512-dimensional latent; overlap is the fraction of the target recording’s variance falling inside that subspace. For a *k*-dimensional reference subspace the chance floor is *k/*512; the ceiling is a spontaneous recording scored against its own predecessor, and the distant comparison replaces the preceding reference with the experiment’s first spontaneous recording. The rate of spontaneous reorganization was measured separately with a fixed 50-dimensional subspace, against a split-half ceiling (each recording scored against its own other half) and a chance floor of 50*/*512. Each experiment’s values were averaged within its organoid before medians and tests.

For the surrogate control, each neuron’s binned firing rate in every stimulation recording was replaced by a memory-free surrogate, fit by least squares and Poisson-sampled: either a constant plus a single sinusoid at the stimulation frequency, or a constant plus sinusoids at up to eight integer multiples of that frequency, stopping below the Nyquist frequency. Spontaneous recordings were replaced by constant-rate Poisson trains at each neuron’s mean rate. Both surrogates repeat exactly once per stimulus period and so contain no energy at *f/*2. Each surrogate was passed through the same model with the same stimulation input, the stimulation axis was refit on its latents, and *f/*2 power was scored against the off-harmonic control frequencies exactly as for the recorded spikes (Figure S11A).

The same 8 Hz and 4 Hz comparison was repeated on the recorded firing rates binned on a 25 ms grid, on which 8 Hz spans exactly five bins per cycle, so no stimulus period spans a non-integer number of bins (a binning alias that can itself generate energy at *f/*2). The ratio of *f/*2 power in stimulation recordings to that in spontaneous recordings at 8 Hz was compared between the 20 ms and 25 ms grids, and 8 Hz was compared with 4 Hz on the alias-free grid, each by paired two-sided Wilcoxon signed-rank test (Figure S11C).

For the cross-organoid control, the same subspace projection was repeated with the reference subspace drawn from another organoid’s spontaneous recordings instead of the same organoid’s. Own-organoid and cross-organoid overlap, each against the same *k/*512 chance floor, were compared paired per organoid by two-sided Wilcoxon signed-rank test (Figure S12F).

### Balance-versus-fail decoder

Within closed-loop cartpole experiments run under reinforcement-learning teaching pulses, gameplay was segmented into episodes bounded by pole failure and labeled by balance duration: sustained-balance episodes lasted at least 30 s and quick-failure episodes less than 10 s, with the intervening band excluded. Within each episode the model’s 20 ms latents were Gaussian-smoothed (≈ 60 ms) and averaged within non-overlapping 2 s windows, giving one feature vector per window.

Balance and failure windows were matched by coarsened exact matching^83^ on mean absolute pole angle, its within-window standard deviation, and population firing rate, then subsampled to equal counts within each matched cell. A logistic-regression decoder (*ℓ*_2_, class-weight balanced, standardized features) was scored by area under the ROC curve under stratified group *k*-fold cross-validation grouped by episode. Significance came from a one-sided episode-level permutation test over 1,000 draws. Experiments with at least 30 windows of each class before matching and three balance episodes were retained. Across organoids, AUCs were averaged within an organoid and tested against 0.5 by one-sided Wilcoxon signed-rank test; an organoid counted as significant when the Fisher-combined permutation *P* of its experiments was below 0.05; the stars of Figures S14A, S14B and S14F mark the same Fisher-combined *P* and the plotted AUCs are the organoid means. The subset with at least 25 balance episodes is reported separately. One matched arm added time in the recording as a further coarsened-exact-matching key (Figure S14B); a second instead matched on time since episode start in 2 s bins, restricting balance windows to those starting within the first 6 s of an episode, the span that failure episodes cover (Figure S14F).

For the episode dimensionality analysis (Figures S14C and S14D), per-episode dynamics were pooled across three organoids that learned the task; effective dimensionality is the participation ratio of the smoothed latent trajectory projected onto that episode’s own top five principal components.

### Regime separation geometry

Regime separation was measured between the latent state distributions of three conditions recorded in the same experiment: spontaneous activity, open-loop stimulation, and closed-loop cartpole control. Each condition’s trajectory was summarized by its mean and covariance and each pair compared by the Gaussian Wasserstein-2 distance, whose square decomposes into a location term (squared centroid displacement), an orientation term (the Bures distance between trace-normalized covariances, as for shape pseudotime) and a size term (change in total latent variance). Because absolute distances are not comparable across experiments, each term is expressed as a multiple of a within-condition drift null computed between matched-duration halves of a single condition; the shape comparison (Figure S13B) reports the orientation term alone. Twenty-seven experiments, from 13 organoids, carry all three conditions. Each pair is reported per organoid, as the mean over its experiments (ratios averaged on the log scale), and the figures, summary values and tests all use these organoid values. The location term was also recomputed after regressing each recording’s population firing rate out of every latent dimension (Figure S13A). For the elapsed-time comparison, the same location term was measured between every pair of closed-loop recordings within an experiment and divided by the within-recording half-split null. Closed-loop recordings ran 15 min each with 45 min of rest, so recordings *k* apart were taken about *k* h apart. Each pair’s drift was expressed as a fraction of that organoid’s spontaneous-to-closed-loop location change and compared with it at each gap by a paired two-sided Wilcoxon signed-rank test on organoid means, Holm-corrected across gaps reached by at least five organoids (25 of the 27 experiments, 13 organoids, those with at least three closed-loop recordings; Figure S13F).

The subspace analyses used the 14 experiments (10 organoids) with at least three sustained-balance episodes, three failure episodes and 200 frames of spontaneous activity, projecting the decoder direction fit on the partly overlapping decoder cohort above (12 organoids). The capture curve projects each experiment’s decoder direction onto the leading principal components of its spontaneous, open-loop or pooled covariance and compares the cumulative captured fraction with the *k/*512 expected of a random direction, read out at *k* components holding 90% of that condition’s variance. Subspace overlap applies the same projection to whole conditions, reporting the fraction of one condition’s variance inside another’s leading subspace (open-loop in spontaneous, closed-loop in spontaneous, closed-loop in open-loop), each against its *k/*512 floor. Subspace overlap does not use the decoder direction and so used all 27 experiments (13 organoids). Because each had a single spontaneous recording, its ceiling was the fraction of the second half’s variance captured by the first half’s subspace, and open- and closed-loop overlap are reported as a percentage of that ceiling per organoid (Table S19). Two further nulls were computed for the balance-versus-failure direction (the difference between mean sustained-balance and mean failure states), in the latent and in binned firing rates. The first compares spontaneous variance along the direction with the same after shifting each spontaneous dimension independently in time, which keeps each dimension’s variance and removes their covariance (one-sided Wilcoxon signed-rank test on the log ratio of the organoids). The second ranks that variance among directions rebuilt from 300 random relabelings of the balance and failure episodes (two-sided Wilcoxon signed-rank test of the percentile against 50; Table S22).

### Closed-loop and open-loop axis alignment

The closed-loop axis, each experiment’s mean closed-loop latent state minus its mean spontaneous state, was compared by cosine similarity with the open-loop axis, the same difference for open-loop stimulation. Significance came from a permutation test that held each experiment’s own spontaneous mean fixed and permuted only which organoid’s open-loop mean was paired with it (never one from the same organoid), with the statistic the mean over organoids of each organoid’s mean cos, isolating each experiment’s own alignment from a population-wide axis shared by chance. The comparison was repeated after projecting each experiment’s own firing-rate axis, fit by ordinary least squares against population firing rate, out of both directions by Gram–Schmidt orthogonalization. Reliability was the cosine similarity between the closed-loop axis estimated independently from two disjoint halves of an experiment’s closed-loop recordings (Figures S13C and S13D).

## Quantification and statistical analysis

All tests are non-parametric and are stated with their exact *P* value, effect size and *n* in the text or figure legend. Unless stated otherwise, *n* denotes organoids. An experiment is one electrode configuration of one organoid, spike-sorted once, and experiments and recordings from the same organoid are repeated measures. All comparisons, including the model comparisons of Figure 2, are tested at the organoid level: permutation tests exchange labels between organoids, never between recordings of one organoid, and paired tests use the per-organoid mean of the per-experiment statistic, so the smallest attainable exact *P* follows the organoid count. The feature-profile comparison of Figure 4F uses the spiking feature as its unit. Model training seeds are technical replicates. Where a panel displays per-seed points but tests a per-experiment distribution, the test runs on seed-averaged values averaged within each organoid, and the quoted *n* is the number of organoids.

Paired comparisons, and one-sample comparisons of a paired difference against zero, used the Wilcoxon signed-rank test; independent groups, including cell lines, used Mann–Whitney *U*; monotonic associations used Spearman *ρ*; proportions were tested against 0.5 with an exact binomial test. Where several model variants were each compared with a common reference, *P* values were Holm-corrected within each design axis (input bin width, training data per experiment, model width, and the source-embedding and active-unit-count conditioning ablations), separately for each metric and cohort; the stimulation-input ablation, scored only on experiments with stimulation, was tested alone, metrics were never corrected across one another, and a comparison that stands alone was not corrected. The three masking-variant contrasts drawn in Figure S7 were Holm-corrected as one family, as were the three pairwise regime contrasts of Figure 7C, two of them one-sided (that the spontaneous–closed-loop shift exceeds each of the other two). Holm-adjusted values are reported as *P*_Holm_. Significance was defined as *P <* 0.05, applied to *P*_Holm_ wherever a family of tests was corrected. All statistics were computed in Python with SciPy and statsmodels (versions under Hardware and software environment and in the key resources table). Tests were two-sided, including all cell-line comparisons, model-variant comparisons and rank correlations, except those asking whether a quantity exceeds an explicit floor, which were one-sided: decoder AUC against a shuffled-label chance level, regime-separation ratios against their within-condition drift null of 1, subspace overlap and capture against their random-subspace chance floor, and the permutation tests other than the genotype and line comparisons, which were two-sided, whose *P* is by definition an upper-tail fraction.

Effect sizes accompany every test in the units of the comparison. Paired contrasts report the median across organoids of the paired differences. Cell-line comparisons report Cliff’s *δ*, and statistics pooled across the two preparations average the two per-preparation *δ*; the one exception is prediction accuracy by line (Figure 3C, Tables S11 and S12), which reports Cohen’s *d* computed on organoid means, with *P* from an exact permutation Mann-Whitney *U* test across organoids. Decoder performance is the median area under the ROC curve with its interquartile range, as a multiple of the matched chance level where one is defined. Correlational results report Spearman *ρ*. Interval estimates are 95% percentile bootstrap confidence intervals of the median over 5,000 resamples of organoids.

Permutation tests shuffle the label whose exchangeability is assumed, never the data as a whole; genotype and line labels were shuffled across organoids within preparation, so no permutation exchanges a sample between the two independent human preparations or between recordings of one organoid. Each analysis subsection states the label shuffled and the number of draws; the developmental clock used 20,000 draws (age permuted within each organoid); the genotype tests on the second principal component and the latent-axis alignment enumerated every relabeling of organoids exactly (924 for the CW20107 versus 14765-X2 preparation, 462 for *SHANK3*, and their product for the joint tests); and the closed-loop–open-loop direction alignment (Figure S13C) used 20,000. Empirical *P* values from random draws use the add-one estimator (1 + # null obs)*/*(1 + *B*), so the smallest attainable value is 1*/*(*B* + 1) and a *P* at that bound is resolution-limited rather than exactly zero; exact enumeration gives *P* = # null obs */N* over all *N* relabelings, so six organoids against six cannot reach a two-sided *P* below 2*/*924 = 0.0022. Every permutation, bootstrap and cross-validation split is generated from a fixed random seed, so all reported *P* values, intervals and fold assignments reproduce exactly. Every tested decoder and the developmental clock were fit under cross-validation respecting the dependence structure of the data (grouped by episode, leave-one-experiment-out, or, for the developmental clock, leave-one-organoid-out), and reported performance is always the held-out value.

No statistical method was used to predetermine sample size; every recording passing the ≥ 30-active-unit quality filter was included, and analysis cohorts follow the inclusion rules stated in each subsection, with one exception: a causal-sweep recording was dropped for failing the pulse-alignment criterion described under Causal sweep. Randomization was not applicable, as groups were defined by cell line, genotype, age or recorded condition rather than by an allocated treatment. Investigators were not blinded during data collection, and all recordings were processed by the same automated pipeline.

## SUPPLEMENTAL VIDEO TITLES AND LEGENDS

**Video S1. PULSE predicts population activity 20 ms ahead in a withheld validation recording, related to Figure 2**. Playback at 10× slower than real time over 9.7 s of recording (110.3 to 120.0 s; the time axis counts from 110 s, and the first 0.3 s precede the model’s first prediction). Mouse organoid at day 41 with 303 units; the organoid was in the training experiments, but this recording (about 10 min of spontaneous activity) was withheld from training. Black, ground-truth population rate (spike counts summed over all units in 20 ms bins) up to the present; orange, the all-recordings model’s prediction for the next 20 ms bin (shaded band), made only from activity before that bin, with the model state carried through the whole recording. Traces are not smoothed. Population Pearson *r* = 0.975 over the whole recording (copying the previous bin, *r* = 0.896). Once a burst has begun, PULSE predicts the jump to its peak in the next bin, but not the first rise out of quiet activity.

**Video S2. Spontaneous, open-loop and closed-loop activity of one organoid on its spontaneous map, related to Figure 7**. Playback at 20× speed; the first 282 s of each regime play in sync. Each regime’s latent states were centered on their own mean, removing the shift in average state (Figure 7C) so that only the directions of moment-to-moment variability remain. A three-dimensional UMAP was fit on the spontaneous states, and open-loop and closed-loop states were projected in to their nearest spontaneous neighbors. Left to right, spontaneous activity, open-loop stimulation and closed-loop cartpole control (grey, all three regimes; tint, that panel’s regime; dot and trail, the current state and the preceding 8 s). Bottom right, time balanced in the current cartpole episode.

**Video S3. Closed-loop cartpole control by the same organoid on its spontaneous map, related to Figure 7**. Playback at 20× real time over the full 867 s recording. Left, time balanced in each episode (top) and the simulated cart and pole (bottom). Middle, pole angle over time; dashed lines mark the 16*^◦^* failure threshold. Right, every closed-loop time point placed on the spontaneous map of Video S2 (grey, spontaneous states; orange, all closed-loop states; trail, the current episode, colored by time balanced).

## References

1. Velasco, S., Paulsen, B., and Arlotta, P. (2020). 3D brain organoids: Studying brain development and disease outside the embryo. Annu. Rev. Neurosci. 43, 375–389. doi: 10.1146/annurev-neuro-070918-050154.

2. Kelley, K.W., and Paşca, S.P. (2022). Human brain organogenesis: Toward a cellular understanding of development and disease. Cell 185, 42–61. doi: 10.1016/j.cell.2021.10.003.

3. Mostajo-Radji, M.A., Schmitz, M.T., Montoya, S.T., and Pollen, A.A. (2020). Reverse engineering human brain evolution using organoid models. Brain Res. 1729, 146582. doi: 10.1016/j.brainres.2019.146582.

4. Trujillo, C.A., Gao, R., Negraes, P.D., Gu, J., Buchanan, J., Preissl, S., Wang, A., Wu, W., Haddad, G.G., Chaim, I.A., et al. (2019). Complex oscillatory waves emerging from cortical organoids model early human brain network development. Cell Stem Cell 25, 558–569.e7. doi: 10.1016/j.stem.2019.08.002.

5. Hernandez, S., Schweiger, H.E., Cline, I., Kaurala, G.A., Robbins, A., Solis, D., Vera-Choqqueccota, S., Geng, J., van der Molen, T., Reyes, F., et al. (2026). Establishing mouse forebrain organoids as models of intrinsic cortical network assembly. Stem Cell Rep. 21, 102832. doi: 10.1016/j.stemcr.2026.102832.

6. van der Molen, T., Spaeth, A., Chini, M., Hernandez, S., Kaurala, G.A., Schweiger, H.E., Duncan, C., McKenna, S., Geng, J., Lim, M., et al. (2026). Preconfigured neuronal firing sequences in human brain organoids. Nat. Neurosci. 29, 123–135. doi: 10.1038/s41593-025-02111-0.

7. Sharf, T., van der Molen, T., Glasauer, S.M.K., Guzman, E., Buccino, A.P., Luna, G., Cheng, Z., Audouard, M., Ranasinghe, K.G., Kudo, K., et al. (2022). Functional neuronal circuitry and oscillatory dynamics in human brain organoids. Nat. Commun. 13, 4403. doi: 10.1038/s41467-022-32115-4.

8. Robbins, A., Schweiger, H.E., Hernandez, S., Spaeth, A., Voitiuk, K., Parks, D.F., van der Molen, T., Geng, J., Cline, I., Kosik, K.S., et al. (2026). Goal-directed learning in cortical organoids. Cell Rep. 45, 116984. doi: 10.1016/j.celrep.2026.116984.

9. Kagan, B.J., Kitchen, A.C., Tran, N.T., Habibollahi, F., Khajehnejad, M., Parker, B.J., Bhat, A., Rollo, B., Razi, A., and Friston, K.J. (2022). *In vitro* neurons learn and exhibit sentience when embodied in a simulated game-world. Neuron 110, 3952–3969.e8. doi: 10.1016/j.neuron.2022.09.001.

10. Cai, H., Ao, Z., Tian, C., Wu, Z., Liu, H., Tchieu, J., Gu, M., Mackie, K., and Guo, F. (2023). Brain organoid reservoir computing for artificial intelligence. Nat. Electron. 6, 1032–1039. doi: 10.1038/s41928-023-01069-w.

11. Mancinelli, S., Bariselli, S., and Lodato, S. (2025). The emergence of electrical activity in human brain organoids. Stem Cell Rep. 20, 102632. doi: 10.1016/j.stemcr.2025.102632.

12. Wen, S., Yin, A., Tseng, P.H., Itti, L., Lebedev, M.A., and Nicolelis, M. (2021). Capturing spike train temporal pattern with wavelet average coefficient for brain machine interface. Sci. Rep. 11, 19020. doi: 10.1038/s41598-021-98578-5.

13. Deneault, E., White, S.H., Rodrigues, D.C., Ross, P.J., Faheem, M., Zaslavsky, K., Wang, Z., Alexandrova, R., Pellecchia, G., Wei, W., et al. (2018). Complete disruption of autism-susceptibility genes by gene editing predominantly reduces functional connectivity of isogenic human neurons. Stem Cell Rep. 11, 1211–1225. doi: 10.1016/j.stemcr.2018.10.003.

14. Frega, M., Selten, M., Mossink, B., Keller, J.M., Linda, K., Moerschen, R., Qu, J., Koerner, P., Jansen, S., Oudakker, A., et al. (2020). Distinct pathogenic genes causing intellectual disability and autism exhibit a common neuronal network hyperactivity phenotype. Cell Rep. 30, 173–186.e6. doi: 10.1016/j.celrep.2019.12.002.

15. Iossifov, I., O’Roak, B.J., Sanders, S.J., Ronemus, M., Krumm, N., Levy, D., Stessman, H.A., Witherspoon, K.T., Vives, L., Patterson, K.E., et al. (2014). The contribution of de novo coding mutations to autism spectrum disorder. Nature 515, 216–221. doi: 10.1038/nature13908.

16. Satterstrom, F.K., Kosmicki, J.A., Wang, J., Breen, M.S., De Rubeis, S., An, J.Y., Peng, M., Collins, R., Grove, J., Klei, L., et al. (2020). Large-scale exome sequencing study implicates both developmental and functional changes in the neurobiology of autism. Cell 180, 568–584.e23. doi: 10.1016/j.cell.2019.12.036.

17. Gordon, A., Yoon, S.J., Bicks, L.K., Martín, J.M., Pintacuda, G., Arteaga, S., Wamsley, B., Guo, Q., Elahi, L., Dolmetsch, R.E., et al. (2026). Developmental convergence and divergence in human stem cell models of autism. Nature 651, 707–719. doi: 10.1038/s41586-025-10047-5.

18. Paulsen, B., Velasco, S., Kedaigle, A.J., Pigoni, M., Quadrato, G., Deo, A.J., Adiconis, X., Uzquiano, A., Sartore, R., Yang, S.M., et al. (2022). Autism genes converge on asynchronous development of shared neuron classes. Nature 602, 268–273. doi: 10.1038/s41586-021-04358-6.

19. Fernandez Garcia, M., Retallick-Townsley, K., Pruitt, A., Davidson, E.A., Balafkan, N., Warrell, J., Huang, T.C., Kibowen, A., Chu, Z., Dai, Y., et al. (2026). Transcriptomic and phenotypic convergence of neurodevelopmental disorder risk genes in vitro and in vivo. Nat. Neurosci. pp. 1079–1094. doi: 10.1038/s41593-026-02247-7.

20. Sundberg, M., Pinson, H., Smith, R.S., Winden, K.D., Venugopal, P., Tai, D.J.C., Gusella, J.F., Talkowski, M.E., Walsh, C.A., Tegmark, M., et al. (2021). 16p11.2 deletion is associated with hyperactivation of human iPSC-derived dopaminergic neuron networks and is rescued by RHOA inhibition in vitro. Nat. Commun. 12, 2897. doi: 10.1038/s41467-021-23113-z.

21. Edelman, G.M. (1987). Neural Darwinism: The Theory of Neuronal Group Selection. New York: Basic Books.

22. Kenet, T., Bibitchkov, D., Tsodyks, M., Grinvald, A., and Arieli, A. (2003). Spontaneously emerging cortical representations of visual attributes. Nature 425, 954–956. doi: 10.1038/nature02078.

23. Luczak, A., Bartó, P., and Harris, K.D. (2009). Spontaneous events outline the realm of possible sensory responses in neocortical populations. Neuron 62, 413–425. doi: 10.1016/j.neuron.2009.03.014.

24. Berkes, P., Orbán, G., Lengyel, M., and Fiser, J. (2011). Spontaneous cortical activity reveals hallmarks of an optimal internal model of the environment. Science 331, 83–87. doi: 10.1126/science.1195870.

25. Golshani, P., Gonçalves, J.T., Khoshkhoo, S., Mostany, R., Smirnakis, S., and Portera-Cailliau, C. (2009). Internally mediated developmental desynchronization of neocortical network activity. J. Neurosci. 29, 10890–10899. doi: 10.1523/JNEUROSCI.2012-09.2009.

26. Chini, M., Pfeffer, T., and Hanganu-Opatz, I. (2022). An increase of inhibition drives the developmental decorrelation of neural activity. eLife 11, e78811. doi: 10.7554/eLife.78811.

27. Antón-Bolaños, N., Sempere-Ferràndez, A., Guillamón-Vivancos, T., Martini, F.J., Pérez-Saiz, L., Gezelius, H., Filipchuk, A., Valdeolmillos, M., and López-Bendito, G. (2019). Prenatal activity from thalamic neurons governs the emergence of functional cortical maps in mice. Science 364, 987–990. doi: 10.1126/science.aav7617.

28. Antón-Bolaños, N., Espinosa, A., and López-Bendito, G. (2018). Developmental interactions between thalamus and cortex: a true love reciprocal story. Curr. Opin. Neurobiol. 52, 33–41. doi: 10.1016/j.conb.2018.04.018.

29. Martini, F.J., Guillamón-Vivancos, T., Moreno-Juan, V., Valdeolmillos, M., and López-Bendito, G. (2021). Spontaneous activity in developing thalamic and cortical sensory networks. Neuron 109, 2519–2534. doi: 10.1016/j.neuron.2021.06.026.

30. Aníbal-Martínez, M., Puche-Aroca, L., Pérez-Montoyo, E., Pumo, G., Madrigal, M.P., Rodríguez-Malmierca, L.M., Martini, F.J., Rijli, F.M., and López-Bendito, G. (2025). A pre-natal window for enhancing spatial resolution of cortical barrel maps. Nat. Commun. 16, 1955. doi: 10.1038/s41467-025-57052-w.

31. Jumper, J., Evans, R., Pritzel, A., Green, T., Figurnov, M., Ronneberger, O., Tunyasuvunakool, K., Bates, R., Žídek, A., Potapenko, A., et al. (2021). Highly accurate protein structure prediction with AlphaFold. Nature 596, 583–589. doi: 10.1038/s41586-021-03819-2.

32. Lin, Z., Akin, H., Rao, R., Hie, B., Zhu, Z., Lu, W., Smetanin, N., Verkuil, R., Kabeli, O., Shmueli, Y., et al. (2023). Evolutionary-scale prediction of atomic-level protein structure with a language model. Science 379, 1123–1130. doi: 10.1126/science.ade2574.

33. Nguyen, E., Poli, M., Durrant, M.G., Thomas, A.W., Kang, B., Sullivan, J., Ng, M.Y., Lewis, A., Patel, A., Lou, A., et al. (2024). Sequence modeling and design from molecular to genome scale with Evo. Science 386, eado9336. doi: 10.1126/science.ado9336.

34. Cui, H., Wang, C., Maan, H., Pang, K., Luo, F., Duan, N., and Wang, B. (2024). scGPT: toward building a foundation model for single-cell multi-omics using generative AI. Nat. Methods 21, 1470–1480. doi: 10.1038/s41592-024-02201-0.

35. Rosen, Y., Roohani, Y., Agrawal, A., Samotorčan, L., Tabula Sapiens Consortium, Quake, S.R., and Leskovec, J. (2026). Universal cell embedding provides a foundation model for cell biology. Nature 656, 183–191. doi: 10.1038/s41586-026-10689-z.

36. Azabou, M., Arora, V., Ganesh, V., Mao, X., Nachimuthu, S.B., Mendelson, M.J., Richards, B.A., Perich, M.G., Lajoie, G., and Dyer, E.L. (2023). A unified, scalable framework for neural population decoding. In Advances in Neural Information Processing Systems vol. 36. URL: https://openreview.net/forum?id=sw2Y0sirtM.

37. Ye, J., Collinger, J.L., Wehbe, L., and Gaunt, R. (2023). Neural Data Transformer 2: Multi-context pretraining for neural spiking activity. In Advances in Neural Information Processing Systems vol. 36. URL: https://openreview.net/forum?id=CBBtMnlTGq.

38. Ye, J., Rizzoglio, F., Ma, X., Smoulder, A., Mao, H., Blumenthal, G.H., Hockeimer, W., Kunigk, N.G., Moore, D.D., Marino, P.J., et al. (2025). A generalist intracortical motor decoder. In Advances in Neural Information Processing Systems vol. 38. URL: https://openreview.net/forum?id=utXSSdD9mt.

39. Zhang, Y., Wang, Y., Azabou, M., Andre, A., Wang, Z., Lyu, H., The International Brain Laboratory, Dyer, E., Paninski, L., and Hurwitz, C. (2025). Neural encoding and decoding at scale. Preprint at 10.48550/arXiv.2504.08201. .

40. Minnick, J.R., Geng, J., Hussain, K., Gonzalez-Ferrer, J., Robbins, A., Mostajo-Radji, M.A., Haussler, D., Eshraghian, J.K., and Teodorescu, M. (2026). SpikeProphecy: A large-scale benchmark for autoregressive neural population forecasting. Preprint at 10.48550/arXiv.2605.12992. .

41. Azabou, M., Pan, K.X., Arora, V., Knight, I.J., Dyer, E.L., and Richards, B.A. (2025). Multi-session, multi-task neural decoding from distinct cell-types and brain regions. In International Conference on Learning Representations. URL: https://mlanthology.org/iclr/2025/azabou2025iclr-multisession/.

42. Arora, V., Lachi, D., Knight, I.J., Azabou, M., Richards, B., Hurwitz, C.L., Siegle, J., and Dyer, E.L. (2025). Know thyself by knowing others: Learning neuron identity from population context. In Advances in Neural Information Processing Systems vol. 38. URL: 10.48550/arXiv.2512.01199.

43. Dao, T., and Gu, A. (2024). Transformers are SSMs: Generalized models and efficient algorithms through structured state space duality. In Proceedings of the 41st International Conference on Machine Learning vol. 235 of Proceedings of Machine Learning Research. PMLR pp. 10041–10071. URL: https://proceedings.mlr.press/v235/dao24a.html.

44. Ryoo, A.H.W., Krishna, N.H., Mao, X., Azabou, M., Dyer, E.L., Perich, M.G., and Lajoie, G. (2025). Generalizable, real-time neural decoding with hybrid state-space models. In Advances in Neural Information Processing Systems vol. 38. URL: https://openreview.net/forum?id=1i4wNFgHDd.

45. Minnick, J.R., Gonzalez-Ferrer, J., Hussain, K., Geng, J., Robbins, A., Mostajo-Radji, M.A., Haussler, D., Eshraghian, J., and Teodorescu, M. (2026). Implicit behavioral decoding from next-step spike forecasts at population scale. Preprint at 10.48550/arXiv.2605.12999. .

46. Skaggs, W.E., McNaughton, B.L., and Gothard, K.M. (1992). An information-theoretic approach to deciphering the hippocampal code. In Advances in Neural Information Processing Systems vol. 5. Morgan-Kaufmann. URL: https://proceedings.neurips.cc/paperfiles/paper/1992/file/5dd9db5e033da9c6fb5ba83c7a7ebea9-Paper.pdf.

47. Pillow, J.W., Shlens, J., Paninski, L., Sher, A., Litke, A.M., Chichilnisky, E.J., and Simoncelli, E.P. (2008). Spatio-temporal correlations and visual signalling in a complete neuronal population. Nature 454, 995–999. doi: 10.1038/nature07140.

48. Cunningham, J.P., and Yu, B.M. (2014). Dimensionality reduction for large-scale neural recordings. Nat. Neurosci. 17, 1500–1509. doi: 10.1038/nn.3776.

49. Hyndman, R.J., and Athanasopoulos, G. (2021). Forecasting: Principles and Practice. 3rd ed.. Melbourne, Australia: OTexts. URL: https://otexts.com/fpp3/.

50. Lueckmann, J.M., Immer, A., Chen, A.B.Y., Li, P.H., Petkova, M.D., Iyer, N.A., Hesselink, L.W., Dev, A., Ihrke, G., Park, W., et al. (2025). ZAPBench: A benchmark for whole-brain activity prediction in zebrafish. In International Conference on Learning Representations (ICLR). URL: https://openreview.net/forum?id=oCHsDpyawq arXiv:2503.02618.

51. Wessberg, J., Stambaugh, C.R., Kralik, J.D., Beck, P.D., Laubach, M., Chapin, J.K., Kim, J., Biggs, S.J., Srinivasan, M.A., and Nicolelis, M.A.L. (2000). Real-time prediction of hand trajectory by ensembles of cortical neurons in primates. Nature 408, 361–365. doi: 10.1038/35042582.

52. Truccolo, W., Eden, U.T., Fellows, M.R., Donoghue, J.P., and Brown, E.N. (2005). A point process framework for relating neural spiking activity to spiking history, neural ensemble, and extrinsic covariate effects. J. Neurophysiol. 93, 1074–1089. doi: 10.1152/jn.00697.2004.

53. Cunningham, J.P., Gilja, V., Ryu, S.I., and Shenoy, K.V. (2009). Methods for estimating neural firing rates, and their application to brain-machine interfaces. Neural Netw. 22, 1235–1246. doi: 10.1016/j.neunet.2009.02.004.

54. Pochinok, I., Hanganu-Opatz, I.L., and Chini, M. (2026). iSTTC: A robust method for accurate estimation of intrinsic neural timescales from single-unit recordings. PLoS Comput. Biol. 22, e1013385. doi: 10.1371/journal.pcbi.1013385.

55. Deshpande, A., Yadav, S., Dao, D.Q., Wu, Z.Y., Hokanson, K.C., Cahill, M.K., Wiita, A.P., Jan, Y.N., Ullian, E.M., and Weiss, L.A. (2017). Cellular phenotypes in human iPSC-derived neurons from a genetic model of autism spectrum disorder. Cell Rep. 21, 2678–2687. doi: 10.1016/j.celrep.2017.11.037.

56. Zhang, H., McCarroll, A., Peyton, L., de León-Guerrerro, S.D., Zhang, S., Gowda, P., Sirkin, D., ElAchwah, M., Duhe, A., Wood, W.G., et al. (2024). Scaled and efficient derivation of loss-of-function alleles in risk genes for neurodevelopmental and psychiatric disorders in human iPSCs. Stem Cell Rep. 19, 1489–1504. doi: 10.1016/j.stemcr.2024.08.003.

57. Pollen, A.A., Bhaduri, A., Andrews, M.G., Nowakowski, T.J., Meyerson, O.S., Mostajo-Radji, M.A., Di Lullo, E., Alvarado, B., Bedolli, M., Dougherty, M.L., et al. (2019). Establishing cerebral organoids as models of human-specific brain evolution. Cell 176, 743–756.e17. doi: 10.1016/j.cell.2019.01.017.

58. Wells, M.F., Nemesh, J., Ghosh, S., Mitchell, J.M., Salick, M.R., Mello, C.J., Meyer, D., Pietilainen, O., Piccioni, F., Guss, E.J., et al. (2023). Natural variation in gene expression and viral susceptibility revealed by neural progenitor cell villages. Cell Stem Cell 30, 312–332.e13. doi: 10.1016/j.stem.2023.01.010.

59. Amin, N.D., Kelley, K.W., Kaganovsky, K., Onesto, M., Hao, J., Miura, Y., McQueen, J.P., Reis, N., Narazaki, G., Li, T., et al. (2024). Generating human neural diversity with a multiplexed morphogen screen in organoids. Cell Stem Cell 31, 1831–1846.e9. doi: 10.1016/j.stem.2024.10.016.

60. He, Z., Dony, L., Fleck, J.S., Szałata, A., Li, K.X., Slišković, I., Lin, H.C., Santel, M., Atamian, A., Quadrato, G., et al. (2024). An integrated transcriptomic cell atlas of human neural organoids. Nature 635, 690–698. doi: 10.1038/s41586-024-08172-8.

61. Wang, Y., Chiola, S., Yang, G., Russell, C., Armstrong, C.J., Wu, Y., Spampanato, J., Tarboton, P., Ullah, H.M.A., Edgar, N.U., et al. (2022). Modeling human telencephalic development and autism-associated SHANK3 deficiency using organoids generated from single neural rosettes. Nat. Commun. 13, 5688. doi: 10.1038/s41467-022-33364-z.

62. Schafer, S.T., Paquola, A.C.M., Stern, S., Gosselin, D., Ku, M., Pena, M., Kuret, T.J.M., Liyanage, M., Mansour, A.A., Jaeger, B.N., et al. (2019). Pathological priming causes developmental gene network heterochronicity in autistic subject-derived neurons. Nat. Neurosci. 22, 243–255. doi: 10.1038/s41593-018-0295-x.

63. Urresti, J., Zhang, P., Moran-Losada, P., Yu, N.K., Negraes, P.D., Trujillo, C.A., Antaki, D., Amar, M., Chau, K., Pramod, A.B., et al. (2021). Cortical organoids model early brain development disrupted by 16p11.2 copy number variants in autism. Mol. Psychiatry 26, 7560–7580. doi: 10.1038/s41380-021-01243-6.

64. Villa, C.E., Cheroni, C., Dotter, C.P., López-Tóbon, A., Oliveira, B., Sacco, R., Yahya, A.Ç., Morandell, J., Gabriele, M., Tavakoli, M.R., et al. (2022). CHD8 haploinsufficiency links autism to transient alterations in excitatory and inhibitory trajectories. Cell Rep. 39, 110615. doi: 10.1016/j.celrep.2022.110615.

65. Rao, S.B., Sun, Z., Brundu, F., Chen, Y., Sun, Y., Zhu, H., Shprintzen, R.J., Tomer, R., Rabadan, R., Leong, K.W., et al. (2025). Aberrant pace of cortical neuron development in brain organoids from patients with 22q11.2 deletion syndrome-associated schizophrenia. Nat. Commun. 16, 6986. doi: 10.1038/s41467-025-62187-x.

66. Sebastian, R., Jin, K., Pavon, N., Bansal, R., Potter, A., Song, Y., Babu, J., Gabriel, R., Sun, Y., Aronow, B., et al. (2023). Schizophrenia-associated nrxn1 deletions induce developmental-timing-and cell-type-specific vulnerabilities in human brain organoids. Nat. Commun. 14, 3770. doi: 10.1038/s41467-023-39420-6.

67. Yoon, S.J., Elahi, L.S., Paşca, A.M., Marton, R.M., Gordon, A., Revah, O., Miura, Y., Walczak, E.M., Holdgate, G.M., Fan, H.C., et al. (2019). Reliability of human cortical organoid generation. Nat. Methods 16, 75–78. doi: 10.1038/s41592-018-0255-0.

68. Velasco, S., Kedaigle, A.J., Simmons, S.K., Nash, A., Rocha, M., Quadrato, G., Paulsen, B., Nguyen, L., Adiconis, X., Regev, A., et al. (2019). Individual brain organoids reproducibly form cell diversity of the human cerebral cortex. Nature 570, 523–527. doi: 10.1038/s41586-019-1289-x.

69. Maass, W., Natschläger, T., and Markram, H. (2002). Real-time computing without stable states: a new framework for neural computation based on perturbations. Neural Comput. 14, 2531–2560. doi: 10.1162/089976602760407955.

70. Buzsáki, G. (2010). Neural syntax: cell assemblies, synapsembles, and readers. Neuron 68, 362–385. doi: 10.1016/j.neuron.2010.09.023.

71. Ziv, Y., Burns, L.D., Cocker, E.D., Hamel, E.O., Ghosh, K.K., Kitch, L.J., El Gamal, A., and Schnitzer, M.J. (2013). Long-term dynamics of CA1 hippocampal place codes. Nat. Neurosci. 16, 264–266. doi: 10.1038/nn.3329.

72. Driscoll, L.N., Pettit, N.L., Minderer, M., Chettih, S.N., and Harvey, C.D. (2017). Dynamic reorganization of neuronal activity patterns in parietal cortex. Cell 170, 986–999.e16. doi: 10.1016/j.cell.2017.07.021.

73. Deitch, D., Rubin, A., and Ziv, Y. (2021). Representational drift in the mouse visual cortex. Current Biology 31, 4327–4339.e6. doi: 10.1016/j.cub.2021.07.062.

74. Schoonover, C.E., Ohashi, S.N., Axel, R., and Fink, A.J.P. (2021). Representational drift in primary olfactory cortex. Nature 594, 541–546. doi: 10.1038/s41586-021-03628-7.

75. Marks, T.D., and Goard, M.J. (2021). Stimulus-dependent representational drift in primary visual cortex. Nat. Commun. 12, 5169. doi: 10.1038/s41467-021-25436-3.

76. Stringer, C., Pachitariu, M., Steinmetz, N., Reddy, C.B., Carandini, M., and Harris, K.D. (2019). Spontaneous behaviors drive multidimensional, brainwide activity. Science 364, eaav7893. doi: 10.1126/science.aav7893.

77. Musall, S., Kaufman, M.T., Juavinett, A.L., Gluf, S., and Churchland, A.K. (2019). Single-trial neural dynamics are dominated by richly varied movements. Nat. Neurosci. 22, 1677–1686. doi: 10.1038/s41593-019-0502-4.

78. Steinmetz, N.A., Aydin, C., Lebedeva, A., Okun, M., Pachitariu, M., Bauza, M., Beau, M., Bhagat, J., Böhm, C., Broux, M., et al. (2021). Neuropixels 2.0: A miniaturized high-density probe for stable, long-term brain recordings. Science 372, eabf4588. doi: 10.1126/science.abf4588.

79. Trapnell, C., Cacchiarelli, D., Grimsby, J., Pokharel, P., Li, S., Morse, M., Lennon, N.J., Livak, K.J., Mikkelsen, T.S., and Rinn, J.L. (2014). The dynamics and regulators of cell fate decisions are revealed by pseudotemporal ordering of single cells. Nat. Biotechnol. 32, 381–386. doi: 10.1038/nbt.2859.

80. Scargle, J.D. (1982). Studies in astronomical time series analysis. II. statistical aspects of spectral analysis of unevenly spaced data. Astrophys. J. 263, 835–853. doi: 10.1086/160554.

81. Ruf, T. (1999). The Lomb-Scargle periodogram in biological rhythm research: analysis of incomplete and unequally spaced time-series. Biol. Rhythm Res. 30, 178–201. doi: 10.1076/brhm.30.2.178.1422.

82. Herrmann, C.S., Murray, M.M., Ionta, S., Hutt, A., and Lefebvre, J. (2016). Shaping intrinsic neural oscillations with periodic stimulation. J. Neurosci. 36, 5328–5337. doi: 10.1523/JNEUROSCI.0236-16.2016.

83. Iacus, S.M., King, G., and Porro, G. (2012). Causal inference without balance checking: Coarsened exact matching. Polit. Anal. 20, 1–24. doi: 10.1093/pan/mpr013.

84. Emani, P.S., Liu, J.J., Clarke, D., et al. (2024). Single-cell genomics and regulatory networks for 388 human brains. Science 384, eadi5199. doi: 10.1126/science.adi5199.

85. BRAIN Initiative Cell Census Network (BICCN) (2021). A multimodal cell census and atlas of the mammalian primary motor cortex. Nature 598, 86–102. doi: 10.1038/s41586-021-03950-0.

86. Rubenstein, J.L.R., and Merzenich, M.M. (2003). Model of autism: increased ratio of excitation/inhibition in key neural systems. Genes Brain Behav. 2, 255–267. doi: 10.1034/j.1601-183X.2003.00037.x.

87. Tanveer, M.S., Mostajo-Radji, M.A., and Wang, G. (2026). A discrete generative model of neuronal spiking activity on microelectrode arrays. Preprint at 10.48550/arXiv.2609.23907. .

88. Park, Y., Franz, C.K., Ryu, H., Luan, H., Cotton, K.Y., Kim, J.U., Chung, T.S., Zhao, S., Vazquez-Guardado, A., Yang, D.S., et al. (2021). Three-dimensional, multifunctional neural interfaces for cortical spheroids and engineered assembloids. Sci. Adv. 7, eabf9153. doi: 10.1126/sciadv.abf9153.

89. Yang, X., Forró, C., Li, T.L., Miura, Y., Zaluska, T.J., Tsai, C.T., Kanton, S., McQueen, J.P., Chen, X., Mollo, V., et al. (2024). Kirigami electronics for long-term electrophysiological recording of human neural organoids and assembloids. Nat. Biotechnol. 42, 1836–1843. doi: 10.1038/s41587-023-02081-3.

90. de Vries, S.E., Siegle, J.H., and Koch, C. (2023). Sharing neurophysiology data from the Allen Brain Observatory. eLife 12, e85550. doi: 10.7554/eLife.85550.

91. Ramos, D.M., Skarnes, W.C., Singleton, A.B., Cookson, M.R., and Ward, M.E. (2021). Tackling neurodegenerative diseases with genomic engineering: A new stem cell initiative from the NIH. Neuron 109, 1080–1083. doi: 10.1016/j.neuron.2021.03.022.

92. Lin, S.S., DeLaura, S., and Jones, E.M. (2020). The CIRM iPSC repository. Stem Cell Res. 44, 101671. doi: 10.1016/j.scr.2019.101671.

93. Nguyen, C.V., Martija, A., Jaklic, D.C., White, R., Yang, M.G., Nano, P.R., Soto, J.A., Mil, J., Rana, D., Martin, J.M., et al. (2026). Thalamic NRXN1-mediated input to human cortical progenitors drives excitatory neurogenesis. Science 393, eady5180. doi: 10.1126/science.ady5180.

94. Tsyporin, J., Tastad, D., Ma, X., Nehme, A., Finn, T., Huebner, L., Liu, G., Gallardo, D., Makhamreh, A., Roberts, J.M., et al. (2021). Transcriptional repression by FEZF2 restricts alternative identities of cortical projection neurons. Cell Rep. 35, 109269. doi: 10.1016/j.celrep.2021.109269.

95. Chen, S., Zhou, Y., Chen, Y., and Gu, J. (2018). fastp: an ultra-fast all-in-one FASTQ preprocessor. Bioinformatics 34, i884–i890. doi: 10.1093/bioinformatics/bty560.

96. Dobin, A., Davis, C.A., Schlesinger, F., Drenkow, J., Zaleski, C., Jha, S., Batut, P., Chaisson, M., and Gingeras, T.R. (2013). STAR: ultrafast universal RNA-seq aligner. Bioinformatics 29, 15–21. doi: 10.1093/bioinformatics/bts635.

97. Liu, D. (2019). Algorithms for efficiently collapsing reads with unique molecular identifiers. PeerJ 7, e8275. doi: 10.7717/peerj.8275.

98. Liao, Y., Smyth, G.K., and Shi, W. (2014). featurecounts: an efficient general purpose program for assigning sequence reads to genomic features. Bioinformatics 30, 923–930. doi: 10.1093/bioinformatics/btt656.

99. Pachitariu, M., Steinmetz, N.A., Kadir, S.N., Carandini, M., and Harris, K.D. (2016). Fast and accurate spike sorting of high-channel count probes with Kilo-Sort. In Advances in Neural Information Processing Systems vol. 29. Curran Associates, Inc. URL: https://proceedings.neurips.cc/paperfiles/paper/2016/file/1145a30ff80745b56fb0cecf65305017-Paper.pdf.

100. Hill, D.N., Mehta, S.B., and Kleinfeld, D. (2011). Quality metrics to accompany spike sorting of extracellular signals. J. Neurosci. 31, 8699–8705. doi: 10.1523/JNEUROSCI.0971-11.2011.

101. van der Molen, T., Lim, M., Bartram, J., Cheng, Z., Robbins, A., Parks, D.F., Petzold, L.R., Hierlemann, A., Haussler, D., Hansma, P.K., et al. (2024). RT-Sort: An action potential propagation-based algorithm for real time spike detection and sorting with millisecond latencies. PLoS One 19, e0312438. doi: 10.1371/journal.pone.0312438.

102. Virtanen, P., Gommers, R., Oliphant, T.E., Haberland, M., Reddy, T., Cournapeau, D., Burovski, E., Peterson, P., Weckesser, W., Bright, J., et al. (2020). SciPy 1.0: Fundamental algorithms for scientific computing in Python. Nat. Methods 17, 261–272. doi: 10.1038/s41592-019-0686-2.

103. McInnes, L., Healy, J., Saul, N., and Großberger, L. (2018). UMAP: Uniform manifold approximation and projection. J. Open Source Softw. 3, 861. doi: 10.21105/joss.00861.

