## Supplementary Figures and Tables for "A foundation model for brain organoid network dynamics uncovers intrinsic computation and convergent disease phenotypes"

#### Supplemental figures

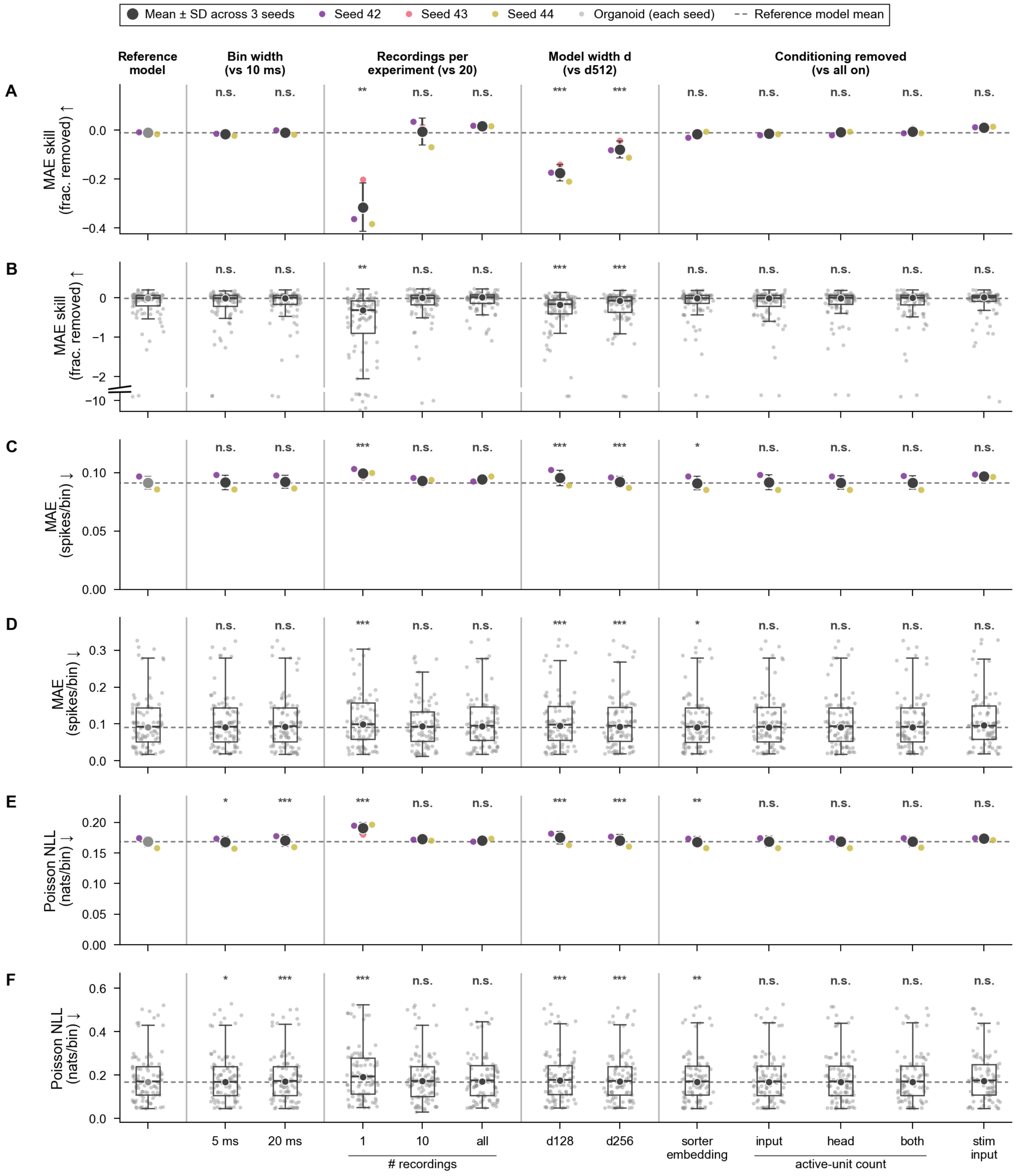

Mean ± SD across 3 seeds
Seed 42
Seed 43
Seed 44
Organoid (each seed)
Reference model mean
Reference model
Bin width (vs 10 ms)
Recordings per experiment (vs 20)
Model width d (vs d512)
Conditioning removed (vs all on)
A
B
C
D
E
F
MAE skill (frac. removed) ↑
MAE (spikes/bin) ↓
Poisson NLL (nats/bin) ↓
n.s.
*
**
***
5 ms
20 ms
1
10
all
### recordings
d128
d256
sorter embedding
input
head
both
active-unit count
stim input

Figure S1: **Per-neuron absolute-error metrics do not resolve model improvements visible in correlation and information-theoretic families, related to Figure 2.**
(A and B) Per-neuron MAE skill score (0, baseline; 1, perfect; higher is better), shown as a seed-level summary (A) above its organoid-level distribution (B).
(C and D) Per-neuron mean absolute error (MAE; spikes/bin, lower is better), shown as a seed-level summary (C) above its organoid-level distribution (D).
(E and F) Per-neuron Poisson negative log-likelihood (nats/bin, lower is better), shown as a seed-level summary (E) above its organoid-level distribution (F).
In (A, C, and E), mean $\pm$ 1 SD across 3 random seeds on the validation organoids ($n = 30$); in (B, D, and F), the organoid-level distribution of the same metric (one point per organoid, experiments averaged within each organoid). Per-cell significance is a paired two-sided Wilcoxon signed-rank test across organoids on organoid means of seed-averaged per-experiment differences from the reference model, Holm-corrected within each design axis and metric, with the stimulation-input ablation tested alone on the $n = 28$ organoids with stimulation. n.s., $P \geq 0.05$; $^{*}P < 0.05$; $^{**}P < 0.01$; $^{***}P < 0.001$.

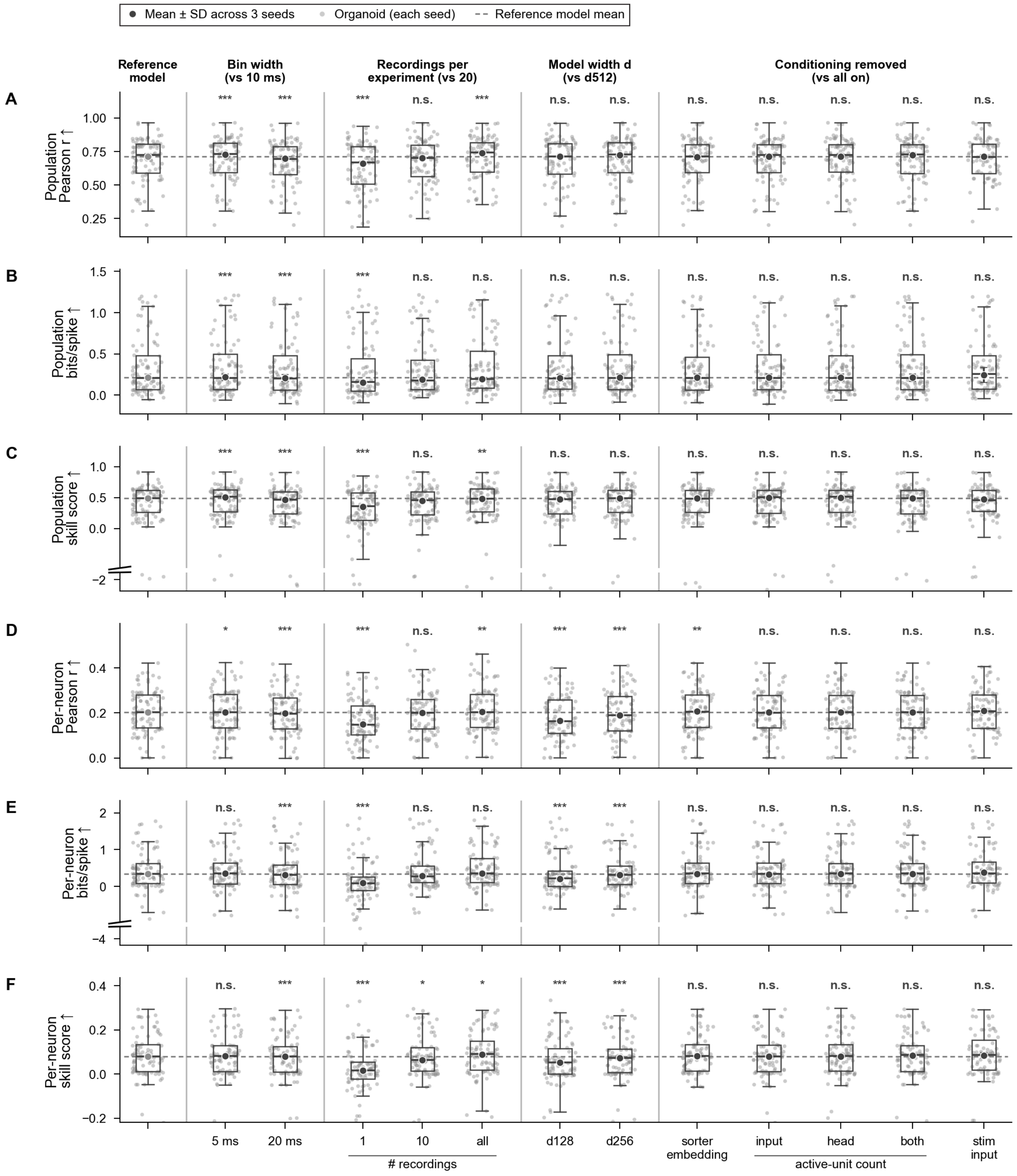

Mean ± SD across 3 seeds
Organoid (each seed)
Reference model mean
Reference model
Bin width (vs 10 ms)
Recordings per experiment (vs 20)
Model width d (vs d512)
Conditioning removed (vs all on)
A
Population Pearson r ↑
B
Population bits/spike ↑
C
Population skill score ↑
D
Per-neuron Pearson r ↑
E
Per-neuron bits/spike ↑
F
Per-neuron skill score ↑
5 ms
20 ms
1
10
all
### recordings
d128
d256
sorter embedding
input
head
both
active-unit count
stim input

Figure S2: **Organoid-level metric distributions are consistent across model design axes, related to Figure 2.**
(A–F) Organoid-level distributions for the validation split ($n = 30$ organoids; one point per organoid) of population Pearson $r$ (A), population bits/spike (B), population skill score (C), per-neuron Pearson $r$ (D), per-neuron bits/spike (E) and per-neuron skill score (F), plotted as box-and-strip distributions across the reference model, the input-binning sweep (5 and 20 ms vs 10 ms; evaluated on a uniform 20 ms forward prediction), the training-data-capacity sweep (cap1, cap10, and capall vs cap20), the model-width sweep ($d128$, $d256$ vs $d512$; $d_{\mathrm{model}} \in \{128, 256, 512\}$), and single-component conditioning-feature ablations against the all-on reference (the spike-sorting source embedding, the input/head active-unit-count channels, and the stimulation input each removed).
Box plots: center line, median; box, IQR; whiskers, 1.5× IQR. Error bars (black): mean ± 1 SD across 3 random seeds. Per-cell significance is a paired two-sided Wilcoxon signed-rank test across organoids on organoid means of seed-averaged per-experiment differences from the reference model, Holm-corrected within each design axis and metric, with the stimulation-input ablation tested alone on the $n = 28$ organoids with stimulation. n.s., $P \geq 0.05$; $^{*}P < 0.05$; $^{**}P < 0.01$; $^{***}P < 0.001$.

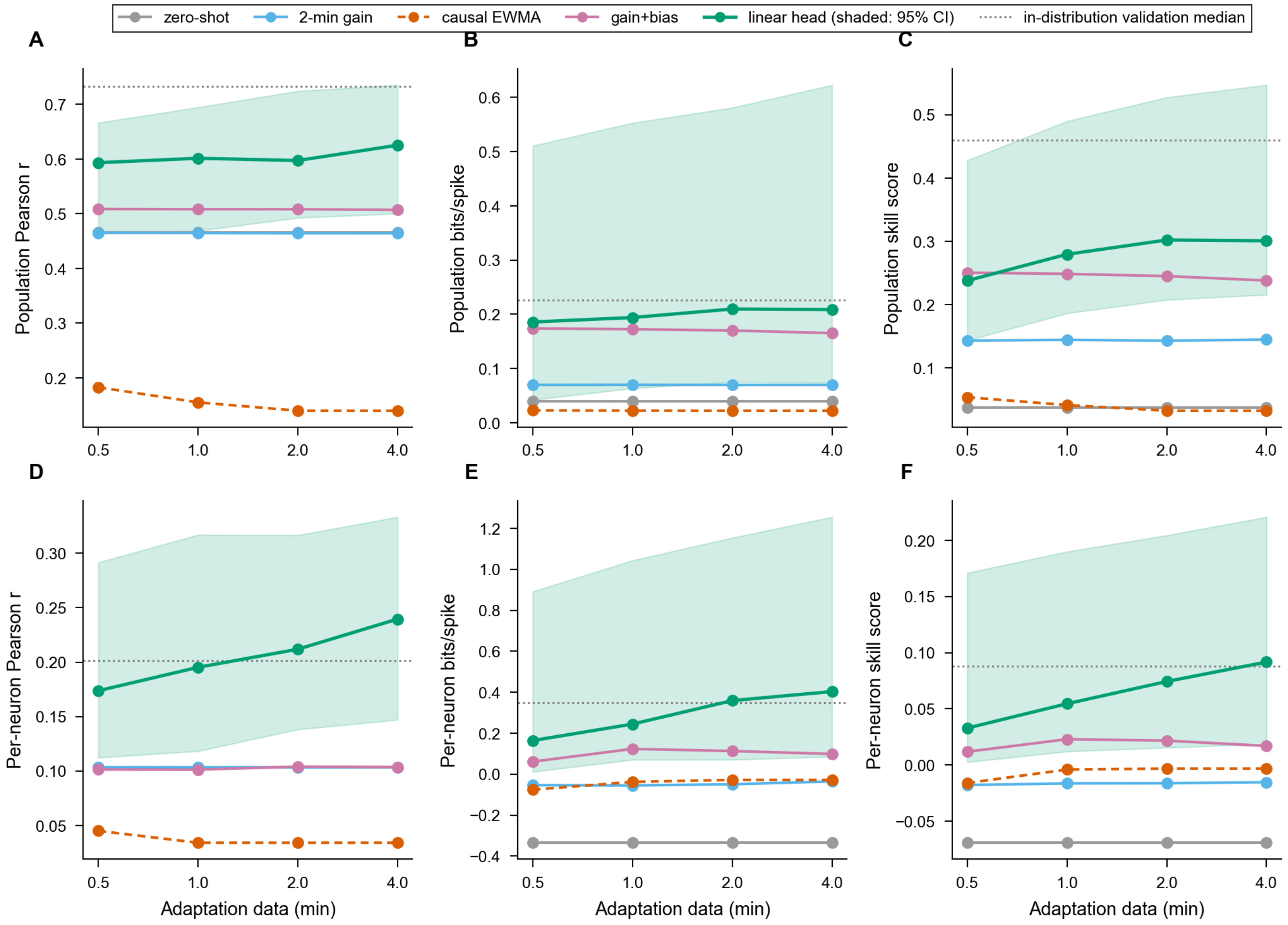


Figure S3: **Two minutes of linear-head fine-tuning rescues rate calibration on held-out experiments, related to Figure 2.**

(A) Predictive recovery on holdout cultures (mouse only, $n = 9$ cultures, six of them 2D monolayers) during linear head adaptation over 0.5 to 4.0 minutes for population Pearson correlation relative to zero-shot performance and causal EWMA reference.

(B) Same as (A), but for population bits per spike.

(C) Same as (A), but for population Poisson skill score.

(D) Same as (A), but for per-neuron Pearson correlation.

(E) Same as (A), but for per-neuron bits per spike.

(F) Same as (A), but for per-neuron Poisson skill score.

Shaded bands, 95% bootstrap CI of the median (5,000 iterations). "2-min gain" is a per-neuron multiplicative rate correction fit on the adaptation window alone; "gain+bias" adds a per-neuron log-linear offset, fit by Poisson regression on the same window; both are refit at each adaptation budget and scored on the same held-out evaluation window. The dashed "in-dist val median" line is zero-shot performance on the disjoint in-distribution validation cohort ($n = 30$ organoids), shown for reference.

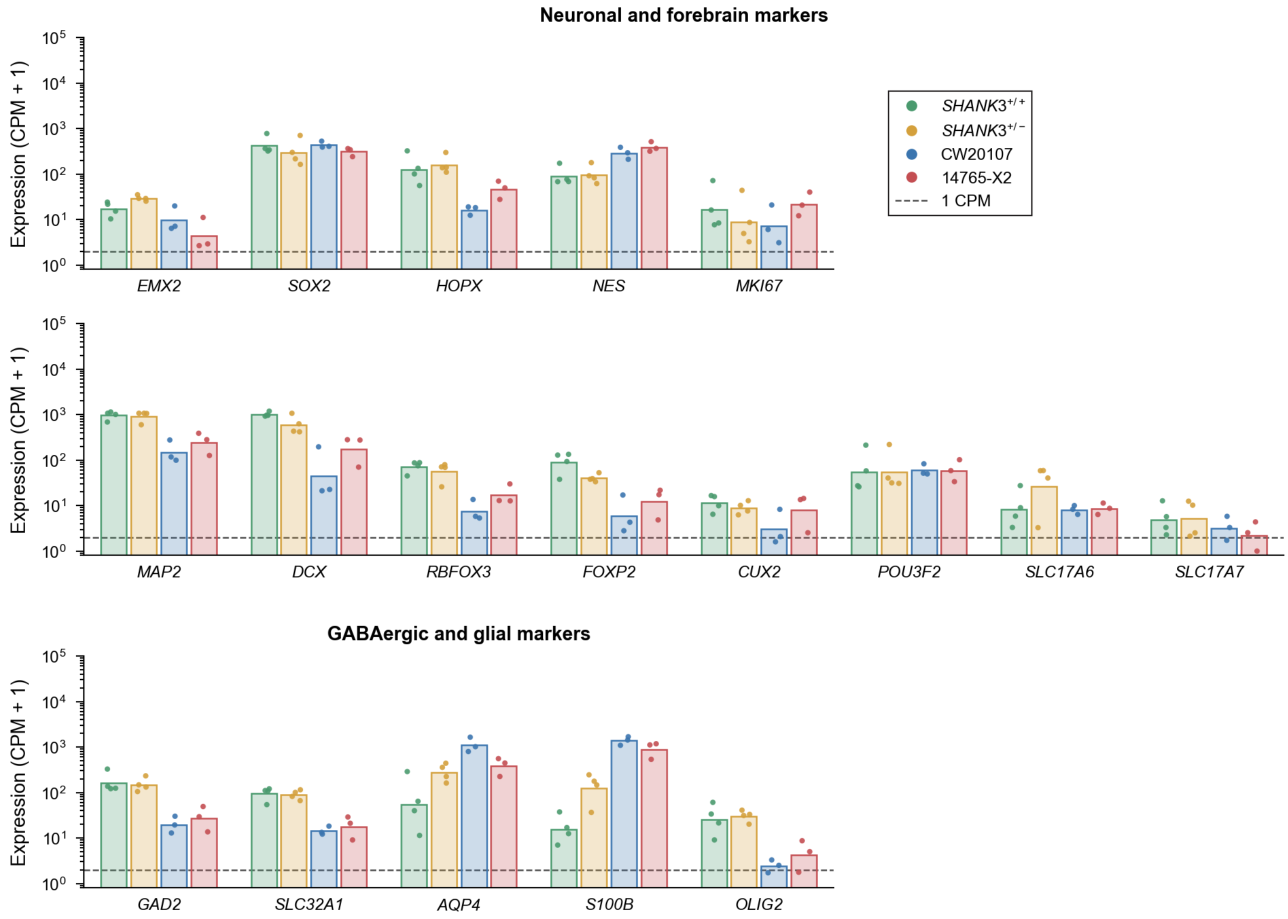


Figure S4: **Both human preparations express neuronal, forebrain, GABAergic and glial marker genes, related to Figure 3.**
Bulk RNA-seq expression of 18 marker genes not shown in Figure 3B, grouped as neuronal and forebrain, and GABAergic and glial, in *SHANK3*$^{+/+}$ ($n = 4$), *SHANK3*$^{+/-}$ ($n = 4$), CW20107 ($n = 3$) and 14765-X2 ($n = 3$) organoid samples (each pooled from three organoids; day 180 for CW20107 and 14765-X2, day 100 for KOLF2.2J), plotted as counts per million (CPM) +1 on a log axis. Dots, samples; bars, geometric mean; dashed line, 1 CPM.

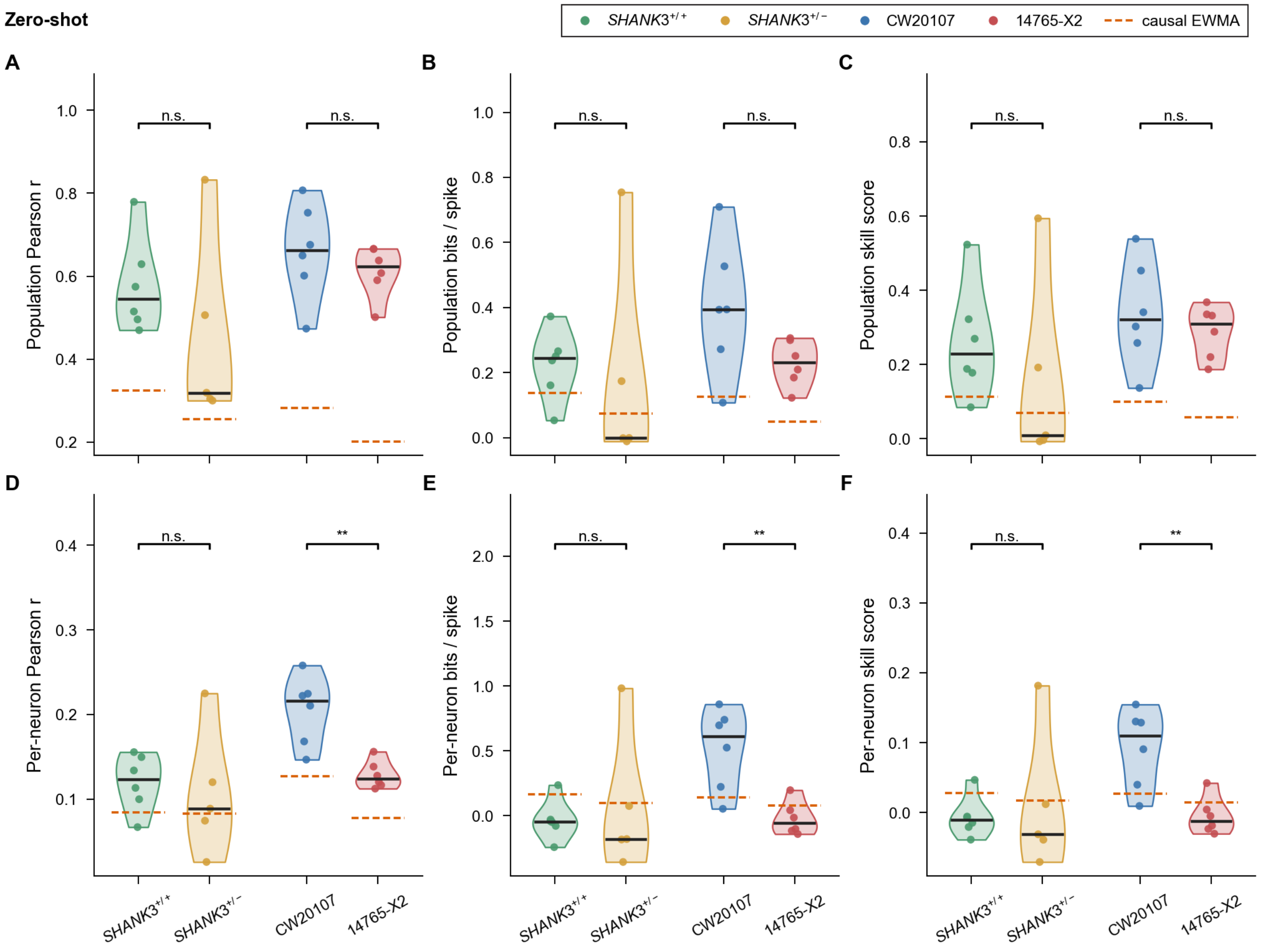


Figure S5: **Zero-shot prediction accuracy by human line, related to Figure 3.** (A–F) All six prediction-accuracy metrics of Figure 2 for zero-shot prediction across both ASD comparisons: *SHANK3*$^{+/+}$ versus *SHANK3*$^{+/-}$ ($n = 6$ versus 5 organoids) and CW20107 versus 14765-X2 ($n = 6$ versus 6 organoids). Points are organoids (the mean of their experiments), bars are medians of organoids, and the dashed line is that line's causal EWMA reference level (median across its organoids). Brackets are two-sided exact permutation Mann–Whitney $U$ tests across organoids (each organoid summarized by the mean of its experiments); $^{*}P < 0.05$, $^{**}P < 0.01$, $^{***}P < 0.001$. The two comparisons come from independent preparations and are never contrasted with each other; effect sizes and $P$ values for every cell are given in Table S12.

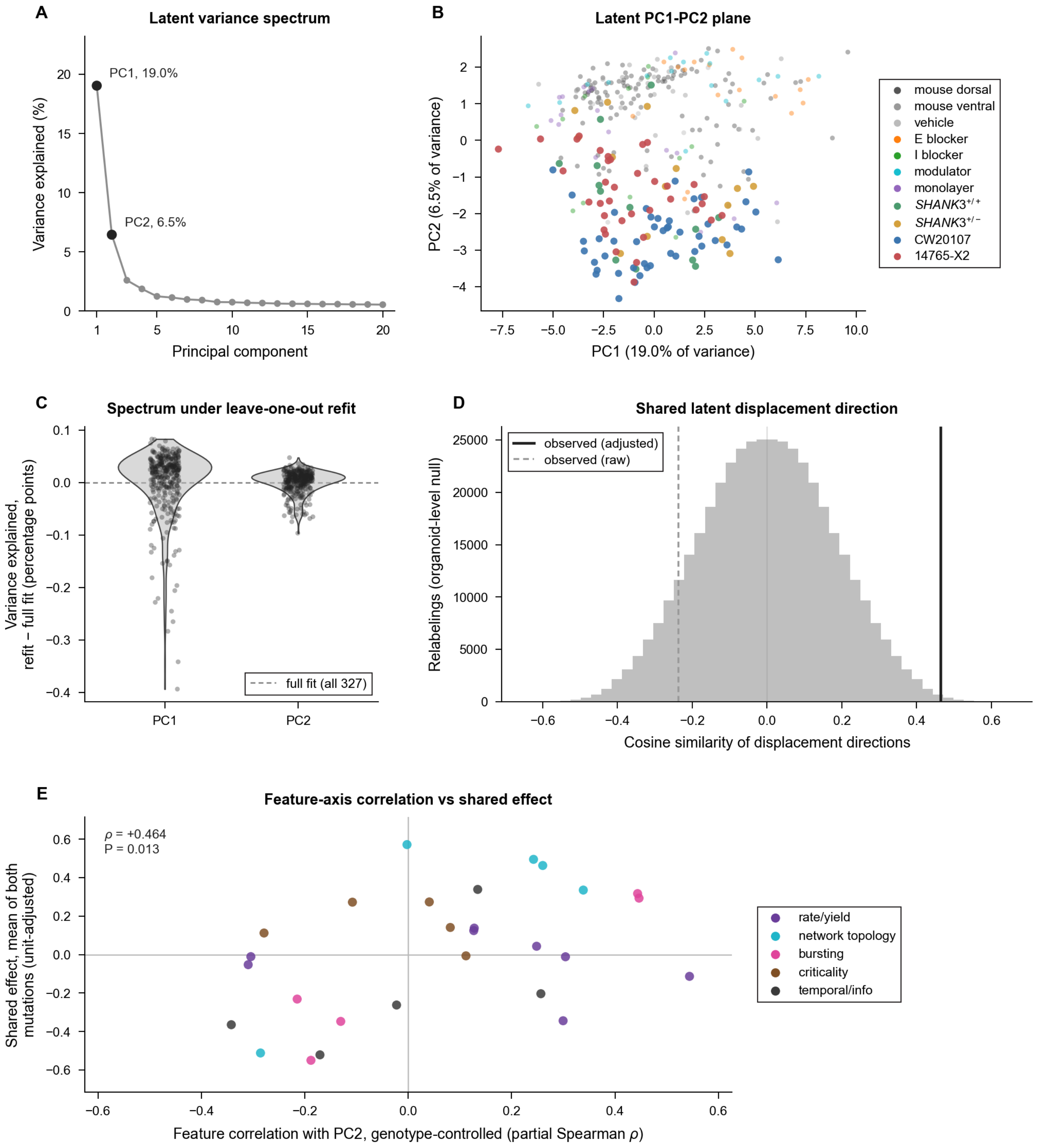
A
Latent variance spectrum
PC1, 19.0%
PC2, 6.5%
Variance explained (%)
Principal component
B
Latent PC1-PC2 plane
PC2 (6.5% of variance)
PC1 (19.0% of variance)
mouse dorsal
mouse ventral
vehicle
E blocker
I blocker
modulator
monolayer
SHANK3+/+
SHANK3+/−
CW20107
14765-X2
C
Spectrum under leave-one-out refit
Variance explained,
refit − full fit (percentage points)
full fit (all 327)
PC1
PC2
D
Shared latent displacement direction
observed (adjusted)
observed (raw)
Relabelings (organoid-level null)
Cosine similarity of displacement directions
E
Feature-axis correlation vs shared effect
ρ = +0.464
P = 0.013
Shared effect, mean of both
mutations (unit-adjusted)
Feature correlation with PC2, genotype-controlled (partial Spearman ρ)
rate/yield
network topology
bursting
criticality
temporal/info

Figure S6: **The latent axis is stable under resampling and aligned with the shared mutation effect, related to Figure 4.**

(A) Scree plot of variance explained by the first 20 principal components of the latent covariance, fitted across all 327 spontaneous recordings.

(B) All 327 recordings in the PC1–PC2 plane, colored by cohort and, for human recordings, by line.

(C) Stability of each component's variance share across 327 leave-one-out refits, plotted as the difference from the full fit. Every point is one refit.

(D) Cosine similarity between the latent displacement directions of the two NPD lines, each against its own control with log unit count regressed out within preparation, against the exact null from all 426,888 relabelings of genotype across organoids within each preparation ($924 \times 462$; histogram). Solid line, observed; dashed line, observed without unit adjustment. The organoid bootstrap 95% confidence interval of the observed cosine is $[-0.006, +0.500]$.

(E) Each spiking feature's partial Spearman correlation with PC2 across human recordings (log unit count, preparation and genotype controlled) against its mean unit-adjusted effect across the two NPD comparisons. Each point is one feature ($n = 28$), colored by family; Spearman $\rho$ and $P$ shown.

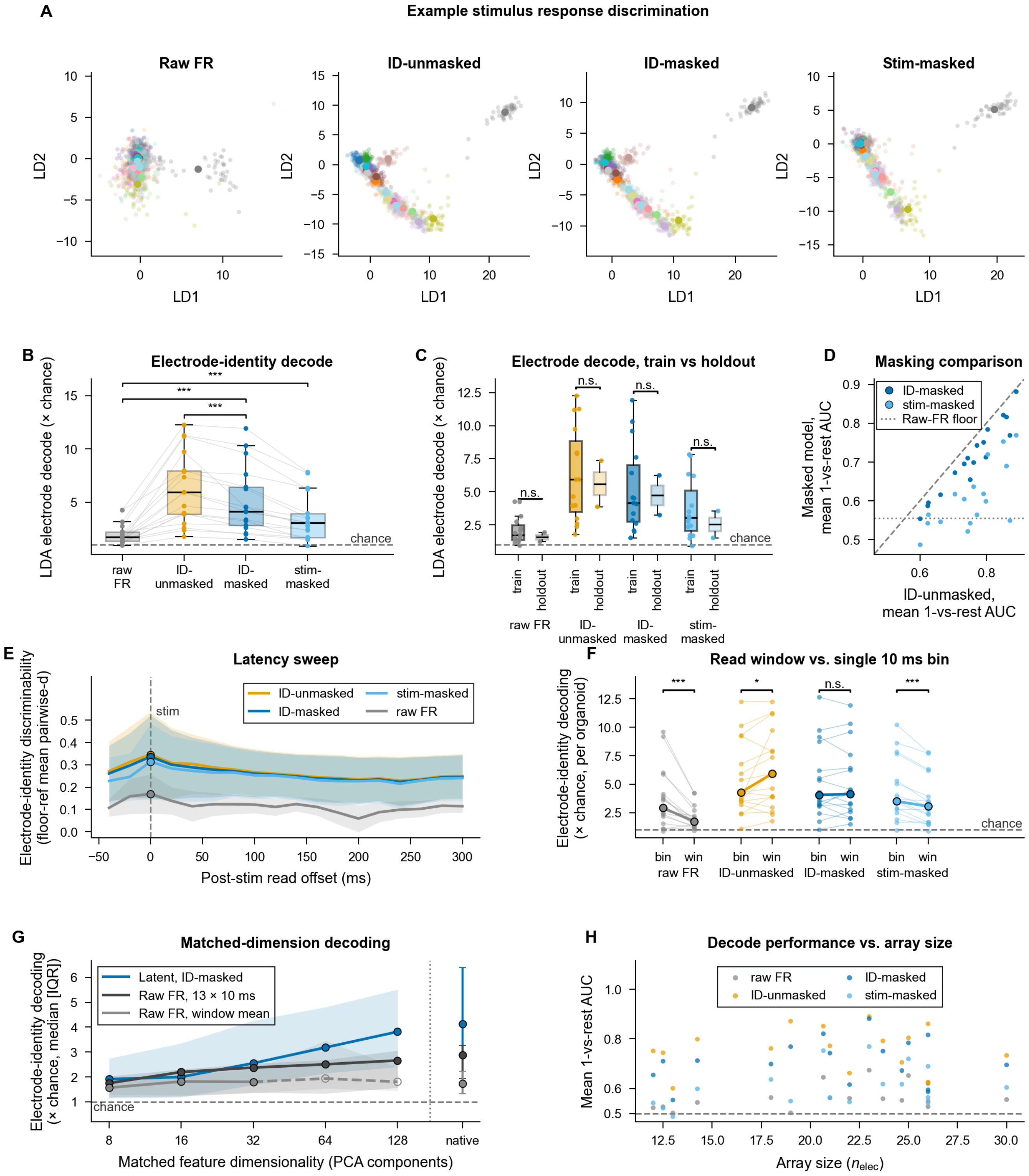

A
Example stimulus response discrimination
Raw FR
ID-unmasked
ID-masked
Stim-masked
LD1
LD2
B
Electrode-identity decode
LDA electrode decode (× chance)
raw FR
ID-unmasked
ID-masked
stim-masked
chance
***
C
Electrode decode, train vs holdout
train
holdout
n.s.
D
Masking comparison
ID-masked
stim-masked
Raw-FR floor
Masked model, mean 1-vs-rest AUC
ID-unmasked, mean 1-vs-rest AUC
E
Latency sweep
Electrode-identity discriminability (floor-ref mean pairwise-d)
raw FR
stim
Post-stim read offset (ms)
F
Read window vs. single 10 ms bin
Electrode-identity decoding (× chance, per organoid)
bin
win
*
G
Matched-dimension decoding
Latent, ID-masked
Raw FR, 13 × 10 ms
Raw FR, window mean
Electrode-identity decoding (× chance, median [IQR])
native
Matched feature dimensionality (PCA components)
H
Decode performance vs. array size
Mean 1-vs-rest AUC
Array size ($n_{elec}$)

Figure S7: **The stimulated electrode is decodable from the frozen latent, robust to readout timing, masking condition, featurization, feature dimensionality, and array size, related to Figure 5.**
(A) Two-dimensional LDA projections of single-trial evoked responses in one example recording (22 stimulation electrodes; colors, electrodes; large points, electrode means), for the raw firing rate and for the latent with the stimulation input intact (ID-unmasked), with electrode identity hidden (ID-masked), or with all stimulation input hidden (stim-masked).
(B) Electrode decoding (LDA balanced accuracy relative to chance) for the four readouts; lines join the same organoid. Brackets, paired two-sided Wilcoxon signed-rank tests, Holm-corrected across the three drawn contrasts ($n = 17$ organoids; tests use per-organoid means).
(C) Decoding in recordings from the pretraining corpus (train, $n = 15$ organoids) and from holdout organoids never used in pretraining (holdout, $n = 2$ organoids); two-sided Mann-Whitney $U$ test on per-organoid means within each readout (all n.s.; with two holdout organoids this comparison is descriptive).
(D) Mean one-vs-rest AUC of the ID-masked and stim-masked latents against the ID-unmasked latent, one point per organoid ($n = 17$ organoids); dashed line, identity; dotted line, median of the raw firing rate.
(E) Post-stimulus readout latency sweep across masking variants (mean $\pm$ s.e.m., $n = 3$ organoids).
(F) Window (120 ms) versus single-bin (10 ms) decoding, scored within arm at matched dimensionality. Lines join paired values; brackets, two-sided Wilcoxon signed-rank test within each arm ($n = 17$ organoids; tests use per-organoid means), each arm a separate comparison.
(G) Decoding at matched PCA dimensionality ($k = 8$ to 128; median [IQR], $n = 17$ organoids); the raw firing-rate, window-mean arm is a subset of the cohort at $k \geq 32$ ($n = 17, 14, 9$ organoids) and is drawn open and dashed.
(H) Decode performance versus number of electrodes in the recording ($n = 17$ organoids).
Boxes, median and IQR; whiskers, 1.5 times the IQR. In (E), line, mean; band, s.e.m. In (G), line, median; band, IQR. $^{*}P < 0.05$; $^{**}P < 0.01$; $^{***}P < 0.001$; n.s., $P \geq 0.05$. One recording was excluded because its pulses could not be aligned to the stimulation artifact (STAR Methods).

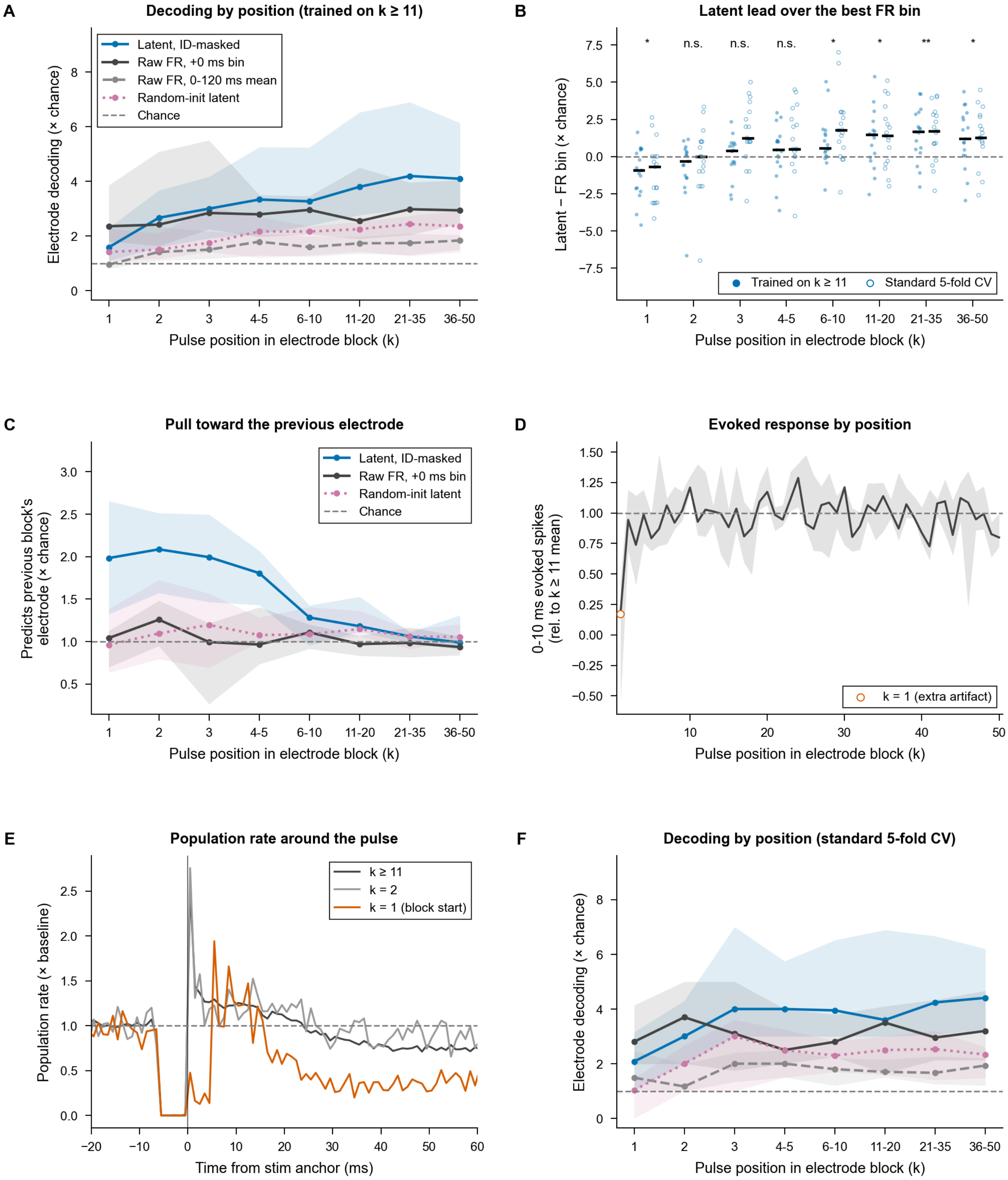

A
Decoding by position (trained on k ≥ 11)
Latent, ID-masked
Raw FR, +0 ms bin
Raw FR, 0-120 ms mean
Random-init latent
Chance
Electrode decoding (× chance)
Pulse position in electrode block (k)
B
Latent lead over the best FR bin
* n.s. n.s. n.s. * * ** *
Latent − FR bin (× chance)
Trained on k ≥ 11
Standard 5-fold CV
Pulse position in electrode block (k)
C
Pull toward the previous electrode
Predicts previous block's electrode (× chance)
Pulse position in electrode block (k)
D
Evoked response by position
0-10 ms evoked spikes (rel. to k ≥ 11 mean)
k = 1 (extra artifact)
Pulse position in electrode block (k)
E
Population rate around the pulse
Population rate (× baseline)
k ≥ 11
k = 2
k = 1 (block start)
Time from stim anchor (ms)
F
Decoding by position (standard 5-fold CV)
Electrode decoding (× chance)
Pulse position in electrode block (k)

Figure S8: **The latent's margin over firing rate builds over successive pulses to the same electrode, related to Figure 5.**
(A) Electrode decoding relative to chance by pulse position $k$ within each electrode's 50-pulse block, for the ID-masked latent, the firing rate in the +0 ms bin, the 0–120 ms mean firing rate and a randomly initialized latent. The decoder was trained on pulses $k \geq 11$ and scored on held-out late pulses and on every earlier pulse. Line, median; band, IQR ($n = 17$ organoids). The latent falls below the +0 ms firing-rate bin at $k = 1$ and exceeds it from about the sixth pulse onward.
(B) Latent minus the +0 ms firing-rate bin by pulse position, one point per organoid, for the decoder trained on $k \geq 11$ (filled) and for the five-fold cross-validation of Figure S7 (open); bars, median. Stars, paired two-sided Wilcoxon signed-rank test on per-organoid means at each position for the decoder trained on $k \geq 11$ ($^{*}P < 0.05$, $^{**}P < 0.01$).
(C) Predictions naming the previous block's electrode, relative to chance, by pulse position. Line, median; band, IQR.
(D) Population spike count 0–10 ms after each pulse, relative to each recording's $k \geq 11$ mean ($n = 9$ organoids with a reliable late response); open circle, $k = 1$, which carries an extra stimulation artifact. Line, median; band, IQR.
(E) Median population rate around the pulse at $k = 1$, $k = 2$ and $k \geq 11$, relative to the pre-pulse baseline ($n = 12$ organoids whose pulse anchor sits on the stimulation artifact).
(F) As in (A), scored by the five-fold cross-validation of Figure S7.
$^{*}P < 0.05$; $^{**}P < 0.01$; n.s., $P \geq 0.05$.

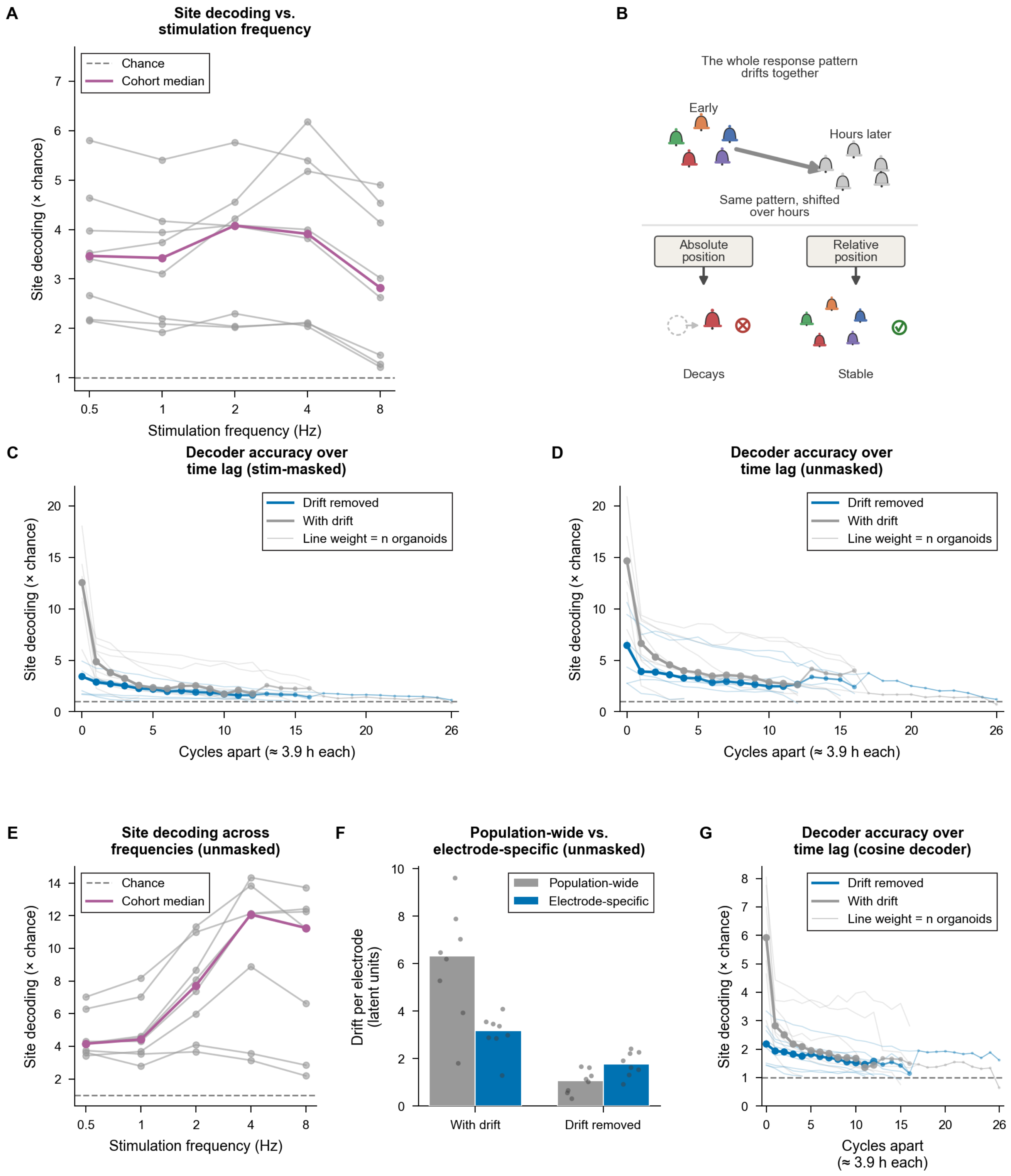

A
Site decoding vs. stimulation frequency
Chance
Cohort median
Site decoding (× chance)
Stimulation frequency (Hz)
B
The whole response pattern drifts together
Early
Hours later
Same pattern, shifted over hours
Absolute position
Relative position
Decays
Stable
C
Decoder accuracy over time lag (stim-masked)
Drift removed
With drift
Line weight = n organoids
Site decoding (× chance)
Cycles apart (≈ 3.9 h each)
D
Decoder accuracy over time lag (unmasked)
Drift removed
With drift
Line weight = n organoids
Site decoding (× chance)
Cycles apart (≈ 3.9 h each)
E
Site decoding across frequencies (unmasked)
Chance
Cohort median
Site decoding (× chance)
Stimulation frequency (Hz)
F
Population-wide vs. electrode-specific (unmasked)
Population-wide
Electrode-specific
Drift per electrode (latent units)
With drift
Drift removed
G
Decoder accuracy over time lag (cosine decoder)
Drift removed
With drift
Line weight = n organoids
Site decoding (× chance)
Cycles apart (≈ 3.9 h each)

Figure S9: **Site identity is decodable at every stimulation frequency; unmasked latents inflate site decoding, and a covariance-free decoder reproduces the readout contrast, related to Figure 5.**
(A) Site decoding relative to chance across stimulation frequency (stim-masked arm; $n = 8$ organoids). Gray lines, organoids; colored line, cohort median.
(B) Schematic: site signatures are carried off together by a common shift; their relative arrangement is preserved, so a readout referenced to each pulse's own baseline survives the drift while an absolute-position readout does not.
(C) Figure 5F in units of chance: site decoding of a decoder fit on one recording and tested on another, as a function of recording separation, for the baseline-subtracted and uncorrected readouts (stim-masked arm, shrinkage LDA; $n = 8$ organoids).
(D) As in (C), with stimulation channels left unmasked ($n = 8$ organoids); (C) and (D) share a y-axis.
(E) As in (A), with stimulation channels left unmasked ($n = 8$ organoids).
(F) Figure 5E recomputed with stimulation channels left unmasked ($n = 8$ organoids). Bars, median; dots, organoids.
(G) Figure 5F recomputed with a cosine nearest-template decoder, which fits no covariance and so cannot be affected by a stale whitening from the training recording, on the same stim-masked arm ($n = 8$ organoids). The baseline-subtracted readout again retained more of its within-cycle accuracy than the uncorrected readout at a separation of 7 to 12 cycles (median 70% vs. 28%; 7 of 7 organoids, $P = 0.016$).

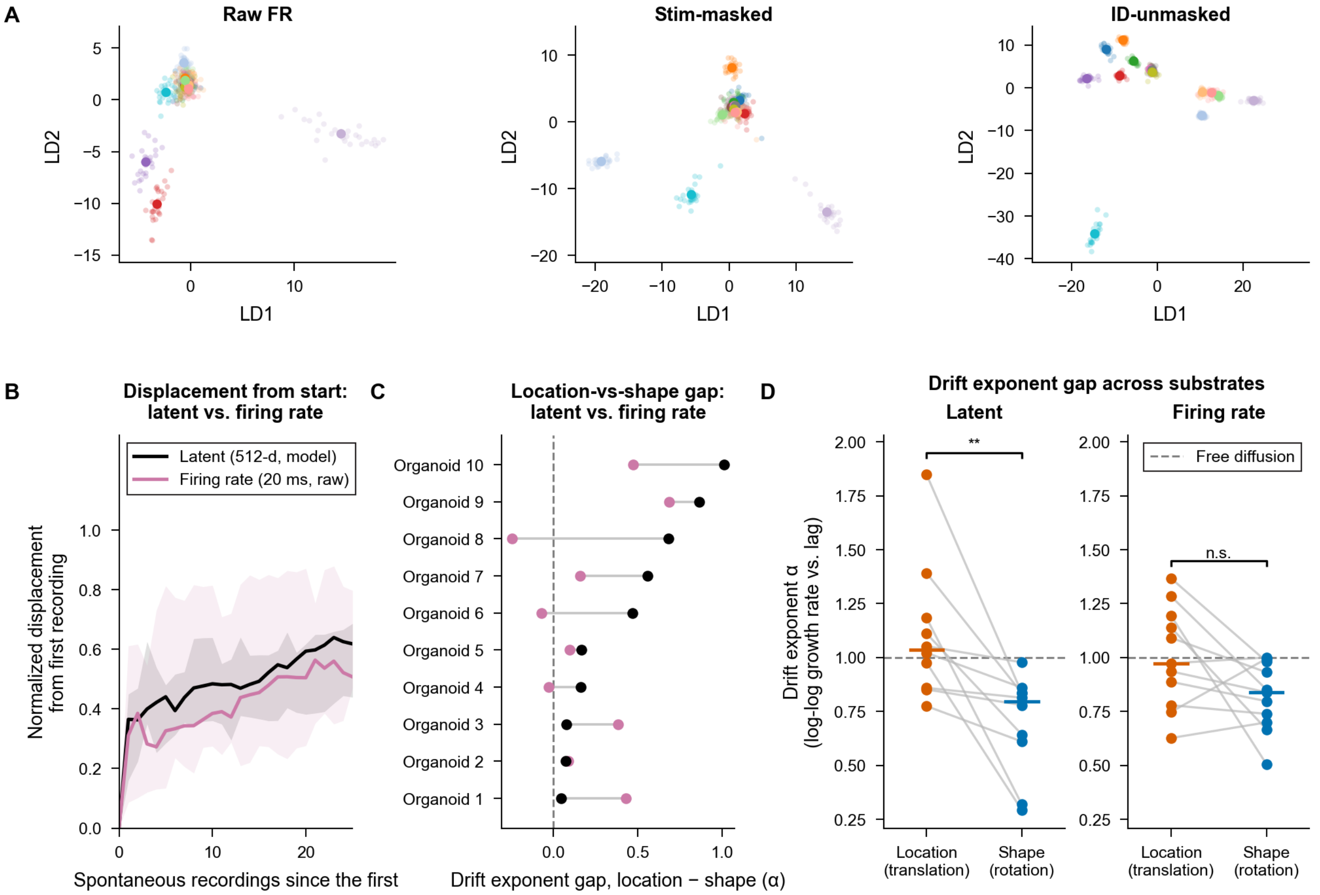


Figure S10: **Latent and firing-rate substrates recover comparable pseudotime drift and site-decoding structure, related to Figure 5.**

(A) Two-dimensional LDA electrode-cluster geometry at 8 Hz stimulation frequency for three input arms (raw FR, stim-masked latent, and ID-unmasked latent).

(B) Pseudotime displacement for the latent walk versus the raw firing-rate trajectory ($n = 10$ organoids). Shaded band, IQR across organoids; bold line, cohort median.

(C) Channel gap forest plot comparing chronological recording-order recovery (location and shape terms) between latent and binned firing rates ($n = 10$ organoids).

(D) Drift exponent (log-log growth rate versus lag) for the location and shape terms, compared between latents ($n = 10$ organoids) and binned firing rates ($n = 11$ organoids). Paired two-sided Wilcoxon signed-rank test on per-organoid means; the location versus shape contrast is significant in the latent but only a trend in firing rates.

$^{*}P < 0.05$; $^{**}P < 0.01$; $^{***}P < 0.001$.

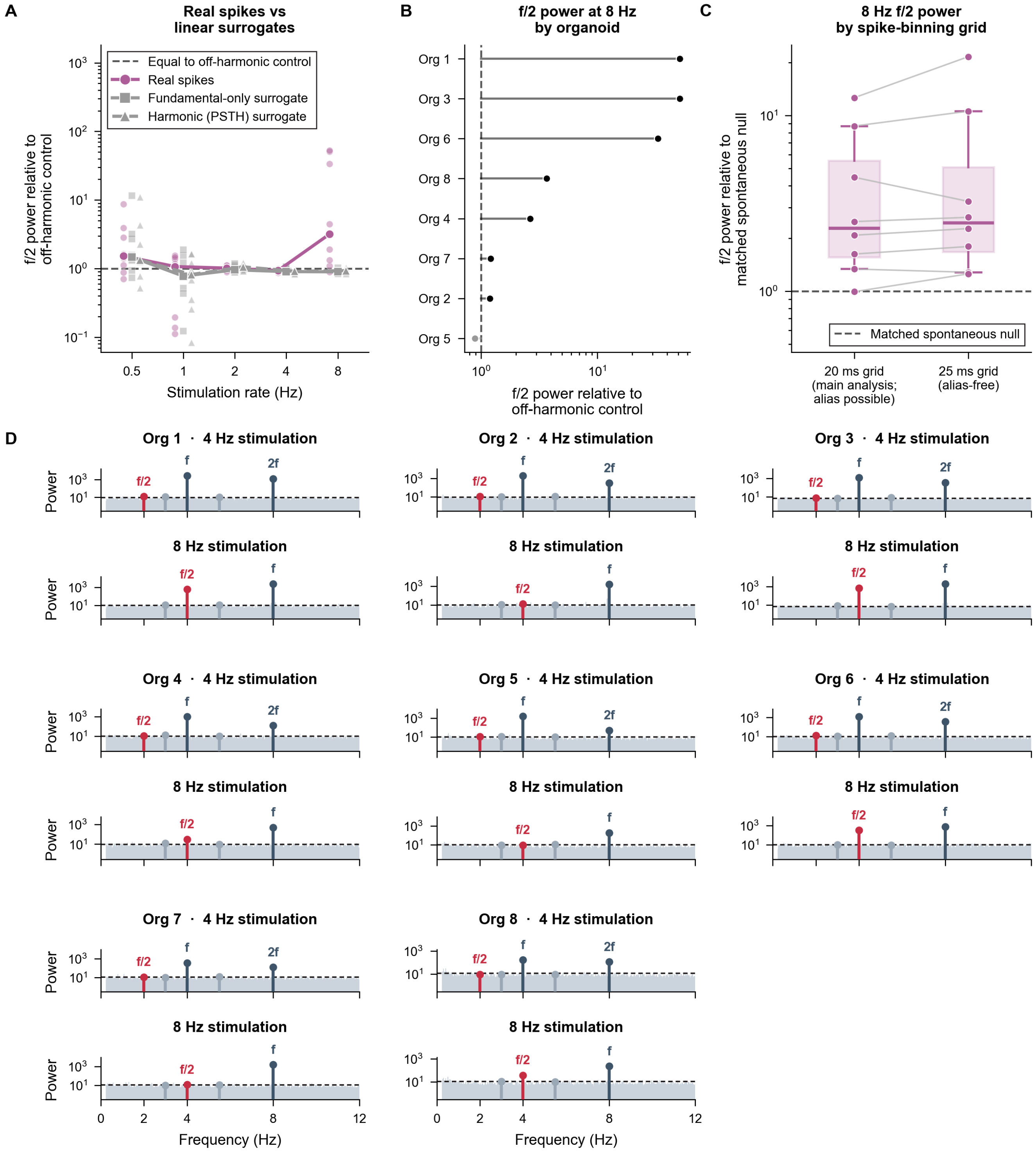

A
Real spikes vs linear surrogates
Equal to off-harmonic control
Real spikes
Fundamental-only surrogate
Harmonic (PSTH) surrogate
f/2 power relative to off-harmonic control
Stimulation rate (Hz)
B
f/2 power at 8 Hz by organoid
Org 1
Org 3
Org 6
Org 8
Org 4
Org 7
Org 2
Org 5
f/2 power relative to off-harmonic control
C
8 Hz f/2 power by spike-binning grid
f/2 power relative to matched spontaneous null
Matched spontaneous null
20 ms grid (main analysis; alias possible)
25 ms grid (alias-free)
D
Org 1 · 4 Hz stimulation
Org 2 · 4 Hz stimulation
Org 3 · 4 Hz stimulation
8 Hz stimulation
Org 4 · 4 Hz stimulation
Org 5 · 4 Hz stimulation
Org 6 · 4 Hz stimulation
Org 7 · 4 Hz stimulation
Org 8 · 4 Hz stimulation
Power
f/2
f
2f
Frequency (Hz)

Figure S11: **The $f/2$ subharmonic is generated by the network and not by the model, related to Figure 6.**
(A) $f/2$ subharmonic power measured on three inputs passed through the same model: recorded spikes, a surrogate carrying only the stimulus fundamental, and a surrogate carrying the fundamental plus harmonics. Both surrogates are period-$1/f$ by construction and contain no $f/2$ ($n = 8$ organoids); power is relative to the off-harmonic control frequencies, as in (B).
(B) Per-organoid $f/2$ power relative to the larger of the two off-harmonic control frequencies (3.0 and 5.5 Hz) in the same recording at 8 Hz stimulation ($n = 8$ organoids). Each point is one organoid; stems extend from equal power (dashed line at 1.0).
(C) $f/2$ subharmonic power in the recorded firing rates on an alias-free 25 ms bin grid, on which 8 Hz falls at exactly 5 bins per cycle so no bin-alternation alias is possible, relative to $f/2$ power in spontaneous recordings and to 4 Hz ($n = 8$ organoids; Table S17).
(D) Power spectrum of each organoid (the mean of its experiments' example blocks) at 4 and 8 Hz stimulation, with the stimulation frequency ($f$), its harmonic ($2f$) and the subharmonic ($f/2$) marked; dashed line, empirical noise ceiling (99.9th percentile of power away from the marked lines; $n = 8$ organoids).

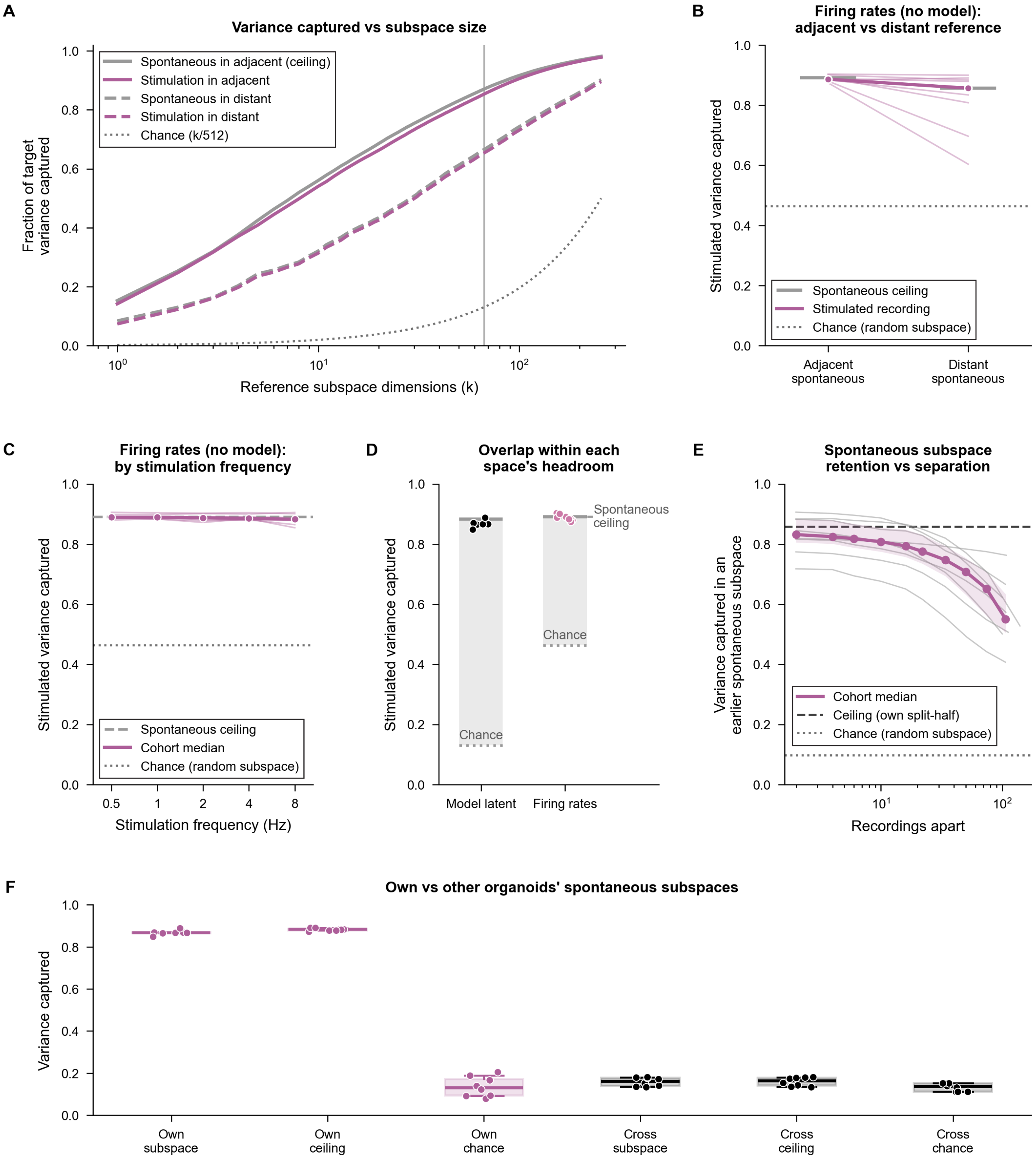
A
Variance captured vs subspace size
Fraction of target variance captured
Reference subspace dimensions (k)
Spontaneous in adjacent (ceiling)
Stimulation in adjacent
Spontaneous in distant
Stimulation in distant
Chance (k/512)
B
Firing rates (no model): adjacent vs distant reference
Stimulated variance captured
Spontaneous ceiling
Stimulated recording
Chance (random subspace)
Adjacent spontaneous
Distant spontaneous
C
Firing rates (no model): by stimulation frequency
Stimulation frequency (Hz)
Spontaneous ceiling
Cohort median
Chance (random subspace)
D
Overlap within each space's headroom
Spontaneous ceiling
Chance
Model latent
Firing rates
E
Spontaneous subspace retention vs separation
Variance captured in an earlier spontaneous subspace
Recordings apart
Cohort median
Ceiling (own split-half)
Chance (random subspace)
F
Own vs other organoids' spontaneous subspaces
Variance captured
Own subspace
Own ceiling
Own chance
Cross subspace
Cross ceiling
Cross chance

Figure S12: **Subspace overlap depends on neither subspace size nor stimulation rate and reproduces without the model, the spontaneous geometry reorganizes continuously, and subspace overlap is organoid-specific, related to Figure 6.**
(A) Fraction of variance captured as spontaneous directions are added one at a time ($k = 1$ to 256 of 512 latent dimensions), for stimulation and spontaneous activity projected onto the adjacent and distant spontaneous recordings, against the $k/512$ diagonal expected of a randomly oriented subspace.
(B) The same overlap measured on binned firing rates with no model in the path, for the adjacent versus the experiment's first (distant) spontaneous recording ($n = 8$ organoids).
(C) The firing-rate overlap split by stimulation rate, the rate-space counterpart of Figure 6G ($n = 8$ organoids).
(D) Stimulation-in-adjacent-spontaneous overlap normalized to each space's own ceiling and chance floor, the one comparison that is fair across spaces of different ambient dimension ($n = 8$ organoids; two-sided Wilcoxon signed-rank test).
(E) Variance of one spontaneous recording retained in a fixed 50-dimensional subspace of another spontaneous recording, as a function of recording separation ($n = 8$ organoids).
(F) Fraction of stimulation-evoked variance falling inside the same organoid's own preceding spontaneous subspace versus inside other organoids' spontaneous subspaces, against the chance floor for a randomly oriented subspace of the same size ($n = 8$ organoids; Table S17).
In (E) and (F), shaded band is IQR; bold line, cohort median. Dashed lines, spontaneous-on-spontaneous ceiling; dotted lines, chance (random subspace). Panels (B) and (C) are firing-rate only: the latent and rate spaces have different ambient dimensions, so their chance floors differ several-fold and the raw captured fractions are not comparable between them.

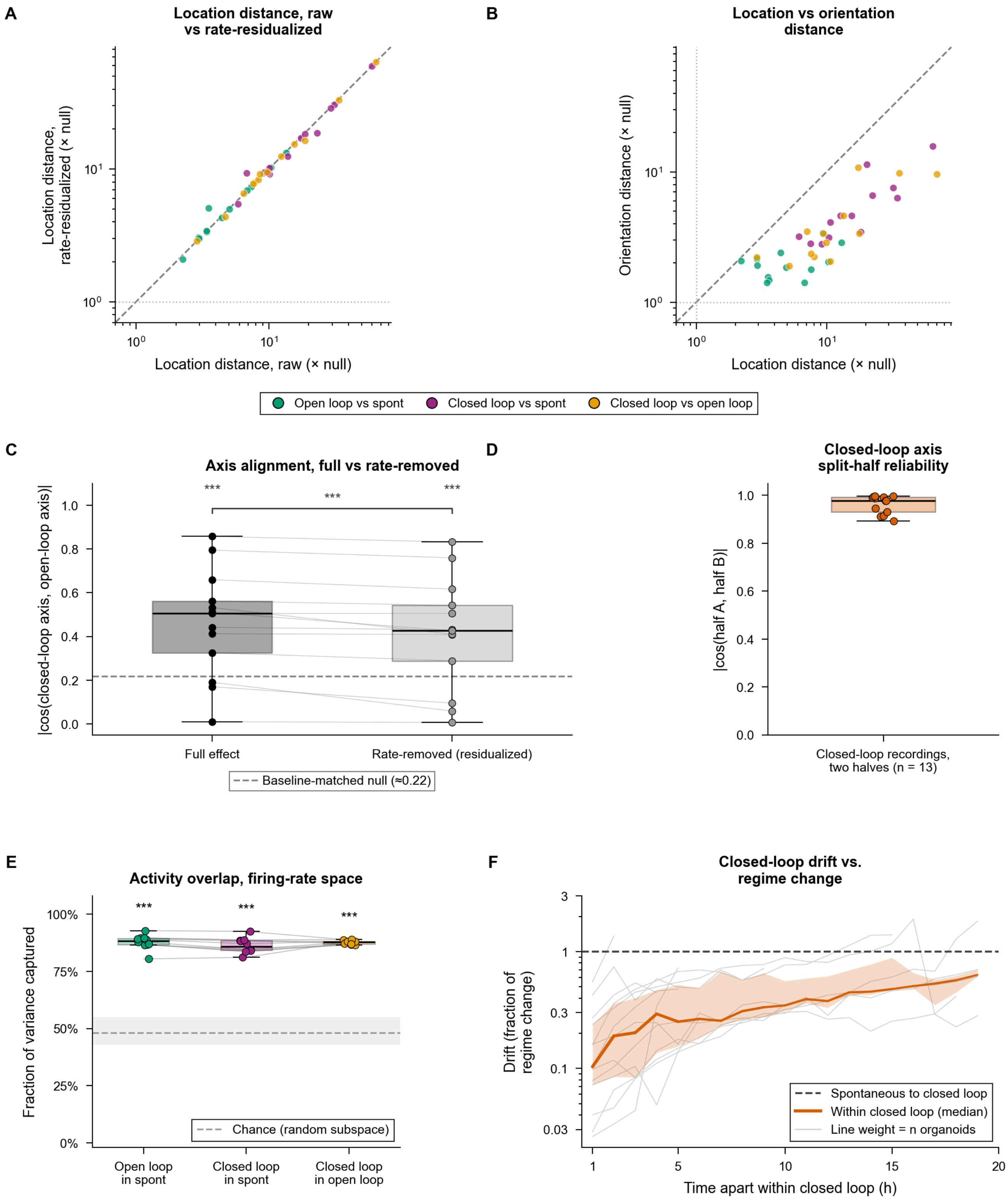
A
Location distance, raw vs rate-residualized
Location distance, rate-residualized (× null)
Location distance, raw (× null)
B
Location vs orientation distance
Orientation distance (× null)
Location distance (× null)
Open loop vs spont
Closed loop vs spont
Closed loop vs open loop
C
Axis alignment, full vs rate-removed
|cos(closed-loop axis, open-loop axis)|
Full effect
Rate-removed (residualized)
Baseline-matched null (≈0.22)
D
Closed-loop axis split-half reliability
|cos(half A, half B)|
Closed-loop recordings, two halves (n = 13)
E
Activity overlap, firing-rate space
Fraction of variance captured
Chance (random subspace)
Open loop in spont
Closed loop in spont
Closed loop in open loop
F
Closed-loop drift vs. regime change
Drift (fraction of regime change)
Time apart within closed loop (h)
Spontaneous to closed loop
Within closed loop (median)
Line weight = n organoids

Figure S13: **Closed-loop state separation survives firing-rate residualization, and the closed-loop axis is reliable and reused in firing-rate space, related to Figure 7.**
(A) Centroid location shift between condition pairs (spontaneous, open-loop, closed-loop) recomputed after residualizing out population firing rate, plotted against the raw shift ($n = 13$ organoids; each point is one organoid, the log-scale mean of its experiments).
(B) Location displacement term plotted against the orientation (shape) displacement term, each relative to the within-condition drift null ($n = 13$ organoids). The ordering of the orientation term among pairs reported in the text uses paired Wilcoxon signed-rank tests on $\log_2$ ratios, one-sided for the two contrasts testing that the spontaneous–closed-loop term exceeds each of the other two and two-sided for the third, Holm-corrected across the three contrasts.
(C) Cosine alignment between the closed-loop axis and the open-loop axis, computed on the full axes and after removing each organoid's own firing-rate axis, benchmarked against a baseline-matched permutation null ($n = 13$ organoids; dots, organoids, each the mean of its experiments; lines join an organoid across the two axes).
(D) Split-half reliability of the closed-loop axis, estimated independently from two disjoint halves of each experiment's closed-loop recordings (dots, organoids, each the mean of its experiments; $n = 13$).
(E) Fraction of variance captured by each pairwise overlap, computed directly on firing rates ($n = 10$ organoids; dots, organoids; lines join organoids with enough balance and failure episodes, the cohort of Figure S14E).
(F) Latent drift between two closed-loop recordings of the same organoid against the time between them (one 15-min recording per hour), as a fraction of that organoid's own spontaneous-to-closed-loop location change (dashed line). Thin lines, organoids (each the mean of its experiments); bold line, median across organoids; band, IQR; line weight, number of organoids reaching each gap (13 at 1 to 2 h, falling to 5 at 19 h; gaps reached by fewer than 5 organoids are omitted). Paired two-sided Wilcoxon signed-rank test against each organoid's own change at each gap, Holm-corrected; statistics in the text.
In (A) and (B), each point is one organoid. In (C) to (E), box plots: center line, median; box, IQR; whiskers, 1.5× IQR. In (C), stars from a permutation test that exchanges whole organoids and bracket from a paired Wilcoxon signed-rank test; in (E), one-sided Wilcoxon signed-rank test against chance of the organoids' margins over their own chance floor; $^{*}P < 0.05$, $^{**}P < 0.01$, $^{***}P < 0.001$.

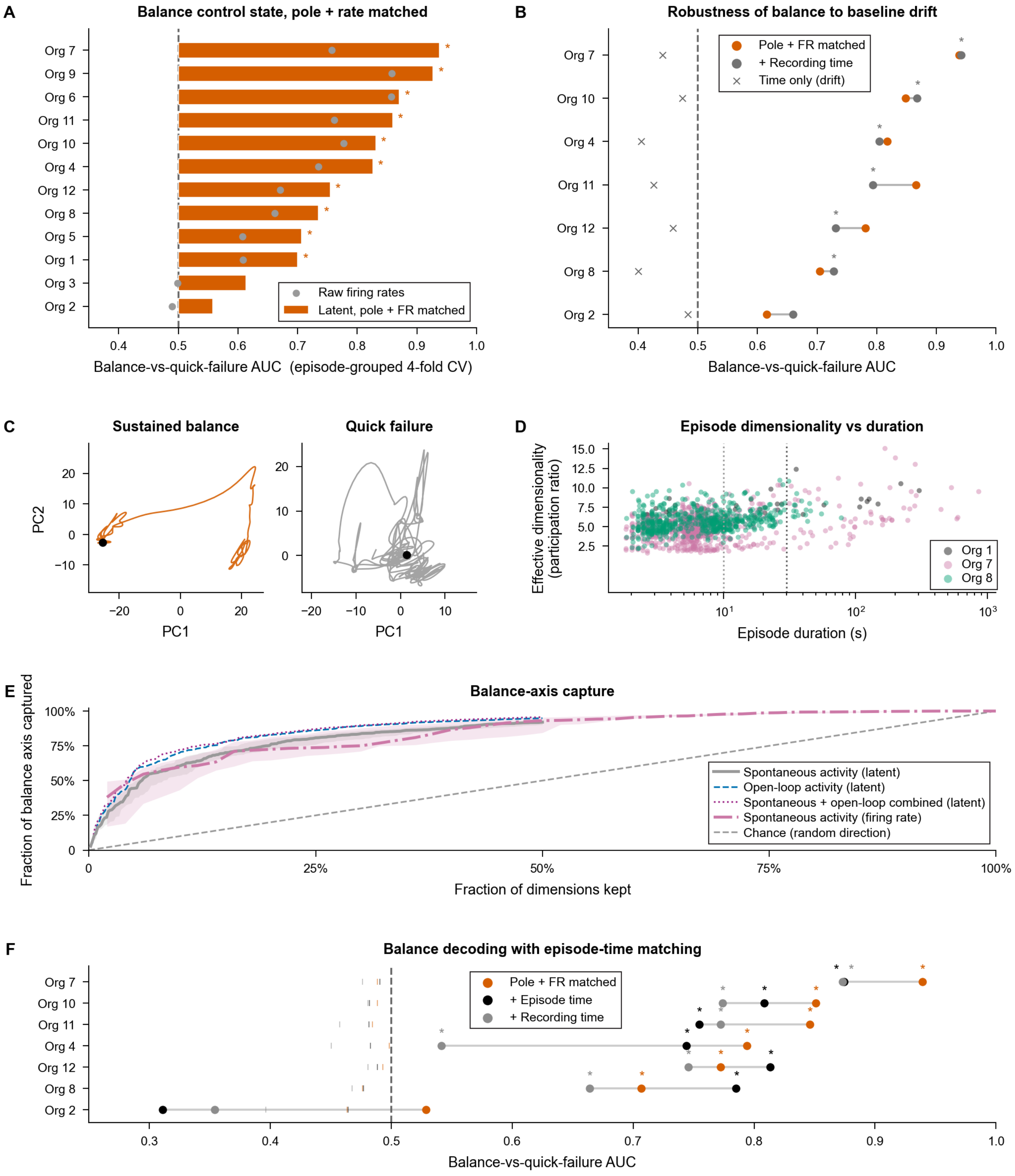

A
Balance control state, pole + rate matched
Raw firing rates
Latent, pole + FR matched
Balance-vs-quick-failure AUC (episode-grouped 4-fold CV)
B
Robustness of balance to baseline drift
Pole + FR matched
+ Recording time
Time only (drift)
Balance-vs-quick-failure AUC
C
Sustained balance
Quick failure
PC1
PC2
D
Episode dimensionality vs duration
Effective dimensionality (participation ratio)
Episode duration (s)
Org 1
Org 7
Org 8
E
Balance-axis capture
Fraction of balance axis captured
Fraction of dimensions kept
Spontaneous activity (latent)
Open-loop activity (latent)
Spontaneous + open-loop combined (latent)
Spontaneous activity (firing rate)
Chance (random direction)
F
Balance decoding with episode-time matching
Pole + FR matched
+ Episode time
+ Recording time
Balance-vs-quick-failure AUC

Figure S14: **Latent balance decoder exceeds firing-rate baseline, and balance episodes trace lower-dimensional orbits, related to Figure 7.**
(A) Matched linear-latent AUC (mean absolute pole angle, its within-window variability and population firing rate matched via coarsened exact matching) versus a per-neuron firing-rate decoder on the same matched windows, one bar per organoid (the mean over its experiments; $n = 12$ organoids); statistics in the text.
(B) Robustness of the balance control-state decoder to slow within-recording drift: matched AUC without versus with time-in-recording added as an additional matching covariate, one pair of points per organoid ($n = 7$ organoids).
(C) Smoothed top-2 principal-component latent orbits for a sustained-balance episode and a quick-failure episode from the same organoid.
(D) Per-episode latent topology (participation-ratio effective dimensionality) plotted against balance-episode duration, points colored by organoid (1,387 episodes from three organoids).
(E) Balance-axis capture curve: cumulative fraction of the balance-versus-failure readout direction captured by the leading principal components of spontaneous, open-loop, or pooled spontaneous and open-loop activity in the latent, and of spontaneous activity in raw firing rates, as more dimensions are retained, against the chance diagonal of a randomly oriented direction ($n = 10$ organoids; each curve is the mean of the organoid's experiments). The x-axis is the fraction of dimensions kept (latent, of 512; firing rate, of each experiment's unit count), so both chance floors fall on one diagonal; latent curves end at 256 of 512 dimensions.
(F) Balance-versus-quick-failure decoding AUC with episode time (time since episode start) added as a matching covariate, alongside the pole- and firing-rate-matched decoder and a decoder matched on both episode time and time in the recording, one point per organoid ($n = 7$ organoids).
In (A, B and F), $^*P < 0.05$ by the Fisher-combined episode-label permutation $P$ of the organoid's experiments (each experiment's own permutation test, 1,000 draws); bars extend from chance (0.5). In (E), shaded band is IQR; bold line, median across organoids.

### Supplemental tables

Table S1: **Data corpus census**, related to Figure 1. Recordings, hours, and experiments stratified by preparation, region, and species. Counts are mutually exclusive and sum to the full corpus; experiments are distinct experiment IDs (one electrode configuration of one organoid, spike-sorted once; four organoids span two rows, so the column sums to more than its total). Org., organoids. Hours are rounded to the nearest hour (corpus total, 914.3 h). The corpus was divided into a training pool (2,027 recordings, 632.3 h) used for pretraining and a holdout set (1,173 recordings, 282.0 h) never trained on (assignment is by experiment, and about 20 of the 63 holdout organoids also contribute training experiments, almost all drug-treated). Human organoids and drug-perturbation recordings were held out by design and contributed zero gradient updates. Within the training pool, the train/validation partition varies by random seed; the experiment-level assignment to training pool versus holdout is fixed.

| Preparation | Region | Species | Hours | Recordings | Experiments | Org. |
|---|---|---|---|---|---|---|
| 3D organoid | Dorsal forebrain | Mouse | 706 | 2,511 | 256 | 66 |
| 3D organoid | Ventral forebrain | Mouse | 10 | 57 | 57 | 26 |
| 3D organoid | Dorsal forebrain | Human† | 19 | 109 | 109 | 23 |
| 2D dissociated | Dorsal forebrain | Mouse | 58 | 161 | 16 | 9 |
| 2D dissociated | Ventral forebrain | Mouse | 110 | 296 | 12 | 4 |
| *Ex vivo* slice | Primary cortex | Mouse | 11 | 66 | 53 | 4 |
| **Total** | | | **914.3** | **3,200** | **503** | **128** |

†Human organoids comprise isogenic control (*SHANK3*$^{+/+}$, $n = 13$) and neuropsychiatric disease (*SHANK3*$^{+/-}$, $n = 14$) hIPSC lines, together with two further hIPSC lines (CW20107, $n = 46$; 14765-X2, $n = 36$); all 109 recordings were held out from pretraining. These experiments come from 6, 5, 6 and 6 organoids, respectively.

Table S2: **PULSE exceeds a causal EWMA reference in distribution and on held-out experiments after two-minute fine-tuning**, related to Figure 2. PULSE and EWMA columns give the median across organoids of each organoid's value (the mean of its experiments' seed-averaged values, 3 model seeds). Training and validation rows use the Figure 2 reference model on the same recordings, seeds and neurons as Table S5, with one EWMA smoothing factor chosen on training recordings. Held-out rows use the all-recordings model, with the EWMA fitted on each recording's 2-minute adaptation window and both scored on the same evaluation window and neurons (Methods). Paired gain, median across organoids of PULSE minus EWMA (not the difference of the two medians). % > EWMA, percentage of organoids in which PULSE exceeded the EWMA. $P$, paired two-sided Wilcoxon signed-rank test across organoids, uncorrected. $n$ is the number of organoids: 86 training, 30 validation and 9 held-out cultures.

| Metric | PULSE | EWMA | Paired gain | % > EWMA | $P$ |
|---|---|---|---|---|---|
| **Training, reference model** ($n = 86$) | | | | | |
| Population Pearson $r$ | 0.749 | 0.122 | +0.576 | 100 | $8.0 \times 10^{-16}$ |
| Population bits/spike | 0.170 | 0.005 | +0.164 | 100 | $8.0 \times 10^{-16}$ |
| Population skill | 0.520 | 0.012 | +0.493 | 100 | $8.0 \times 10^{-16}$ |
| Per-neuron Pearson $r$ | 0.284 | 0.043 | +0.236 | 100 | $8.0 \times 10^{-16}$ |
| Per-neuron median bits/spike | 0.659 | −0.018 | +0.721 | 100 | $8.0 \times 10^{-16}$ |
| Per-neuron skill | 0.150 | −0.006 | +0.155 | 100 | $8.0 \times 10^{-16}$ |
| **Validation, reference model** ($n = 30$) | | | | | |
| Population Pearson $r$ | 0.723 | 0.135 | +0.522 | 100 | $1.9 \times 10^{-9}$ |
| Population bits/spike | 0.217 | 0.011 | +0.203 | 93 | $1.9 \times 10^{-8}$ |
| Population skill | 0.467 | 0.015 | +0.426 | 93 | $4.4 \times 10^{-6}$ |
| Per-neuron Pearson $r$ | 0.202 | 0.033 | +0.145 | 100 | $1.9 \times 10^{-9}$ |
| Per-neuron median bits/spike | 0.339 | −0.042 | +0.532 | 90 | $2.8 \times 10^{-6}$ |
| Per-neuron skill | 0.088 | −0.009 | +0.102 | 90 | $6.9 \times 10^{-6}$ |
| **Held-out mouse, all-recordings model, zero-shot** ($n = 9$) | | | | | |
| Population Pearson $r$ | 0.466 | 0.140 | +0.278 | 100 | 0.004 |
| Population bits/spike | 0.040 | 0.022 | +0.039 | 56 | 0.25 |
| Population skill | 0.037 | 0.032 | +0.036 | 56 | 0.73 |
| Per-neuron Pearson $r$ | 0.104 | 0.034 | +0.066 | 89 | 0.008 |
| Per-neuron median bits/spike | −0.335 | −0.028 | −0.287 | 11 | 0.05 |
| Per-neuron skill | −0.069 | −0.003 | −0.045 | 11 | 0.04 |
| **Held-out mouse, all-recordings model,** +2 **min linear head** ($n = 9$) | | | | | |
| Population Pearson $r$ | 0.597 | 0.140 | +0.468 | 100 | 0.004 |
| Population bits/spike | 0.210 | 0.022 | +0.187 | 100 | 0.004 |
| Population skill | 0.302 | 0.032 | +0.270 | 100 | 0.004 |
| Per-neuron Pearson $r$ | 0.212 | 0.034 | +0.189 | 100 | 0.004 |
| Per-neuron median bits/spike | 0.359 | −0.028 | +0.410 | 100 | 0.004 |
| Per-neuron skill | 0.074 | −0.003 | +0.089 | 89 | 0.008 |

Table S3: **Selection of the EWMA smoothing factor**, related to Figure 2. Poisson negative log-likelihood (NLL, nats per 20 ms bin per neuron, lower is better) of the causal EWMA for each smoothing factor $\alpha$, pooled over all neurons and bins of the training (86 organoids) and validation (30) data of Table S2. $\tau$, time constant of the average ($-\Delta t/\ln(1-\alpha)$, $\Delta t = 20$ ms). The factor was chosen on training recordings only (bold) and applied unchanged to validation. NLL fell monotonically toward slower smoothing on both splits; the last step ($\alpha = 0.01$ to 0.005) lowered it by less than 0.001, against a gap of 0.046 (training) and 0.035 (validation) to PULSE.

| $\alpha$ | $\tau$ (s) | Training NLL | Validation NLL |
|---|---|---|---|
| **0.005** | **3.99** | **0.2460** | **0.2418** |
| 0.01 | 1.99 | 0.2463 | 0.2421 |
| 0.02 | 0.99 | 0.2479 | 0.2437 |
| 0.05 | 0.39 | 0.2559 | 0.2514 |
| 0.1 | 0.19 | 0.2747 | 0.2695 |
| 0.2 | 0.09 | 0.3183 | 0.3115 |
| 0.4 | 0.04 | 0.4129 | 0.4026 |
| PULSE (reference model) | | 0.2003 | 0.2069 |

Table S4: **Paired comparisons of every Figures 2B–2E design-axis level against the reference configuration, validation organoids**, related to Figure 2. For each level, three rows give the median across organoids of the paired difference (Δ, level − reference, in units of $10^{-3}$, from each organoid's mean of its experiments' seed-averaged values over 3 random seeds); the percentage of organoids (%) in which the level scored better than the reference (higher, or lower for MAE and Poisson NLL); and the Holm-corrected $P$ of a paired two-sided Wilcoxon signed-rank test. Holm correction is applied per metric within each axis (bin width, recordings per experiment, model width, and the sorter-embedding and active-unit-count ablations together). †The stimulation-input ablation is scored only on experiments with evoked-stimulation recordings and is its own uncorrected test (28 organoids). $n$ is the number of organoids: 30 validation; experiments from one organoid are averaged before every statistic, and $P$ values are across organoids. Unit count, active-unit-count conditioning. Pop., population; PN, per-neuron; bits, median bits/spike for PN; MAE, mean absolute error; NLL, Poisson negative log-likelihood.

| Level | | Pop. $r$ | Pop. bits | Pop. skill | PN $r$ | PN bits | PN skill | PN MAE | PN NLL |
|---|---|---|---|---|---|---|---|---|---|
| **Bin width (vs 10 ms)** | | | | | | | | | |
| 5 ms ($n = 30$) | Δ | +6.60 | +4.91 | +10.85 | +1.29 | +0.42 | +0.13 | +0.01 | −0.16 |
| | % | 83 | 87 | 83 | 60 | 53 | 50 | 43 | 70 |
| | $P$ | $6.3\times10^{-5}$ | $6.3\times10^{-5}$ | $1.2\times10^{-5}$ | 0.02 | 0.89 | 0.70 | 0.54 | 0.04 |
| 20 ms ($n = 30$) | Δ | −16.57 | −10.88 | −20.62 | −4.74 | −23.71 | −6.48 | +0.28 | +0.97 |
| | % | 3 | 7 | 3 | 7 | 13 | 10 | 43 | 7 |
| | $P$ | $7.5\times10^{-9}$ | $9.3\times10^{-8}$ | $5.1\times10^{-7}$ | $1.9\times10^{-8}$ | $2.8\times10^{-5}$ | $2.4\times10^{-5}$ | 0.27 | $3.6\times10^{-5}$ |
| **Recordings per experiment (vs 20)** | | | | | | | | | |
| 1 ($n = 30$) | Δ | −30.37 | −34.84 | −65.73 | −46.41 | −242.85 | −60.67 | +3.34 | +12.68 |
| | % | 20 | 20 | 10 | 0 | 0 | 0 | 23 | 7 |
| | $P$ | $2.4\times10^{-4}$ | $7.1\times10^{-5}$ | $7.0\times10^{-6}$ | $5.6\times10^{-9}$ | $5.6\times10^{-9}$ | $5.6\times10^{-9}$ | $1.0\times10^{-4}$ | $5.6\times10^{-8}$ |
| 10 ($n = 30$) | Δ | −5.48 | −5.27 | −11.19 | −2.60 | −43.13 | −8.47 | −0.18 | −0.52 |
| | % | 47 | 33 | 37 | 37 | 33 | 27 | 50 | 53 |
| | $P$ | 0.28 | 0.40 | 0.20 | 0.16 | 0.18 | 0.03 | 1.00 | 0.90 |
| All ($n = 30$) | Δ | +12.12 | +6.00 | +13.24 | +3.58 | +9.55 | +3.58 | −0.04 | −2.00 |
| | % | 83 | 67 | 80 | 73 | 60 | 67 | 50 | 57 |
| | $P$ | $7.6\times10^{-4}$ | 0.40 | 0.009 | 0.003 | 0.18 | 0.03 | 1.00 | 0.90 |
| **Model width (vs $d$512)** | | | | | | | | | |
| $d$128 ($n = 30$) | Δ | −4.46 | −3.85 | −11.58 | −21.11 | −119.46 | −23.67 | +2.41 | +3.98 |
| | % | 43 | 30 | 33 | 3 | 7 | 7 | 10 | 7 |
| | $P$ | 0.33 | 0.28 | 0.18 | $2.0\times10^{-6}$ | $2.1\times10^{-5}$ | $3.5\times10^{-5}$ | $2.0\times10^{-6}$ | $1.2\times10^{-5}$ |
| $d$256 ($n = 30$) | Δ | +2.29 | +1.08 | +4.78 | −8.09 | −40.40 | −6.52 | +1.38 | +1.10 |
| | % | 67 | 57 | 60 | 13 | 7 | 10 | 3 | 13 |
| | $P$ | 0.07 | 0.28 | 0.18 | $1.4\times10^{-5}$ | $2.1\times10^{-5}$ | $1.0\times10^{-5}$ | $5.2\times10^{-8}$ | $2.8\times10^{-4}$ |

Table S4: **Paired comparisons of every Figures 2B–2E design-axis level against the reference configuration, validation organoids (continued)**, related to Figure 2.

| Level | | Pop. $r$ | Pop. bits | Pop. skill | PN $r$ | PN bits | PN skill | PN MAE | PN NLL |
|---|---|---|---|---|---|---|---|---|---|
| **Conditioning removed (vs all on)** | | | | | | | | | |
| Sorter embedding ($n = 30$) | $\Delta$ | +0.07 | +0.02 | +0.09 | +1.41 | +4.05 | +1.58 | −0.23 | −0.40 |
| | % | 50 | 50 | 53 | 87 | 73 | 73 | 70 | 87 |
| | $P$ | 1.00 | 1.00 | 1.00 | 0.001 | 0.68 | 0.29 | 0.03 | 0.002 |
| Unit count, input ($n = 30$) | $\Delta$ | −0.54 | −0.30 | −1.00 | −0.09 | −0.17 | −0.51 | +0.03 | +0.03 |
| | % | 40 | 33 | 40 | 43 | 43 | 40 | 43 | 43 |
| | $P$ | 1.00 | 1.00 | 1.00 | 0.81 | 0.79 | 0.37 | 0.79 | 1.00 |
| Unit count, head ($n = 30$) | $\Delta$ | +0.77 | +0.21 | +0.09 | +0.16 | −1.20 | −0.00 | −0.01 | −0.00 |
| | % | 57 | 53 | 50 | 67 | 40 | 50 | 57 | 53 |
| | $P$ | 1.00 | 1.00 | 1.00 | 0.81 | 0.79 | 0.76 | 0.71 | 1.00 |
| Unit count, both ($n = 30$) | $\Delta$ | −0.52 | +0.04 | +0.01 | −0.14 | +0.44 | +0.33 | +0.03 | +0.01 |
| | % | 33 | 50 | 50 | 33 | 53 | 60 | 47 | 50 |
| | $P$ | 1.00 | 1.00 | 1.00 | 0.81 | 1.00 | 0.66 | 0.79 | 1.00 |
| Stimulation† ($n = 28$) | $\Delta$ | −1.00 | −0.23 | −0.37 | −0.39 | −0.69 | −0.36 | +0.02 | +0.09 |
| | % | 36 | 46 | 46 | 36 | 46 | 46 | 43 | 46 |
| | $P$ | 0.44 | 0.66 | 0.61 | 0.11 | 0.69 | 0.13 | 0.47 | 0.52 |

Table S5: **Input-binning resolution comparison across data splits**, related to Figure 2. Median across organoids of each organoid's value (the mean of its experiments' seed-averaged values, 3 random seeds), for training, validation, and the drug-clean out-of-distribution holdout cohort. All models downsample to a uniform 20 ms output grid, so the comparison is at a common output resolution rather than at each model's native one. Bold marks the best level within each split and metric; for MAE and Poisson NLL, lower is better. $n$ is the number of organoids: 86 training, 30 validation and 20 holdout; experiments from one organoid are averaged before every statistic, and validation experiments are drawn from training organoids.

| Metric | 5 ms | 10 ms | 20 ms |
|---|---|---|---|
| **Training** | | | |
| Organoids ($n$) | 85 | 86 | 85 |
| Population Pearson $r$ | 0.7423 | 0.7489 | **0.7622** |
| Per-neuron Pearson $r$ | 0.2840 | 0.2838 | **0.2966** |
| Population bits/spike | 0.1668 | 0.1702 | **0.1720** |
| Per-neuron median bits/spike | 0.6187 | **0.6592** | 0.6559 |
| Per-neuron MAE (spikes/bin) | 0.1026 | **0.1007** | 0.1019 |
| Per-neuron Poisson NLL (nats/bin) | 0.1864 | **0.1844** | 0.1867 |
| **Validation** | | | |
| Organoids ($n$) | 30 | 30 | 30 |
| Population Pearson $r$ | **0.7381** | 0.7230 | 0.6970 |
| Per-neuron Pearson $r$ | **0.2029** | 0.2019 | 0.1965 |
| Population bits/spike | **0.2198** | 0.2167 | 0.2121 |
| Per-neuron median bits/spike | **0.3443** | 0.3392 | 0.3136 |
| Per-neuron MAE (spikes/bin) | **0.0910** | **0.0910** | 0.0915 |
| Per-neuron Poisson NLL (nats/bin) | **0.1664** | 0.1666 | 0.1680 |
| **Holdout (drug-clean)** | | | |
| Organoids ($n$) | 20 | 20 | 20 |
| Population Pearson $r$ | 0.4942 | 0.4967 | **0.4988** |
| Per-neuron Pearson $r$ | **0.0975** | 0.0957 | 0.0957 |
| Population bits/spike | 0.2006 | **0.2055** | 0.1904 |
| Per-neuron median bits/spike | -0.2755 | **-0.2568** | -0.2667 |
| Per-neuron MAE (spikes/bin) | 0.0612 | **0.0592** | 0.0603 |
| Per-neuron Poisson NLL (nats/bin) | 0.1401 | 0.1395 | **0.1394** |

Table S6: **Training-data depth sweep across data splits**, related to Figure 2. Median across organoids of each organoid's value (the mean of its experiments' seed-averaged values, 3 random seeds), for training, validation, and the drug-clean out-of-distribution holdout cohort. Levels are the cap on recordings per organoid, mapping to 266 / 566 / 780 / 1,756 training recordings. Bold marks the best level within each split and metric; for MAE and Poisson NLL, lower is better. $n$ is the number of organoids: up to 86 training, 30 validation and 20 holdout; experiments from one organoid are averaged before every statistic, and validation experiments are drawn from training organoids.

| Metric | cap1 | cap10 | cap20 | capall |
|---|---|---|---|---|
| **Training** | | | | |
| Organoids ($n$) | 80 | 85 | 86 | 86 |
| Population Pearson $r$ | 0.7405 | **0.7497** | 0.7489 | 0.7261 |
| Per-neuron Pearson $r$ | 0.2654 | 0.2816 | **0.2838** | 0.2665 |
| Population bits/spike | 0.1677 | 0.1677 | **0.1702** | 0.1598 |
| Per-neuron median bits/spike | 0.6208 | 0.6560 | **0.6592** | 0.6128 |
| Per-neuron MAE (spikes/bin) | 0.1020 | 0.1023 | **0.1007** | 0.1028 |
| Per-neuron Poisson NLL (nats/bin) | 0.1833 | **0.1813** | 0.1844 | 0.1834 |
| **Validation** | | | | |
| Organoids ($n$) | 30 | 30 | 30 | 30 |
| Population Pearson $r$ | 0.6382 | 0.7162 | 0.7230 | **0.7333** |
| Per-neuron Pearson $r$ | 0.1543 | 0.1986 | **0.2019** | 0.2012 |
| Population bits/spike | 0.1515 | 0.2008 | 0.2167 | **0.2262** |
| Per-neuron median bits/spike | 0.0533 | 0.2905 | 0.3392 | **0.3487** |
| Per-neuron MAE (spikes/bin) | 0.1015 | 0.0932 | **0.0910** | 0.0943 |
| Per-neuron Poisson NLL (nats/bin) | 0.1961 | 0.1720 | **0.1666** | 0.1700 |
| **Holdout (drug-clean)** | | | | |
| Organoids ($n$) | 20 | 20 | 20 | 20 |
| Population Pearson $r$ | 0.4931 | 0.5132 | 0.4967 | **0.5192** |
| Per-neuron Pearson $r$ | 0.0885 | 0.0946 | 0.0957 | **0.1039** |
| Population bits/spike | 0.2030 | **0.2057** | 0.2055 | 0.1834 |
| Per-neuron median bits/spike | -0.3836 | -0.3006 | -0.2568 | **-0.1318** |
| Per-neuron MAE (spikes/bin) | 0.0617 | 0.0615 | 0.0592 | **0.0577** |
| Per-neuron Poisson NLL (nats/bin) | 0.1425 | 0.1408 | 0.1395 | **0.1374** |

Table S7: **Model width scaling comparison across data splits**, related to Figure 2. Median across organoids of each organoid's value (the mean of its experiments' seed-averaged values, 3 random seeds), for training, validation, and the drug-clean out-of-distribution holdout cohort. $d_{\text{state}}$, layer count and every other hyperparameter are held fixed across widths. Bold marks the best level within each split and metric; for MAE and Poisson NLL, lower is better. $n$ is the number of organoids: 86 training, 30 validation and 20 holdout; experiments from one organoid are averaged before every statistic, and validation experiments are drawn from training organoids.

| Metric | $d128$ | $d256$ | $d512$ |
|---|---|---|---|
| **Training** | | | |
| Organoids ($n$) | 86 | 86 | 86 |
| Population Pearson $r$ | 0.6436 | 0.6826 | **0.7489** |
| Per-neuron Pearson $r$ | 0.1713 | 0.2203 | **0.2838** |
| Population bits/spike | 0.1276 | 0.1555 | **0.1702** |
| Per-neuron median bits/spike | 0.1601 | 0.3988 | **0.6592** |
| Per-neuron MAE (spikes/bin) | 0.1102 | 0.1058 | **0.1007** |
| Per-neuron Poisson NLL (nats/bin) | 0.2004 | 0.1886 | **0.1844** |
| **Validation** | | | |
| Organoids ($n$) | 30 | 30 | 30 |
| Population Pearson $r$ | 0.7235 | **0.7321** | 0.7230 |
| Per-neuron Pearson $r$ | 0.1677 | 0.1885 | **0.2019** |
| Population bits/spike | 0.2129 | **0.2178** | 0.2167 |
| Per-neuron median bits/spike | 0.1860 | 0.3062 | **0.3392** |
| Per-neuron MAE (spikes/bin) | 0.0931 | 0.0914 | **0.0910** |
| Per-neuron Poisson NLL (nats/bin) | 0.1712 | 0.1679 | **0.1666** |
| **Holdout (drug-clean)** | | | |
| Organoids ($n$) | 20 | 20 | 20 |
| Population Pearson $r$ | **0.5288** | 0.5154 | 0.4967 |
| Per-neuron Pearson $r$ | 0.1026 | **0.1067** | 0.0957 |
| Population bits/spike | **0.2152** | 0.2120 | 0.2055 |
| Per-neuron median bits/spike | **-0.1681** | -0.1942 | -0.2568 |
| Per-neuron MAE (spikes/bin) | 0.0618 | 0.0614 | **0.0592** |
| Per-neuron Poisson NLL (nats/bin) | **0.1389** | **0.1389** | 0.1395 |

Table S8: **Conditioning-feature ablation comparison**, related to Figure 2. Median across organoids of each organoid's value (the mean of its experiments' seed-averaged values, 3 random seeds). Each ablation is a separately trained checkpoint with one conditioning input removed, not a masking of the deployed model at inference. Reported on training and validation only. The stimulation-input ablation is scored on the 36 organoids with evoked-stimulation recordings and is reported in the main text rather than here, because its cohort differs from every other row of this table. $n$ is the number of organoids: 86 training and 30 validation; experiments from one organoid are averaged before every statistic, and validation experiments are drawn from training organoids.

| Condition | Population Pearson $r$ | Per-neuron Pearson $r$ | Population bits/spike | Per-neuron median bits/spike | Per-neuron MAE (spikes/bin) | Per-neuron Poisson NLL (nats/bin) |
|---|---|---|---|---|---|---|
| **Training** ($n = 86$) | | | | | | |
| All-on baseline | 0.7489 | 0.2841 | 0.1702 | 0.6592 | 0.1007 | 0.1844 |
| Sorter embedding off | 0.7457 | 0.2873 | 0.1704 | 0.6955 | 0.1004 | 0.1832 |
| Neuron count (input off) | 0.7462 | 0.2842 | 0.1693 | 0.6514 | 0.1011 | 0.1844 |
| Neuron count (head off) | 0.7490 | 0.2831 | 0.1702 | 0.6574 | 0.1008 | 0.1844 |
| Neuron count (both off) | 0.7465 | 0.2843 | 0.1697 | 0.6534 | 0.1009 | 0.1844 |
| **Validation** ($n = 30$) | | | | | | |
| All-on baseline | 0.7230 | 0.2019 | 0.2167 | 0.3392 | 0.0910 | 0.1666 |
| Sorter embedding off | 0.7235 | 0.2034 | 0.2185 | 0.3445 | 0.0903 | 0.1660 |
| Neuron count (input off) | 0.7254 | 0.2017 | 0.2156 | 0.3392 | 0.0905 | 0.1662 |
| Neuron count (head off) | 0.7259 | 0.2022 | 0.2160 | 0.3387 | 0.0900 | 0.1658 |
| Neuron count (both off) | 0.7244 | 0.2023 | 0.2158 | 0.3398 | 0.0923 | 0.1675 |

Table S9: **Paired comparisons of every Figures 2B–2E design-axis level against the reference configuration, training organoids**, related to Figure 2. For each level, three rows give the median across organoids of the paired difference (Δ, level − reference, in units of $10^{-3}$, from each organoid's mean of its experiments' seed-averaged values over 3 random seeds); the percentage of organoids (%) in which the level scored better than the reference (higher, or lower for MAE and Poisson NLL); and the Holm-corrected $P$ of a paired two-sided Wilcoxon signed-rank test. Holm correction is applied per metric within each axis (bin width, recordings per experiment, model width, and the sorter-embedding and active-unit-count ablations together). †The stimulation-input ablation is scored only on experiments with evoked-stimulation recordings and is its own uncorrected test (36 organoids). $n$ is the number of organoids: 86 training; experiments from one organoid are averaged before every statistic, and $P$ values are across organoids. Unit count, active-unit-count conditioning. Pop., population; PN, per-neuron; bits, median bits/spike for PN; MAE, mean absolute error; NLL, Poisson negative log-likelihood.

| Level | | Pop. $r$ | Pop. bits | Pop. skill | PN $r$ | PN bits | PN skill | PN MAE | PN NLL |
|---|---|---|---|---|---|---|---|---|---|
| **Conditioning removed (vs all on)** | | | | | | | | | |
| Sorter embedding ($n = 86$) | Δ | −0.15 | +0.08 | +0.49 | +5.18 | +32.19 | +8.38 | −0.46 | −0.95 |
| | % | 45 | 51 | 55 | 92 | 95 | 93 | 85 | 92 |
| | $P$ | 0.12 | 0.90 | 1.00 | $3.7\times10^{-12}$ | $1.2\times10^{-12}$ | $3.0\times10^{-12}$ | $5.5\times10^{-10}$ | $3.9\times10^{-12}$ |
| Unit count, input ($n = 86$) | Δ | −0.63 | −0.42 | −0.94 | −0.45 | −1.92 | −0.41 | +0.17 | +0.11 |
| | % | 34 | 22 | 23 | 28 | 35 | 36 | 17 | 29 |
| | $P$ | $1.8\times10^{-4}$ | $1.7\times10^{-6}$ | $6.5\times10^{-7}$ | $6.2\times10^{-5}$ | 0.02 | 0.04 | $5.4\times10^{-10}$ | $1.6\times10^{-6}$ |
| Unit count, head ($n = 86$) | Δ | +0.08 | −0.05 | −0.12 | −0.09 | +0.28 | −0.06 | +0.01 | +0.03 |
| | % | 52 | 47 | 45 | 44 | 52 | 48 | 44 | 36 |
| | $P$ | 0.48 | 0.97 | 1.00 | 0.48 | 0.52 | 1.00 | 0.14 | 0.04 |
| Unit count, both ($n = 86$) | Δ | −0.75 | −0.26 | −0.71 | −0.19 | −1.08 | −0.02 | +0.02 | +0.06 |
| | % | 24 | 23 | 24 | 38 | 43 | 49 | 37 | 34 |
| | $P$ | $4.8\times10^{-5}$ | $9.5\times10^{-5}$ | $5.7\times10^{-5}$ | 0.003 | 0.03 | 1.00 | 0.003 | 0.001 |
| Stimulation† ($n = 36$) | Δ | −3.26 | −4.03 | −6.43 | −1.44 | −8.22 | −2.06 | +0.19 | +0.47 |
| | % | 6 | 3 | 0 | 11 | 17 | 14 | 25 | 8 |
| | $P$ | $6.0\times10^{-9}$ | $7.3\times10^{-10}$ | $2.9\times10^{-11}$ | $7.4\times10^{-9}$ | $3.2\times10^{-6}$ | $1.1\times10^{-7}$ | $6.6\times10^{-5}$ | $1.3\times10^{-9}$ |

Table S10: **Genome-wide normalized expression of the bulk RNA-seq organoid samples**, related to Figure 3. Provided as a separate Excel file. Counts per million (CPM) for every Ensembl gene detected in the 14 samples of Figures 3B and S4 (each a pool of three organoids), with a Metadata tab giving each sample's line, genotype, protocol, age at collection, sequencing run and library size.

Table S11: **Prediction accuracy of held-out mouse cultures and the two human control lines, zero-shot and after a 2-minute fine-tuned linear head**, related to Figure 3. Tests are run across cultures (the 20 held-out mouse experiments come from 9 cultures; 17 of the experiments are on 2D monolayers). Each culture is summarised by the mean of its recordings' seed-averaged values (3 model seeds), and group columns give the median across cultures of that summary; column headings give the number of cultures. $P$ values are two-sided Mann–Whitney $U$ tests across cultures with an exact permutation null, one test per metric and stage, with metrics treated as separate comparisons. The two mouse–control comparisons form one Holm family per metric and stage ($P_{\mathrm{Holm}}$); Cohen's $d$ (on culture means) is mouse − control, so $d > 0$ means the human control line is harder to predict. The control–control comparison is a single test per metric and stage (uncorrected); $d$ is CW20107 − *SHANK3*$^{+/+}$, so $d > 0$ means *SHANK3*$^{+/+}$ is harder to predict. The smallest attainable $P$ is 0.0004 for 9 versus 6 cultures and 0.0022 for 6 versus 6. The two control lines differ in preparation protocol, recording dates, age range and unit yield, so this comparison measures a difference between preparations, not between control genotypes.

| | Median | | | Mouse vs CW20107 | | Mouse vs *SHANK3*$^{+/+}$ | | CW20107 vs *SHANK3*$^{+/+}$ | |
|---|---|---|---|---|---|---|---|---|---|
| Metric | Mouse ($n = 9$) | CW20107 ($n = 6$) | *SHANK3*$^{+/+}$ ($n = 6$) | $d$ | $P_{\mathrm{Holm}}$ | $d$ | $P_{\mathrm{Holm}}$ | $d$ | $P$ |
| **Zero-shot** | | | | | | | | | |
| Population Pearson $r$ | 0.466 | 0.663 | 0.545 | -1.33 | 0.051 | -0.75 | 0.224 | 0.71 | 0.240 |
| Population bits/spike | 0.040 | 0.394 | 0.244 | -1.15 | 0.072 | -0.42 | 0.272 | 1.07 | 0.065 |
| Population skill | 0.037 | 0.322 | 0.229 | -1.34 | 0.051 | -0.93 | 0.145 | 0.52 | 0.394 |
| Per-neuron Pearson $r$ | 0.104 | 0.216 | 0.124 | -2.01 | 0.0096 | -0.34 | 0.607 | 2.28 | 0.0087 |
| Per-neuron bits/spike | -0.335 | 0.611 | -0.046 | -2.66 | 0.0032 | -1.01 | 0.066 | 2.21 | 0.0087 |
| Per-neuron skill | -0.069 | 0.110 | -0.011 | -2.65 | 0.0032 | -1.22 | 0.018 | 2.19 | 0.0087 |
| +2 **min fine-tuned head** | | | | | | | | | |
| Population Pearson $r$ | 0.597 | 0.713 | 0.606 | -0.95 | 0.176 | -0.23 | 0.776 | 0.68 | 0.180 |
| Population bits/spike | 0.210 | 0.539 | 0.240 | -0.87 | 0.227 | 0.13 | 0.776 | 1.36 | 0.041 |
| Population skill | 0.302 | 0.449 | 0.275 | -0.68 | 0.544 | -0.02 | 0.864 | 0.58 | 0.180 |
| Per-neuron Pearson $r$ | 0.212 | 0.302 | 0.180 | -0.86 | 0.544 | 0.59 | 0.544 | 1.86 | 0.041 |
| Per-neuron bits/spike | 0.359 | 1.289 | 0.420 | -1.18 | 0.099 | 0.40 | 0.689 | 2.66 | 0.0087 |
| Per-neuron skill | 0.074 | 0.237 | 0.079 | -1.16 | 0.176 | 0.30 | 0.864 | 2.09 | 0.0087 |

Table S12: **Prediction accuracy by human line, zero-shot and after a 2-minute fine-tuned linear head, with the causal EWMA baseline**, related to Figure 3. Companion to Figures 3C and S5; every $d$ and $p$ here is the value behind those figures' significance brackets. The causal EWMA block gives the baseline drawn as orange dashes in both figures; it is not tested between lines, so its $d$ and $p$ cells are empty. Tests are run across organoids. Each organoid is summarised by the mean of its recordings' seed-averaged values (3 model seeds), and line columns give the median across organoids of that summary. Column headings give the number of organoids per line: 6 versus 5 (*SHANK3*$^{+/+}$ versus *SHANK3*$^{+/-}$) and 6 versus 6 (CW20107 versus 14765-X2). Cohen's $d$ is computed on organoid means and is unaffected − affected, so $d > 0$ means the affected line is harder to predict; $p$ is a two-sided Mann–Whitney $U$ test across organoids with an exact permutation null (all relabelings of the organoids), one test per metric, with each metric treated as a separate comparison and not corrected across metrics. The smallest attainable $p$ is 0.0043 for 6 versus 5 organoids and 0.0022 for 6 versus 6. The two comparisons come from independent preparations, and are reported side by side but never pooled or contrasted with each other; no $p$ in this table compares one comparison's control line against the other's.

| | SHANK3 arm (6 vs 5) | | | | 14765-X2 arm (6 vs 6) | | | |
|---|---|---|---|---|---|---|---|---|
| Metric | *SHANK3*$^{+/+}$ | *SHANK3*$^{+/-}$ | $d$ | $p$ | CW20107 | 14765-X2 | $d$ | $p$ |
| **Zero-shot** | | | | | | | | |
| Population Pearson $r$ | 0.545 | 0.319 | 0.71 | 0.247 | 0.663 | 0.623 | 0.52 | 0.394 |
| Population bits/spike | 0.244 | -0.001 | 0.17 | 0.247 | 0.394 | 0.230 | 1.11 | 0.132 |
| Population skill | 0.229 | 0.009 | 0.50 | 0.329 | 0.322 | 0.310 | 0.44 | 0.589 |
| Per-neuron Pearson $r$ | 0.124 | 0.089 | 0.24 | 0.537 | 0.216 | 0.124 | 2.47 | 0.0043 |
| Per-neuron bits/spike | -0.046 | -0.180 | -0.27 | 0.792 | 0.611 | -0.059 | 2.23 | 0.0043 |
| Per-neuron skill | -0.011 | -0.031 | -0.25 | 0.662 | 0.110 | -0.012 | 2.19 | 0.0087 |
| **+2 min fine-tuned head** | | | | | | | | |
| Population Pearson $r$ | 0.606 | 0.416 | 0.72 | 0.247 | 0.713 | 0.694 | 0.32 | 0.485 |
| Population bits/spike | 0.240 | 0.138 | 0.07 | 0.329 | 0.539 | 0.335 | 1.18 | 0.132 |
| Population skill | 0.275 | 0.148 | 0.41 | 0.247 | 0.449 | 0.430 | 0.31 | 0.699 |
| Per-neuron Pearson $r$ | 0.180 | 0.134 | 0.49 | 0.429 | 0.302 | 0.219 | 2.39 | 0.0043 |
| Per-neuron bits/spike | 0.420 | 0.287 | -0.26 | 0.792 | 1.289 | 0.645 | 2.18 | 0.0087 |
| Per-neuron skill | 0.079 | 0.054 | -0.05 | 0.792 | 0.237 | 0.125 | 2.01 | 0.0087 |
| **Causal EWMA baseline** | | | | | | | | |
| Population Pearson $r$ | 0.325 | 0.256 | – | – | 0.283 | 0.202 | – | – |
| Population bits/spike | 0.138 | 0.074 | – | – | 0.126 | 0.049 | – | – |
| Population skill | 0.114 | 0.070 | – | – | 0.100 | 0.058 | – | – |
| Per-neuron Pearson $r$ | 0.084 | 0.083 | – | – | 0.127 | 0.077 | – | – |
| Per-neuron bits/spike | 0.164 | 0.097 | – | – | 0.139 | 0.079 | – | – |
| Per-neuron skill | 0.028 | 0.018 | – | – | 0.027 | 0.014 | – | – |

Table S13: **Summary statistics for the NPD network phenotype**, related to Figure 4. Every comparison is one NPD line vs its own control within its own preparation (CWSF: CW20107 vs 14765-X2; KOLF: $SHANK3^{+/+}$ vs $SHANK3^{+/-}$); nothing here pools recordings across the two. Tests are run across organoids; recordings of the same organoid are repeated measures. Every $P$ is a permutation $P$ with genotype permuted across organoids within preparation (exact enumeration), except the clock-ordering and unit-count-control $P$ in block C (age shuffled within organoid, 20,000 draws). $n$ is organoids. All panels use corrected preprocessing (scaler off; see Methods).

**A. Convergent latent displacement across two independent NPD models (Figure S6D)**

| Quantity | Statistic | Perm. $P$ |
|---|---|---|
| Latent displacement cosine (within-prep, primary) | +0.464 (62.3°) | 0.0019 |
| organoid bootstrap 95% CI [−0.006, +0.500] (exact null, 426,888 relabelings) | | |
| Feature-profile agreement (28-d $\delta$-profile), raw | +0.063 | 0.445 |
| Feature-profile agreement (28-d $\delta$-profile), adjusted | +0.709 | 0.013 |
| same-sign features: 15/28 raw, 20/28 adjusted (exact null, 426,888 relabelings) | | |
| PC2's share of the adjusted-displacement cosine: 29% | | |
| PC1/PC2 correlation with sorted unit count (human, $n = 23$ organoids, organoid means): $\rho = -0.72$ / $\rho = +0.76$ | | |
| *Robustness (cosine, perm. $P$)* | | |
| Raw (unadjusted) | −0.236 | 0.789 |
| + age covariate | +0.474 | 0.0014 |
| Control-only slope (refit per permutation) | +0.584 | $5.7 \times 10^{-4}$ |
| Drop PC1 (full latent space) | +0.078 | 0.361 |
| *Same-sign counts by feature family (raw / adjusted, of n)* | | |
| bursting ($n = 5$) | 5/5 | 5/5 |
| criticality ($n = 5$) | 2/5 | 3/5 |
| network topology ($n = 5$) | 5/5 | 5/5 |
| rate/yield ($n = 8$) | 1/8 | 2/8 |
| temporal/info ($n = 5$) | 2/5 | 5/5 |
| *Leave-one-arm-out (feature-profile Cliff's $\delta$, perm. $P$ across organoids of the tested arm)* | | |
| Direction from 14765-X2, tested on SHANK3 | +0.429 | 0.227 |
| Direction from SHANK3, tested on 14765-X2 (CWSF) | +0.176 | 0.422 |

**B. Unit-adjusted PC2 displacement vs own control (Figure 4E)**

| Comparison | $n_{\mathrm{ctrl}}$ | $n_{\mathrm{NPD}}$ | Cliff's $\delta$ (unit-adjusted) | Perm. $P$ |
|---|---|---|---|---|
| 14765-X2 vs CW20107 (own prep) | 6 | 6 | +0.411 | 0.037 (*) |
| $SHANK3^{+/-}$ vs $SHANK3^{+/+}$ (own prep) | 6 | 5 | +0.429 | 0.262 (n.s.) |
| **Joint (both preparations)** | | | +0.420 | 0.022 |
| *Raw (unadjusted), for reference* | | | | |
| 14765-X2 vs CW20107 (own prep) | | | +0.635 | 0.0087 |
| $SHANK3^{+/-}$ vs $SHANK3^{+/+}$ (own prep) | | | +0.253 | 0.444 |
| Joint (both preparations) | | | +0.444 | 0.021 |

Table S13: **Summary statistics for the NPD network phenotype (continued)**, related to Figure 4.

**C. Developmental clock, organoid level (Figure 4A–C)**

| Quantity | $n_{\text{ctrl}}$ | $n_{\text{NPD}}$ | Statistic | Perm. $P$ |
|---|---|---|---|---|
| *Clock ordering within organoid: per-organoid Spearman $\rho$ of predicted on actual age, mean over organoids (median; organoids above zero); $P$ with age shuffled within organoid* | | | | |
| CW20107, leave-one-organoid-out fit | 6 | | +0.46 (+0.64; 5/6) | 0.0015 |
| Sorted-unit-count control (same protocol) | 6 | | +0.12 (+0.15; 4/6) | 0.226 |
| 14765-X2 (projection, no refit) | 6 | | +0.70 (+0.74; 6/6) | $1.0 \times 10^{-4}$ |
| SHANK3$^{+/+}$ (projection; $\rho$ of organoid means) | 6 | | +0.543 | 0.149 |
| SHANK3$^{+/-}$ (projection; $\rho$ of organoid means) | | 5 | −0.100 | 0.608 |
| *Clock offset, Cliff's $\delta$ (age shuffled within organoid; genotype permuted across organoids)* | | | | |
| 14765-X2 vs CW20107 | 6 | 6 | +0.449 | 0.0022 (**) |
| SHANK3$^{+/-}$ vs SHANK3$^{+/+}$ | 6 | 5 | −0.022 | 0.950 (n.s.) |

Table S14: **Electrode decoding errors point back to the previous electrode, as expected mainly of a memory of earlier pulses rather than a clock**, related to Figure 5. In the causal sweep each electrode is stimulated in one contiguous stretch of 50 pulses, so electrode identity also marks a time within the recording. A memory of earlier pulses can only bias predictions toward the preceding electrode, at the start of a stretch; a clock or slow drift would bias them toward the following electrode at its end as well. A decoder trained on pulses 11 to 50 of each electrode predicted held-out pulses. Pull, the fraction of predictions equal to the neighboring electrode, as a multiple of chance (median over organoids): toward the preceding electrode for pulses 1 to 5, and toward the following electrode for pulses 46 to 50. The untrained model has the same architecture with random weights. Each organoid's pull is the mean over its recordings. Preceding > following counts organoids, and $P$ is a paired two-sided Wilcoxon signed-rank test on per-organoid means of the two pulls ($n = 17$ organoids).

| Feature | Preceding pull | Following pull | Preceding > following | $P$ |
|---|---|---|---|---|
| Model latent | 1.98 | 1.20 | 14/17 | 0.0013 |
| Firing rate | 0.95 | 1.14 | 6/17 | 0.40 |
| Untrained model latent | 1.19 | 0.97 | 13/17 | 0.03 |

Table S15: **Electrode-specific structure changes more slowly than the global drift**, related to Figure 5. Shared drift: fraction of each held-out cycle's drift variance common to all electrodes, per organoid. Electrode decoding: LDA decoder fit on one cycle and tested on another, as a multiple of chance; per organoid, the mean over its experiments of the median over all cycle pairs at that separation. The uncorrected readout is the evoked latent state as recorded; the per-pulse readout subtracts each response's own pre-stimulus baseline. Far / within, each organoid's accuracy at 7–12 cycles apart divided by its within-cycle accuracy; one organoid never reached that separation. $P$, paired two-sided Wilcoxon signed-rank test of far / within between the readouts, run on per-organoid means (8 organoids for the shared-drift and within-cycle values, 7 for the far-separation and decoder-decay comparison). Separation, wall-clock time between the two recordings of each cycle pair. Values are median [IQR] across organoids.

| | | | |
|---|---|---|---|
| **Drift shared across all electrodes ($n = 8$ organoids)** | | | |
| Uncorrected readout | 76.2% [72.8–80.2%] | – | – |
| Per-pulse readout | 8.6% [7.7–9.5%] | – | – |
| **Electrode decoding (× chance)** | | | |
| | Within cycle ($n = 8$ organoids) | 7–12 cycles apart ($n = 7$ organoids) | Far / within |
| Per-pulse readout | 3.43 [1.96–3.61] | 1.86 [1.47–2.06] | 0.53 [0.52–0.68] |
| Uncorrected readout | 12.55 [10.95–13.42] | 2.29 [1.75–3.28] | 0.21 [0.16–0.28] |
| Uncorrected declined faster than per-pulse | 7/7 organoids, $P = 0.016$ | | |
| Separation at 7–12 cycles | median 32.1 h, maximum 63.3 h | | |

Table S16: **Stimulation generates a subharmonic yet stays inside the spontaneous subspace**, related to Figure 6. Subharmonic: $f/2$ power relative to the larger of two off-harmonic control frequencies (3.0 and 5.5 Hz) in the same recording, on the model's 20 ms latent grid; median [IQR] across organoids, each organoid's value the mean of its experiments' median ratios (geometric mean); $P$, two-sided Wilcoxon signed-rank test against 1. Surrogate rows replace the recorded spikes with firing-rate surrogates carrying only the stimulation frequency, or the stimulation frequency and its harmonics, and are scored with the same ratio. Subspace overlap: fraction of one recording's variance inside the leading principal components of a reference recording, retaining 90% of the reference's variance ($k$ = 23–122); the every-$k$ row gives the range of cohort medians over all sampled subspace sizes and the lowest single organoid at any size. Chance, a randomly oriented subspace of the same size ($k/512$). Ceiling, a spontaneous recording scored against its preceding spontaneous recording. $P$ for stimulation in preceding spontaneous, one-sided Wilcoxon signed-rank test against chance; for the first spontaneous recording, two-sided paired Wilcoxon signed-rank test against the preceding one. Firing-rate rows repeat the procedure on binned firing rates. Spontaneous subspace decay: fixed 50-dimensional subspace, with each recording's split-half self-capture as the ceiling and 50/512 as chance. Same-cycle and most distant, the closest and farthest separation bins of spontaneous-recording pairs. Paired change, difference in captured variance between those bins in percentage points (pp); $P$, two-sided Wilcoxon signed-rank test. Values are medians across the 8 organoids, each organoid's value being the mean of its experiments' values; $n$ above counts organoids, and every $P$ is the test run on those per-organoid values, so the smallest attainable exact $P$ is 0.0078 (two-sided) or 0.0039 (one-sided).

| Quantity | Median [IQR] / value | $n$ above | $P$ |
|---|---|---|---|
| **Subharmonic ($f/2$ relative to off-harmonic controls)** | | | |
| 4 Hz stimulation | 0.93 [0.87–1.00] | 2/8 | – |
| 8 Hz stimulation | 3.17 [1.21–37.92] | 7/8 | 0.016 |
| 8 Hz, surrogate with stimulation frequency only | 0.90 [0.85–0.92] | – | – |
| 8 Hz, surrogate with stimulation frequency and harmonics | 0.93 [0.88–0.95] | – | – |
| **Subspace overlap ($k$ retaining 90% of reference variance)** | | | |
| Stimulation in preceding spontaneous | 86.7 [86.6–87.0]% | 8/8 | 0.0039 |
| Spontaneous-on-spontaneous ceiling | 88.3 [87.8–88.6]% | – | – |
| Stimulation as % of ceiling | 98.5 [97.7–98.7]% | – | – |
| Chance (randomly oriented subspace) | 13.1% | – | – |
| Stimulation as % of spontaneous, every $k$ from 1 to 256 | 96.8–99.6% (lowest organoid 91.7%) | – | – |
| Frequency range: 0.5 Hz → 8 Hz | 87.7 → 85.1% | – | – |
| Firing rate: stimulation as % of firing-rate ceiling | 99.7 [99.2–99.9]% | – | – |
| Firing rate: chance | 46.4% | – | – |
| Stimulation onto first (distant) spontaneous recording | 66.7 [64.3–67.7]% | – | 0.0078 |
| Paired drop, adjacent → distant | 19.5 pp | – | – |
| Spontaneous onto first spontaneous recording (distant ceiling) | 68.5 [65.1–69.2]% | – | – |
| Stimulation as % of distant ceiling | 98.2 [97.8–99.5]% | – | – |
| **Spontaneous subspace decay (fixed 50-dimensional subspace)** | | | |
| Split-half ceiling | 85.7% | – | – |
| Chance (50/512) | 9.8% | – | – |
| Same-cycle, % of ceiling | 97.0 [94.0–102.9]% | – | – |
| Most distant, % of ceiling | 64.2 [59.0–73.9]% | – | – |
| Paired change, same-cycle → most distant | −27.6 pp | 8/8 | 0.0078 |

Table S17: **Three controls rule out a binning alias, a shared-encoder artifact and an episode-time confound**, related to Figures 6 and 7. Subharmonic: this check is on recorded firing rates re-binned at 20 ms versus 25 ms (no model latents), median across organoids of $f/2$ power in stimulation recordings divided by median $f/2$ power in spontaneous recordings at the same bin width (a stimulation-versus-spontaneous ratio, distinct from the off-harmonic-floor ratio behind the main-figure 0.930/3.17 values), at 8 Hz stimulation on a 25 ms bin grid, on which 8 Hz falls at exactly 5 bins per cycle so no bin-alternation alias is possible; compared with the 4 Hz control frequency and with the paper's 20 ms grid. Cross-organoid subspace null: median fraction of stimulation-evoked variance inside the same organoid's own preceding spontaneous subspace versus inside other organoids' spontaneous subspaces, against the chance floor for a randomly oriented subspace of the same size. Episode-time-matched balance decoder: median matched-AUC for the balance-versus-fast-fail latent decoder (Figure S14A) reproduced on the well-powered subset, before and after adding time since episode start as an additional coarsened-exact-matching covariate. $n$ above counts organoids above the comparison in the subharmonic and cross-organoid rows, and organoids in the decoder row (for the decoder, significant with and without the covariate); two-sided Wilcoxon signed-rank tests throughout, run on per-organoid means (subharmonic and cross-organoid rows: $n = 8$ organoids; decoder: $n = 7$ organoids).

| Control | Comparison | Value | | $n$ above | $P$ |
|---|---|---|---|---|---|
| **Subharmonic, alias-free 25 ms grid** | $f/2$ at 8 Hz vs. spontaneous null | 2.46× | | 8/8 | – |
| | 8 Hz vs. 4 Hz (25 ms grid) | – | | 8/8 | 0.0078 |
| | 20 ms grid vs. 25 ms grid (no alias difference) | – | | – | 0.15 |
| **Cross-organoid subspace null** | Own vs. other organoids' spontaneous subspace (chance ≈ 0.14) | 0.867 0.162 | vs. | 8/8 | 0.0078 |
| **Episode-time-matched balance decoder** | Matched AUC without vs. with episode time as a covariate | 0.794 0.785 | vs. | 6/7 | 0.22 |

Table S18: **Closed-loop control moves the network's average state further than open-loop stimulation**, related to Figure 7. Statistics behind Figure 7C and Figure S13A, B and F. Location and orientation: one-sample Wilcoxon signed-rank tests of $\log_2$ ratios against one, and paired tests between regime pairs (> rows; one-sided for the two contrasts with the spontaneous–closed-loop shift, two-sided for the third), each family Holm-corrected across its three tests. Drift: paired Wilcoxon signed-rank test of drift against each organoid's own shift at each gap, Holm-corrected across gaps; one closed-loop recording ran per hour. Every value is the mean over an organoid's experiments (ratios averaged on the log scale), $n$ is organoids ($n = 13$ unless a gap is reached by fewer; gaps reached by at least five organoids), and every $P$ comes from the same test run on those organoid values. Median [interquartile range], across organoids.

| Measure | Value | $n$ | $P$ |
|---|---|---|---|
| *Location shift, multiple of within-recording movement (Figure 7C)* | | | |
| Spontaneous–closed-loop, median | 15.5× | 13 | – |
| Spontaneous–open-loop, median | 4.4× | 13 | – |
| Open-loop–closed-loop, median | 9.9× | 13 | – |
| Each pair above 1× | 13/13 each | 13 | $3.7 \times 10^{-4}$ each |
| Spontaneous–closed-loop > spontaneous–open-loop | 13/13 | 13 | $3.7 \times 10^{-4}$ |
| Spontaneous–open-loop > open-loop–closed-loop | 2/13 | 13 | 0.012 |
| Spontaneous–closed-loop > open-loop–closed-loop | 11/13 | 13 | 0.047 |
| *Orientation change, same units (Figure S13B)* | | | |
| Spontaneous–closed-loop, median | 4.6× | 13 | – |
| Spontaneous–open-loop, median | 1.9× | 13 | – |
| Open-loop–closed-loop, median | 3.4× | 13 | – |
| Each pair above 1× | 13/13 each | 13 | $3.7 \times 10^{-4}$ each |
| Spontaneous–closed-loop > spontaneous–open-loop | 13/13 | 13 | $3.7 \times 10^{-4}$ |
| Spontaneous–open-loop > open-loop–closed-loop | 2/13 | 13 | 0.012 |
| Spontaneous–closed-loop > open-loop–closed-loop | 11/13 | 13 | 0.012 |
| *Location shift kept after residualizing firing rate (Figure S13A)* | | | |
| Spontaneous–closed-loop, median | 98.6% | 13 | – |
| Spontaneous–open-loop, median | 99.6% | 13 | – |
| Open-loop–closed-loop, median | 99.8% | 13 | – |
| *Closed-loop drift, fraction of the spontaneous–closed-loop shift (Figure S13F)* | | | |
| 1 h apart, median [IQR] | 0.10 [0.07–0.24] | 13 | 0.0046 |
| 9 h apart, median [IQR] | 0.33 [0.28–0.66] | 8 | 0.1 |
| Every gap from 1 to 5 h | below 1 | 11–13 | $\leq 0.026$ |
| Ever reached the shift, organoids (median time) | 4/13 (13.5 h) | 13 | – |

Table S19: **Closed-loop and open-loop activity fall inside the spontaneous subspace in model latent space**, related to Figure 7. Median fraction of one regime's variance captured by the leading principal components of another regime's activity, with interquartile range (IQR) and chance floor (the fraction expected of a randomly oriented subspace of the same size). Subspace size is the number of principal components holding 90% of the reference regime's variance (median $k = 70$ of 512 dimensions). Each experiment had a single spontaneous recording, so the ceiling splits it at its midpoint and projects the second half onto the first half's subspace; the percentage of ceiling is the median across organoids of the ratio. $n = 13$ organoids. Above chance counts organoids; each comparison exceeds chance in every organoid (one-sided Wilcoxon signed-rank test, $P = 1.2 \times 10^{-4}$ for each, the smallest exact value for 13 organoids). Each value is the mean over an organoid's experiments; medians and IQRs are across organoids.

| Comparison | Median (%) | IQR (%) | Chance (%) | Above chance |
|---|---|---|---|---|
| Open-loop in spontaneous | 81.6 | 80.7–85.3 | 13.8 | 13/13 |
| Closed-loop in spontaneous | 75.4 | 72.7–76.9 | 13.8 | 13/13 |
| Closed-loop in open-loop | 80.4 | 78.0–82.2 | 13.6 | 13/13 |
| Spontaneous second half in first half (ceiling) | 87.9 | 87.5–88.8 | 12.9 | 13/13 |
| *As % of ceiling* | | | | |
| Open-loop in spontaneous | 95.0 | 92.1–96.0 | – | – |
| Closed-loop in spontaneous | 85.7 | 81.8–88.2 | – | – |

Table S20: **Closed-loop activity and the direction separating balance from failure reuse spontaneous directions**, related to Figure 7. Statistics behind Figure 7F and G and Figures S13C and D and S14; subspace overlap medians are in Tables S19 and S21, and balance-direction nulls in Table S22. Axis alignment: null from 20,000 permutations of stimulation-condition identity, matched for subspace dimensionality. Decoders: significance from a one-sided episode-level permutation test (1,000 draws); latent against firing rate by two-sided paired Wilcoxon signed-rank test. Other tests are one-sided Wilcoxon signed-rank tests against chance. Every value is the mean over an organoid's experiments and $n$ is organoids ($n = 13$ unless a subset is smaller); every $P$ comes from a test on those organoid values (the axis-alignment null exchanges whole organoids; decoder AUCs are tested against 0.5 by one-sided Wilcoxon signed-rank test, with an organoid counted as significant when the Fisher-combined permutation $P$ of its experiments is below 0.05). Median [interquartile range], across organoids.

| Measure | Value | $n$ | $P$ |
|---|---|---|---|
| *Variance captured by spontaneous PCs (Figure 7F, 7G)* | | | |
| Open-loop, leading 10 PCs, median | 56.0% | 13 | – |
| Closed-loop, leading 10 PCs, median | 44.7% | 13 | – |
| Each pair above chance (Figure 7G) | 13/13 each | 13 | $1.2 \times 10^{-4}$ each |
| *Closed-loop axis versus open-loop axis (Figure S13C, S13D)* | | | |
| Mean $\lvert\cos\rvert$ across organoids, observed vs null | 0.460 vs 0.227 | 13 | $5.0 \times 10^{-5}$ |
| Same, firing-rate axis removed | 0.414 vs 0.212 | 13 | $5.0 \times 10^{-5}$ |
| Drop on removing that axis, mean (organoids dropped) | 0.045 (12/13) | 13 | $2.4 \times 10^{-4}$ |
| Split-half cosine, median [IQR] | 0.979 [0.930–0.992] | 13 | – |
| *Balance versus failure, matched decoder AUC (Figure S14A, S14B)* | | | |
| Model latent, median [IQR] (significant) | 0.790 [0.705–0.862] (10/12) | 12 | $2.4 \times 10^{-4}$ |
| Model latent, $\geq 25$ balance episodes (significant) | 0.832 [0.772–0.846] (6/6) | 6 | each $\leq 0.001$ |
| Firing rate, median [IQR] | 0.703 [0.608–0.765] | 12 | 0.0012 |
| Latent minus firing rate, paired (latent higher) | +0.087 (12/12) | 12 | $4.9 \times 10^{-4}$ |
| Model latent, unmatched, median [IQR] | 0.850 [0.780–0.887] | 12 | – |
| Plus time in recording as matching key (significant) | 0.817 to 0.793 (6/7) | 7 | – |
| Episodes, topology analysis (Figure S14C, S14D) | 1,387 | 3 | – |
| *Balance direction captured by spontaneous PCs (Figure S14E)* | | | |
| Model latent vs random direction (above) | 62.7 [56.6–67.6]% vs 11.2% (10/10) | 10 | $9.8 \times 10^{-4}$ |
| Firing rate at half the dimensions (above) | 93.1% vs 50% (10/10) | 10 | $9.8 \times 10^{-4}$ |

Table S21: **Closed-loop and open-loop activity fall inside the spontaneous subspace in binned firing rates**, related to Figure 7. Median fraction of one regime's variance captured by the leading principal components of another regime's activity, with interquartile range (IQR) and chance floor (the fraction expected of a randomly oriented subspace of the same size). Computed identically but on per-neuron binned firing rates instead of model latents. The chance floor is higher (∼50% vs. ∼14%) because the ambient dimensionality (median 134 sorted units per organoid) is lower than the 512-dimensional latent. $n = 10$ organoids. Above chance counts organoids; each comparison exceeds chance in every organoid (one-sided Wilcoxon signed-rank test, $P = 9.8 \times 10^{-4}$ for each, the smallest exact value for 10 organoids). Each value is the mean over an organoid's experiments; medians and IQRs are across organoids.

| Comparison | Median (%) | IQR (%) | Chance (%) | Above chance |
|---|---|---|---|---|
| Open-loop in spontaneous | 88.4 | 86.7–89.5 | 48.5 | 10/10 |
| Closed-loop in spontaneous | 85.9 | 84.2–88.5 | 48.5 | 10/10 |
| Closed-loop in open-loop | 87.9 | 87.0–88.1 | 47.2 | 10/10 |

Table S22: **The balance-versus-failure direction is a mode of spontaneous activity but is not singled out among closed-loop directions**, related to Figure 7. The direction is the difference between mean sustained-balance and mean quick-failure states. Scrambled floor, spontaneous variance along the direction after each dimension of spontaneous activity is shifted independently in time, which keeps each dimension's variance and removes their covariance. Shuffled-label axes, the same direction rebuilt from randomly relabeled episodes (300 draws); a percentile of 50 means the real direction is as aligned with spontaneous activity as a typical closed-loop direction. Compare within a column only: the two spaces differ in dimensionality. $n = 10$ organoids; each value is the mean over an organoid's experiments (ratios averaged on the log scale), medians and ranges are across organoids, and each $P$ is a Wilcoxon signed-rank test on the same organoid values.

| Measure | Model latent | Firing rate |
|---|---|---|
| Spontaneous variance along the axis, ratio to scrambled floor, median (range) | 9.7 (1.8–16.8) | 2.6 (0.9–5.2) |
| Organoids above 2× the scrambled floor | 9/10 | 7/10 |
| $P$, ratio $> 1$ (one-sided Wilcoxon) | $9.8 \times 10^{-4}$ | 0.0049 |
| Percentile among shuffled-label axes, median (IQR) | 26 (18–50) | 49 (34–59) |
| $P$, percentile $\neq 50$ (two-sided Wilcoxon) | 0.035 | 0.75 |
